# Benchmarking strategies for cross-species integration of single-cell RNA sequencing data

**DOI:** 10.1101/2022.09.27.509674

**Authors:** Yuyao Song, Zhichao Miao, Alvis Brazma, Irene Papatheodorou

**Affiliations:** European Molecular Biology Laboratory-European Bioinformatics Institute (EMBL-EBI), Wellcome Genome Campus, Hinxton, United Kingdom

## Abstract

The growing number of available single cell gene expression datasets from different species creates opportunities to explore evolutionary relationships between cell types across species. Cross-species integration of single-cell RNA-sequencing data has been particularly informative in this context. However, in order to do so robustly it is essential to have rigorous benchmarking and appropriate guidelines to ensure that integration results truly reflect biology. We benchmarked 28 combinations of gene homology mapping methods and data integration algorithms in a variety of biological settings. We examined the capability of each strategy to perform species-mixing of known homologous cell types and to preserve biological heterogeneity using 9 established metrics. We also developed a new biology conservation metric to address the maintenance of cell type distinguishability. Overall, scANVI, scVI and SeuratV4 methods achieved a balance between species-mixing and biology conservation. For evolutionarily distant species, including in-paralogs was beneficial. SAMap outperformed when integrating whole-body atlases between species with challenging gene homology annotation. We provided our freely available cross-species integration and assessment pipeline to help analyse new data and develop new algorithms.

## Introduction

Animal cells show conspicuous diversity and have fundamental unities across species. Recently, single-cell RNA-sequencing (scRNA-seq) and single-nucleus RNA-sequencing (snRNA-seq) have been applied in a diversity of species, generating full-body cell atlases from transcriptome profiles of millions of individual cells ^1–7^. Comparing cellular expression profiles cross-species provides insights into the origin and evolution of organs and cell types ^6, 8–10^, highlights species-specific expression patterns ^11, 12^, as well as further delineates how cell types execute their functions in health and disease 13,14.

Due to millions of years of evolution, the gene expression profiles of evolutionarily related cell types from different species exhibit significant global transcriptional shifts ^11, 15, 16^. Mapping genes based on homology is the first step to put cells from different species into the same expression space for comparison. After achieving such mapping, in a joint analysis of scRNA-seq/snRNA-seq data on the raw expression space a “batch effect” arises because cells from the same species usually show higher degree of transcriptomic similarity, instead of with their cross-species counterparts ^16, 17^. We term this “species effect” to distinguish from pure technical batch effects between same-species samples ^18^. By correcting for species effect during joint analysis of scRNA-seq data, we can find a latent representation of cells from different species on which they form clusters based on transcriptomic identity. The goal is to keep the cells of homologous type mixed regardless of the species, while cells of different types are separated.

Various data integration methods originally designed for batch correction or cross-condition integration have been rapidly adapted to cross-species analysis ^16^. A recent benchmark of batch correction methods ^17^ highlighted top-performing algorithms including: mutual nearest neighbours based fastMNN ^19^; iterative clustering based Harmony ^20^; LIGER which utilises integrative non-negative matrix factorization (iNMF) ^21^ and its recent upgrade LIGER UINMF that also takes unshared features ^22^; a panorama-stitching algorithm Scanorama ^23^; a probabilistic model with distributions specified by deep neural networks scVI ^24^ and its semi-supervised extension scANVI ^25^; as well as SeuratV4 methods, which uses canonical correlation analysis (CCA) or reciprocal principal component analysis (RPCA) to identify anchors between datasets then uses dynamic time warping to align the subspaces ^26–28^.

Although a variety of batch-correction algorithms are available, it is still very challenging to generate an informative cross-species joint embedding ^16, 17, 29^. On one hand, species effects can be much stronger than average technical batch effects, so moderate methods frequently fail to integrate the data ^17, 29^. On the other hand, species-specific populations may become obscure due to overfitting ^16^. Importantly, mapping genes from different species by gene homology can cause significant information loss since homology annotation can be challenging for species without a well-annotated genome and between evolutionally distant species ^30^. The method SAMap tackles this issue by reciprocally and iteratively updating a gene-gene mapping graph from *de-novo* basic local alignment search tool (BLAST) analysis and a cell-cell mapping graph to stitch atlases between species ^29^. SAMap can lay much stronger mapping of homologous cell types across distant species and is capable of discovering gene paralog substitution events. However, SAMap is computationally intensive and designed for whole-body alignment, while methods optimised for large-scale integration may be more scalable to multiple datasets and more species.

It is unclear which strategy for cross-species integration yields the most informative results and how the selection of homologous genes and integration algorithms affect the observable relationships between cells. This is because strategies have not been tested head-to-head across a variety of organs, species and biological systems. Furthermore, the number of species involved in the integration and the time that species have diverged can impact the output characteristics differently among strategies.

To address this, we developed the BENchmarking strateGies for cross-species integrAtion of singLe-cell RNA sequencing data (BENGAL) pipeline. We ran BENGAL to examine 28 cross-species integration strategies, covering 4 ways of mapping genes by homology and 10 integration algorithms in a variety of biological systems.

A key premise required to examine the validity of integration output is high-quality annotations of cell types and confident cell type homology between species. In this study, we focus our analysis on the known one-to-one homologous cell types in vertebrates. We reason that only if known one-to-one homologous cell types are well-integrated, is it then possible to perform deeper clustering analysis to identify cell types at a higher granularity or to decipher complex cell type taxonomy. Hence, an informative integration should achieve mixing of homologous populations between species, separation of species-specific populations, and maintenance of cell type distinguishability on the latent representation.

To quantitatively analyse integration results, we use established batch correction metrics and biology conservation metrics to analyse the integration of homologous cell types between species, and the loss of biological heterogeneity, respectively. We develop a new biology conservation metric to address loss of cell type distinguishability due to integration. We further perform cross-annotation of cell types using the integrated data to determine if the integration enables cell type annotation transfer. Finally, by observing the behaviour of different integration strategies in tasks with certain examination points, we provide guidelines for appropriate strategy selection aligned to cross-species integration goals.

## Results

### Integration of cross-species scRNA-seq data

The BENGAL pipeline performs cross-species integration and assessment of integration results using 28 strategies, as illustrated in Fig. 1. Quality control (QC) and curation of cell ontology annotation of input data is required prior to running the pipeline and is performed in an input-specific manner. (see Methods for details and recommended practices). During integration, the pipeline first translates orthologous genes between species using ENSEMBL multiple species comparison tool ^31^ and concatenates raw count matrices from different species. We compared three approaches for cross-species gene homology mapping: mapping using only one-to-one orthologs; mappings including one-to-many or many-to-many orthologs by selecting those with a high average expression level, or with a strong homology confidence (see Methods). Notably, LIGER UINMF also takes unshared features, therefore genes without annotated homology are added on top of the mapped genes. SAMap requires *de-novo* reciprocal BLAST to construct a gene-gene homology graph and we therefore followed the standalone workflow. We fed the concatenated raw count matrix to 9 integration algorithms, some of which were top-performers from previous benchmarking of data integration ^17^, including fastMNN, Harmony, LIGER, LIGER UINMF, Scanorama, scVI, scANVI, SeuratV4CCA and SeuratV4RPCA.

**Figure 1.**
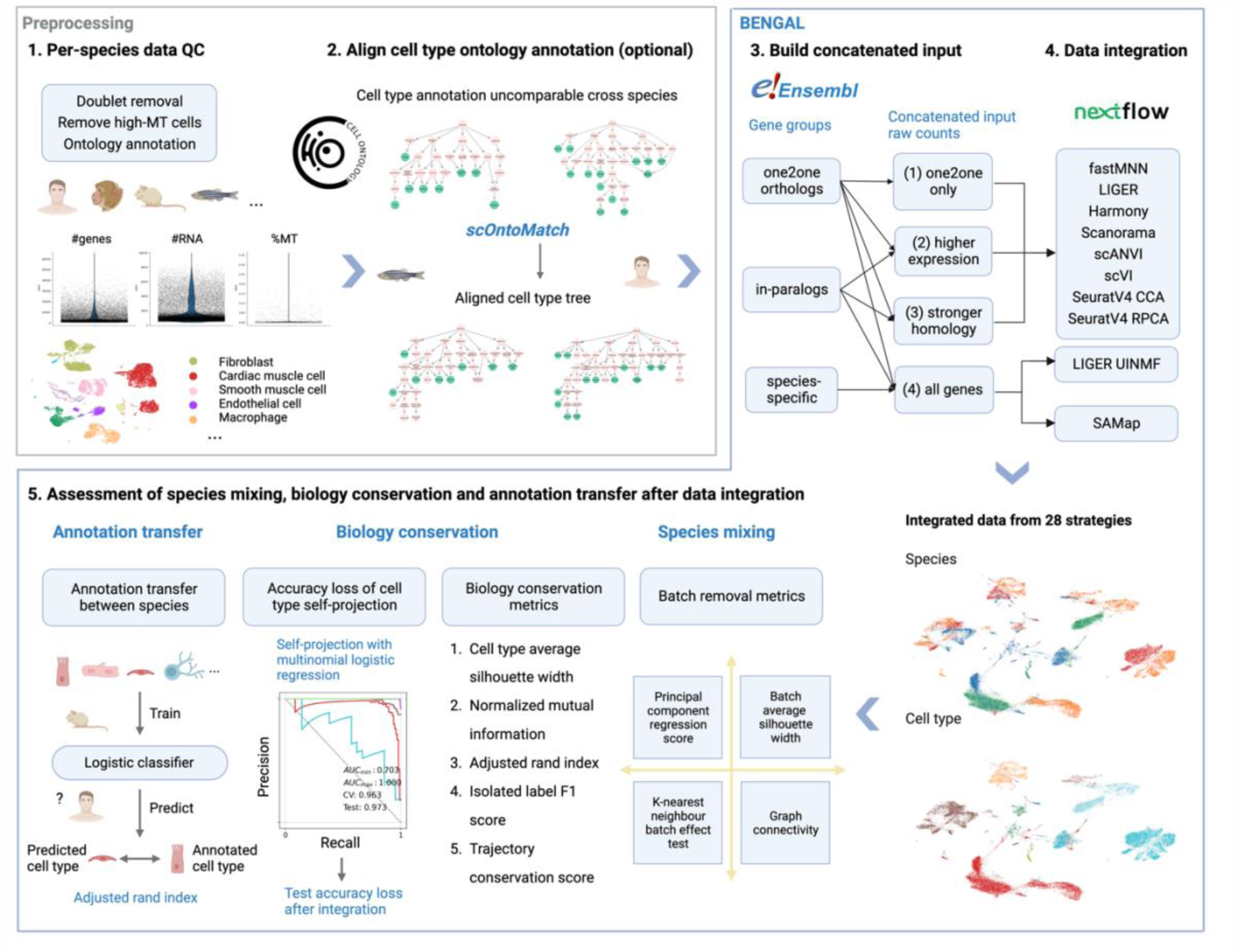
Schematic of BENGAL pipeline. 1) Quality control of input data is performed prior to data integration and is not part of BENGAL. Potential doublets and low-quality cells expressing high mitochondrial genes should be removed. Cell ontology annotation is collected from atlases or data portals, or curated from the originally published annotation 2) When ontology annotation granularity is incomparable across datasets, one-to-one homology between cell types needs to be robustly aligned. We developed scOntoMatch to find the appropriate annotation granularity and align cell type hierarchies across given datasets (see Methods). 3) Genes are grouped and translated across species by homology defined in ENSEMBL multiple species comparison tool. Raw count matrices are then concatenated across species using four possible homology matching methods respective to method inputs. 4) Run 9 integration algorithms to generate integrated output. 5) Perform integration assessment from species mixing, biology conservation and cell type annotation transfer. BENGAL, BENchmarking inteGration strAtegies for cross-species anaLysis of single cell transcriptomics data; QC, quality control; MT, mitochondrial; SCCAF, single cell clustering assessment framework; AUC, area under the curve; CV, cross-validation. Created with BioRender.com.

### Output assessment

Integration outputs were assessed from three main aspects, species mixing, biology conservation and annotation transfer. We computed and compared 4 batch correction metrics and 5 biology conservation metrics for species mixing and biology conservation, respectively (Fig. 1, see Methods for metrics selection and applicability, see Supplementary Notes for metrics principle). We also calculated these metrics in 3 types of homology concatenated, unintegrated data, to put the raw scores into context. Metrics scores in integrated data from 27 strategies and in unintegrated data were min-max scaled per task to equalise their discriminative power (see Methods for scaling detail). The species mixing score is the average of applicable scaled batch correction metrics, while the biology conservation score is the average of applicable scaled biology conservation metrics. The integrated score is a weighted average of species mixing score and biology conservation score with a 40/60 weighting ^17^. It is worth noting that batch correction metrics are not applicable SAMap outputs, so we did not include it in the scoring and ranking (see Supplementary notes for reasons). Instead, we show Uniform Manifold Approximation and Projection (UMAP) for visual inspection and alignment score to quantify the percentage of cross-species neighbours (see Methods for alignment score calculation).

Importantly, we developed a new metric, Accuracy Loss of Cell type Self-projection (ALCS), to target the most unwanted artefact of cross-species integration. ALCS quantifies the degree of blending between cell types per-species after integration, thus indicating the tendency of overcorrection of cross-species heterogeneity that may obscure species-specific cell types. We employed the self-projection concept in machine learning implemented in our previous work Single Cell Clustering Assessment Framework (SCCAF)^32^ and used the test accuracy of self-projection to indicate how well cell type labels are distinguishable from each other on a given feature representation. The loss of self-projection accuracy after integration, compared with unintegrated per-species data, measures the loss of cell type distinguishability due to integration (see Methods for further details of ALCS). We report ALCS separately to highlight its importance in our cross-species integration setting.

For annotation transfer, a multinomial logistic classifier built upon the SCCAF framework was trained on one species and used to annotate cell types in another species based on the shared features in integrated embedding. We assessed annotation transfer by calculating Adjusted Rand Index (ARI) between the original and transferred annotation (see Methods for details).

### Benchmarking metrics of different integration strategies varied widely

We ran the BENGAL pipeline for 16 cross-species integration tasks spanning a variety of biological scenarios to observe species mixing and biology conservation (see Table 1 for task design and Methods for tasks abbreviation). We explored performance of the different strategies in 3 different adult tissues, namely pancreas, hippocampus and heart, as well as in whole-body embryonic development. Using heart data from five species, we explored the upper limit of the number of species to include in one integration with 4 tasks. In addition, we performed pairwise integration to examine the impact of divergent time between species on integration output with 10 tasks.

**Table 1.** Overview of the reference tasks for benchmarking strategies for cross-species integration of scRNA-seq data. The "Number of cells per batch" refers to the cells that have passed the quality control criteria in the original literature and were included in the integration analysis for each integrated batch. The "Number of homologous genes analysed in this study" pertains to the genes that were analysed in the integration tasks. Only genes that meet the following criteria were included: 1) quantified in all datasets involved, 2) can be mapped to an ENSEMBL gene ID, and 3) have homology annotation across all studied species. To identify one-to-one orthologs and in-paralogs, including one-to-many and many-to-many orthologs, we used the ENSEMBL multiple species comparison tool (version 106) and accessed it via biomaRt (v2.46.3).

|  | Task name | Species | Technology | Number of cells per batch | Number of homologous genes analysed in this study |  | Challenge presented |
| --- | --- | --- | --- | --- | --- | --- | --- |
|  |  |  |  |  | one2one | one2many and many2many |  |
| 1 | Pancreas_hs_mm<sup>44</sup> | *Homo sapiens*,<br>*Mus musculus* | inDrop | hs: 8,402; mm: 1,875 | 11,248 | 664 | Basic performance |
| 2 | Hippocampus_hs_mu_ss<sup>33</sup> | *Homo sapiens*,<br>*Macaca mulatta*,<br>*Sus scrofa* | snRNA-seq | hs: 9,170, 8,214, 7,907, 6,893, 6,091, 5,555; mu: 19,092, 8,678, 8,337; ss: 13,168, 12,869, 10,814 | 12,557 | 24 | Basic performance, large data size, complex subcluster structure |
| 3 | Embryo_xt_dr<sup>34,35</sup> | *Xenopus tropicalis*,<br>*Danio rerio* | inDrops | xt: 123,632; dr: 36,627 | 5,188 | 1,749 | Whole-body data, less well-annotated homology, developmental trajectory |
| 4 | Heart_hs_mf<sub>1,3</sub> | *Homo sapiens*,<br>*Macaca fascicularis*, *Mus musculus* | scRNA-seq with 10× Genomics 3' V3.1 droplet-based sequencing (*H.sapiens*), snRNA-seq with DNBelab C Series Single-Cell Library Prep Set (*M.fascicularis*) | hs: 12,747; mf: 10,465 | 12,998 | 398 | Cross-study and cross-technology integration, gradual addition of species in one integration, far evolutionary distance, challenging homology annotation |
| 5 | Heart_hs_mf_mm<sub>1,3,36</sub> | *Homo sapiens*,<br>*Macaca fascicularis* | same with 4, scRNA-seq with 10× Genomics 3' V2 (*M.musculus*) | same with 4, mm: 7,402 | 11,202 | 199 |  |
| 6 | Heart_hs_mf_mm_xl<sub>1,3,36,45</sub> | *Homo sapiens*,<br>*Macaca fascicularis*, *Mus musculus*, *Xenopus laevis* | same with 5, microwell-seq (*X.laevis*) | same with 5, xl: 21,427 | 6,609 | 99 |  |
| 7 | Heart_hs_mf_<br>mm_xl_dr<br>1,3,5,36,45 | *Homo sapiens, Macaca fascicularis, Mus musculus, Xenopus laevis, Danio rerio* | same with 6, microwell-seq (*D.rerio*) | same with 6, dr: 3,193 | 4,267 | 82 |  |

To provide an overview of strategies’ performance, we calculated the mean and standard deviation of integrated scores for all strategies using the 7 reference tasks (excluding pairwise integration tasks as they overlap with the heart tasks, Table1, Fig. 2a). Overall, major differences were given by integration algorithms but not homology methods. Strategies that achieved successful integrations stayed largely consistent across tasks, while the relative species mixing and biology conservation scores varied. In general, scANVI, scVI, SeuratV4 CCA, SeuratV4 RPCA, harmony and fastMNN performed well across the board. For each task, we analysed strategies’ relative species mixing and biology conservation (Fig. 2b). scVI, harmony and SeuratV4 RPCA ranked top3 in terms of average species mixing across tasks, while for biology conservation, it was scANVI, SeuratV4 CCA and SeuratV4 RPCA (Extended Data Table 1). Across all tasks, we observed a trade-off between species mixing and biology conservation in most strategies (Fig. 2b, see Extended Data Fig. 1-16 for detailed metrics and scores of each task). SAMap performed well in all tasks, yielding strong cross-species alignment (see UMAP visualisation and alignment score in Extended data Fig. 17).

**Figure 2.**
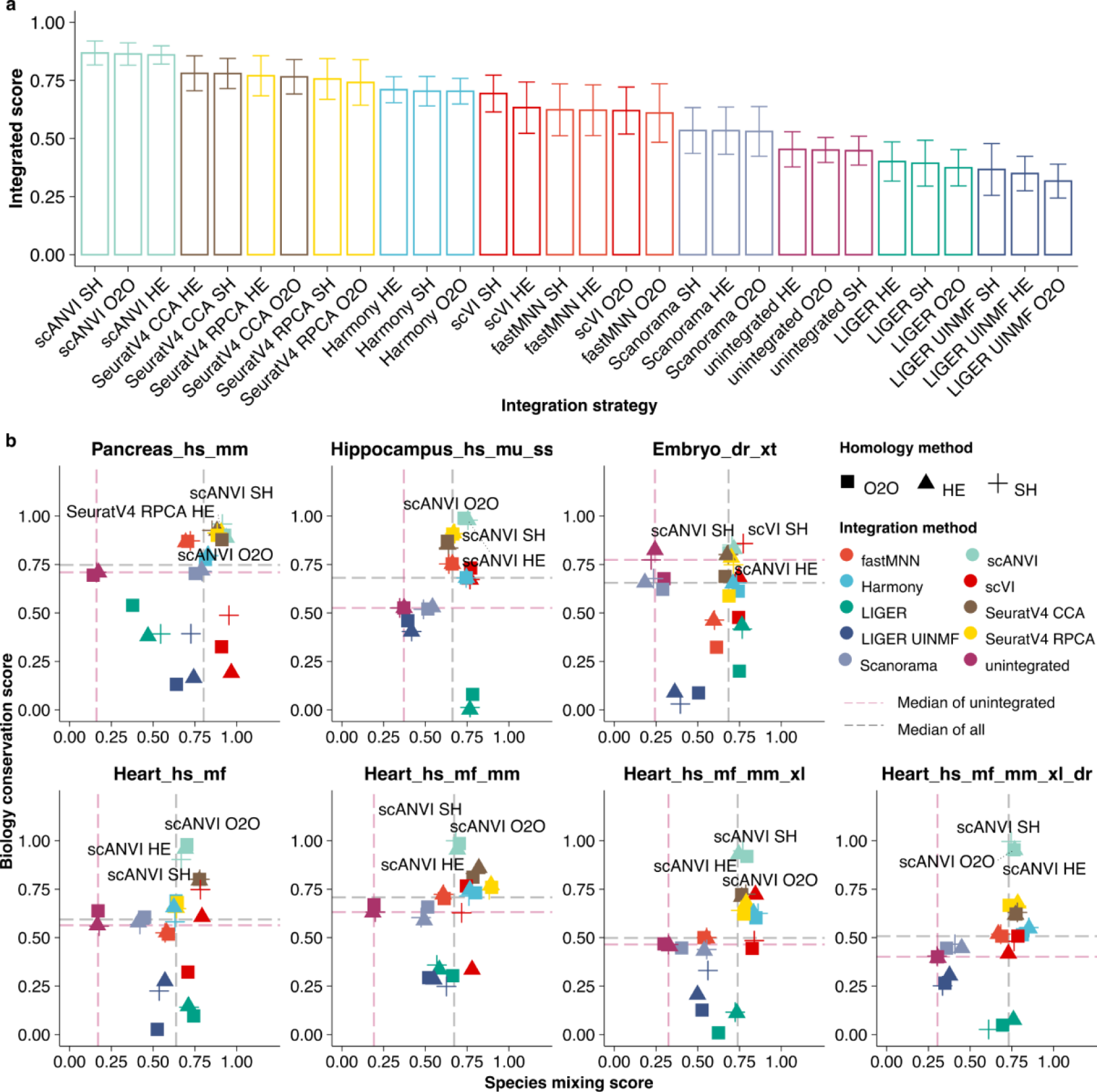
Benchmarking scores and metrics of different integration strategies on cross-species analysis. a) We used 4 batch removal metrics and 5 biology conservation metrics to examine the output of 28 data integration strategies and rank the strategies based on 7 cross-species data integration tasks (see Methods for metrics and tasks details). Batch removal metrics and biological conservation metrics are min-max scaled per task. The species mixing score is the average of scaled batch removal metrics applicable to the output type of the integration algorithm, while the biological conservation score is the average of scaled applicable biology conservation metrics to the output type and to the integration task (see Methods for metrics and scaling details). The integrated score is a weighted average of species mixing score and biology conservation score with 40/60 weight, respectively. The mean integrated score across 4 tasks is represented by the height of the bar plot and error bars indicate standard deviation. b) the species mixing score and biology conservation score of 7 example tasks are shown (see Table 1 for task details and Methods for task naming). O2O, only use one-to-one orthologs; HE, one-to-one orthologs plus one-to-many and many-to-many orthologs matched by higher average expression level; SH, one-to-one orthologs plus one-to-many and many-to-many orthologs matched by stronger homology confidence. hs, *Homo sapiens*, human; mf, *Macaca fascicularis*, long-tailed macaque; mu, *Macaca mulatta*, rhesus macaque; mm, *Mus musculus*, mouse; ss, *Sus scrofa*, pig; xl, *Xenopus laevis*, african clawed frog; xt, *Xenopus tropicalis*, western clawed frog; dr, *Danio rerio*, zebrafish.

### Loss of cell type distinguishability after integration

After examining output characteristics using established metrics for species mixing and biology conservation, we calculated ALCS to observe the conservation of cell type distinguishability. Fig. 3a shows the ALCS of various strategies in 7 example tasks, ordered by increasing maximum divergence time among integrated species. We found that as the integration involves more species that have diverged for a longer time, there is a general increase in ALCS across all strategies, but to a different extent. Overall, LIGER, LIGER UINMF and fastMNN have noticeably higher ALCS. As an example, we show the Uniform Manifold Approximation and Projection (UMAP) visualisation of scANVI HE and LIGER O2O integration of the hippocampus_hs_mu_ss task (Fig. 3b). Though both methods were capable of species mixing, LIGER merged several groups of cell types, for instance among EC, CA1 Sub, GC and Oligo (red dashed circle), as well as Endo with Vas (black dashed circle), whereas these cell types were kept distinguishable in scANVI. The relative ALCS stayed consistent in the 10 pairwise integration strategies of species in the heart tasks, where we also observed higher ALCS between species pairs that have diverged for a longer time (Extended data Fig. 18).

**Figure 3.**
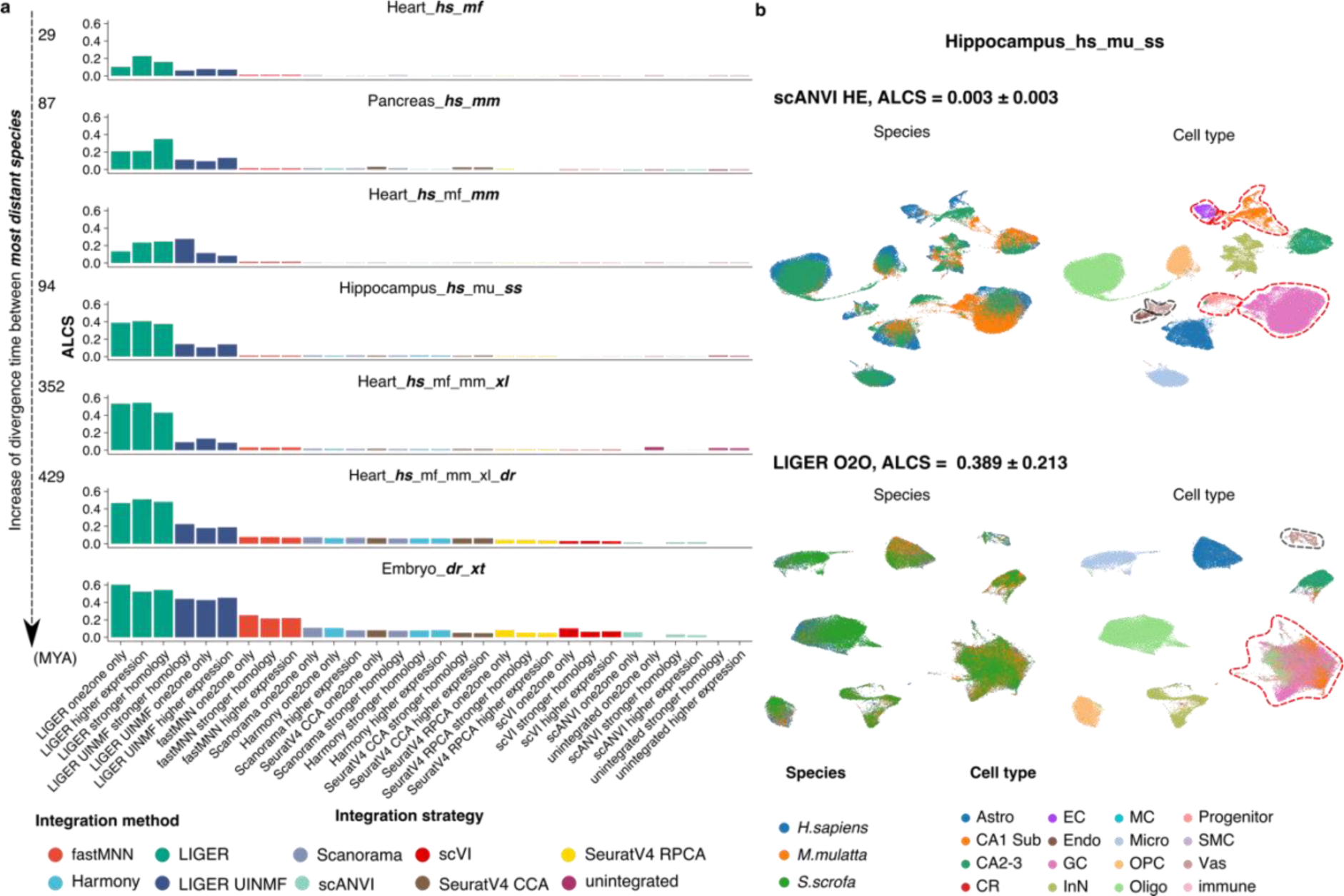
Loss of cell type distinguishability after integration is measured by accuracy loss of cell type self-projection. a) the ALCS of different integration strategies in 7 cross-species integration tasks. ALCS is the decrease in test accuracy of a cell type classifier trained on integrated data, compared with unintegrated data (see Methods for details). A high ALCS suggests that cell types become indistinguishable from each other after the integration and indicates overcorrection. Tasks are ordered by increasing divergent time between the most distant species (bold oblique). b) example of strategies with high and low ALCS in the Hippocampus_hs_mu_ss task. In scANVI HE integrated data, all cell types are clearly distinguishable after integration. However, In LIGER O2O results, EC, CA1 Sub, GC and some Oligo (red dashed circle), as well as Endo with Vas (dark grey dashed circle) became merged. ALCS, accuracy loss of cell type self-projection; MYA, million years ago; O2O, only use one-to-one orthologs; HE, one-to-one orthologs plus one-to-many and many-to-many orthologs matched by higher average expression level; SH, one-to-one orthologs plus one-to-many and many-to-many orthologs matched by stronger homology confidence. hs, *Homo sapiens*, human; mf, *Macaca fascicularis*, long-tailed macaque; mu, *Macaca mulatta*, rhesus macaque; mm, *Mus musculus*, mouse; ss, *Sus scrofa*, pig; xl, *Xenopus laevis*, african clawed frog; xt, *Xenopus tropicalis*, western clawed frog; dr, *Danio rerio*, zebrafish. Astro, astrocytes; CR, Cajal-Retzius cells; EC, entorhinal cortex; Endo, endothelial cells; GC, granule cell; InN, inhibitory neurons; MC, mossy cell; Micro, microglia cells; OPC, oligodendrocyte progenitor cells; Oligo, oligodendrocytes; NP, neuronal progenitors; SMC, smooth muscle cells; Vas, vasculature.

We conclude that cell type distinguishability is in general at risk when integrating evolutionarily distant species, but scANVI and scVI can produce more robust results than others. When the goal of cross-species integration is to uncover cell type homology, it is important to choose methods with low ALCS, to draw correct interpretations of cell type similarity based on the integrated embedding.

### Integration of human and mouse pancreas, and human, rhesus macaque and pig hippocampal-entorhinal system

We demonstrate BENGAL, starting from the integration between human and mouse pancreas (Pancreas_hs_mm task, Fig. 4a-c), and human, rhesus macaque and pig hippocampal-entorhinal system ^33^ (Hippocampus_hs_mu_ss task, Fig. 4d-f). These two scenarios have clearly distinguishable cell types and evident one-to-one cell type homology. The data were generated in the same study thus reducing technical batch effects. In comparison with the Pancreas_hs_mm task, the Hippocampus_hs_mu_ss task challenged the strategies with a larger data size and more complex subcluster structures (Table 1).

**Figure 4.**
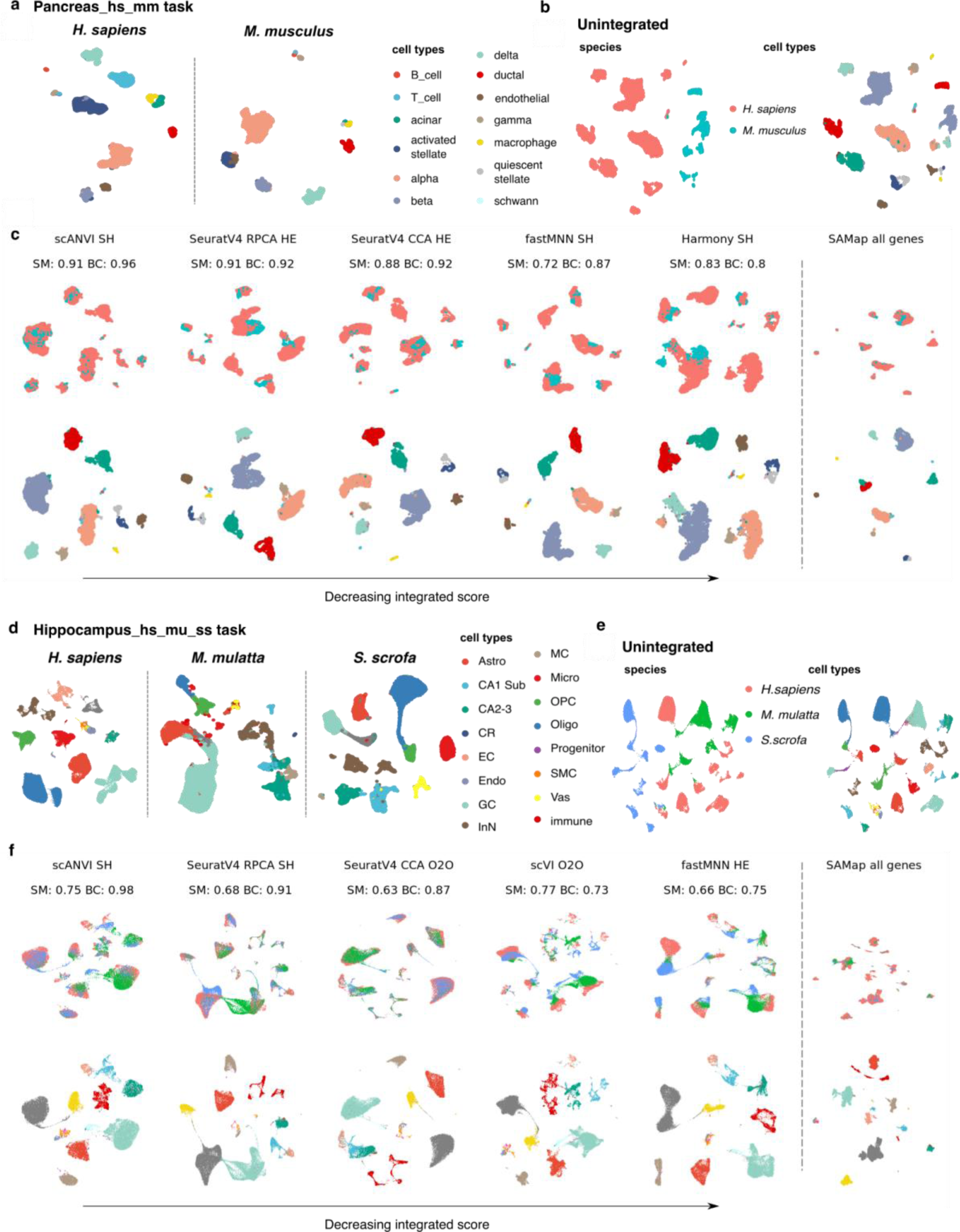
Integration results of human and mouse pancreas and human, rhesus macaque and wild boar hippocampal-entorhinal system. a) UMAP visualisation of cell types in human and mouse pancreatic tasks. b) UMAP visualisation of unintegrated data of the pancreas task. In unintegrated data, cells cluster primarily by species and homologous cell types form different species from separate clusters. One-to-one orthologs between human and mouse were used to concatenate raw count matrices, then processed by standard analysis pipeline (see Methods). c) UMAP visualisation of integration results of five top-performing algorithms and SAMap of the pancreas task, organised by decreasing integrated score and coloured by species or cell type. All strategies were ranked by integrated score and for the top 5 integration algorithms, the homology strategy with the highest integrated score was shown. The respective species mixing and biology conservation scores were also shown. The scores were not calculated for SAMap (not applicable, see Methods for details). D-F, UMAP visualisation of original data, unintegrated data and integration results of five top-performing strategies and SAMap of the hippocampus task. SM, species mixing score; BC, biology conservation score; O2O, only use one-to-one orthologs; HE, one-to-one orthologs plus one-to-many and many-to-many orthologs matched by higher average expression level; SH, one-to-one orthologs plus one-to-many and many-to-many orthologs matched by stronger homology confidence. hs, *Homo sapiens*, human; mu, *Macaca mulatta*, rhesus macaque; mm, *Mus musculus*, mouse; ss, *Sus scrofa*, pig; Astro, astrocytes; CR, Cajal-Retzius cells; EC, entorhinal cortex; Endo, endothelial cells; GC, granule cell; InN, inhibitory neurons; MC, mossy cell; Micro, microglia cells; OPC, oligodendrocyte progenitor cells; Oligo, oligodendrocytes; NP, neuronal progenitors; SMC, smooth muscle cells; Vas, vasculature.

Prior to integration, cells clustered strongly by species of origin (Fig. 4b, e). After integration and assessment, we observed diverse outputs from different strategies (Fig. 4c, f, Extended Data Fig. 1, 2). Overall, the integrated scores ranged between 0.33-0.94 for the Pancreas_hs_mm task or 0.31-0.89 for the Hippocampus_hs_mu_ss task. Batch correction score was between 0.14 and 0.97 or 0.35 and 0.78, and biology conservation score between 0.13 and 0.96, or 0 and 0.99, for the pancreas or Hippocampus_hs_mu_ss task, respectively. In both cases, harmony, fastMNN, scANVI, scVI, SeuratV4 CCA, SeuratV4 RPCA and SAMap were able to integrate the data while scanorama, LIGER and LIGER UNIMF did not integrate some homologous cell type pairs, such as beta cells and delta cells (Supplementary Fig. 1). Subcluster structures were visible on the UMAPs by scVI/scANVI but not on outputs by other methods in the Hippocampus_hs_mu_ss task (Fig. 4f). We hypothesize that heterogeneity within each cell type were better preserved in these deep learning-based methods, compared with nearest neighbour-based methods. SAMap gave a strong cross-species alignment, leading to a high degree of overlap between shared cell types on the UMAPs.

### Integrating embryonic development between xenopus and zebrafish

Next, we integrated the whole-body embryonic development process between xenopus and zebrafish. This is more challenging than integrating mature tissues because cell types are less distinct and a continuous trajectory exists among cells from different developmental stages. Based on mapped homologous cell types and recorded hours post fertilisation (HPF) between a zebrafish and xenopus embryonic development dataset ^34, 35^, we examined species mixing and biology conservation of various integration outputs (Fig. 5a, b). We observed an overall lower rate of success across strategies in the Embryo_dr_xt task. Algorithms that achieved higher integrated scores than unintegrated data include scVI/scANVI, SeuratV4 methods and Harmony, and they have comparable species mixing and biology conservation scores (Fig. 5b). However, due to the homology matching process, maximum 6,937 homologous genes among 17,330 and 9,661 genes were included in the analysis for zebrafish and xenopus, respectively, while SAMap were able to include all genes due to its utilisation of a gene-to-gene BLAST graph.

**Figure 5.**
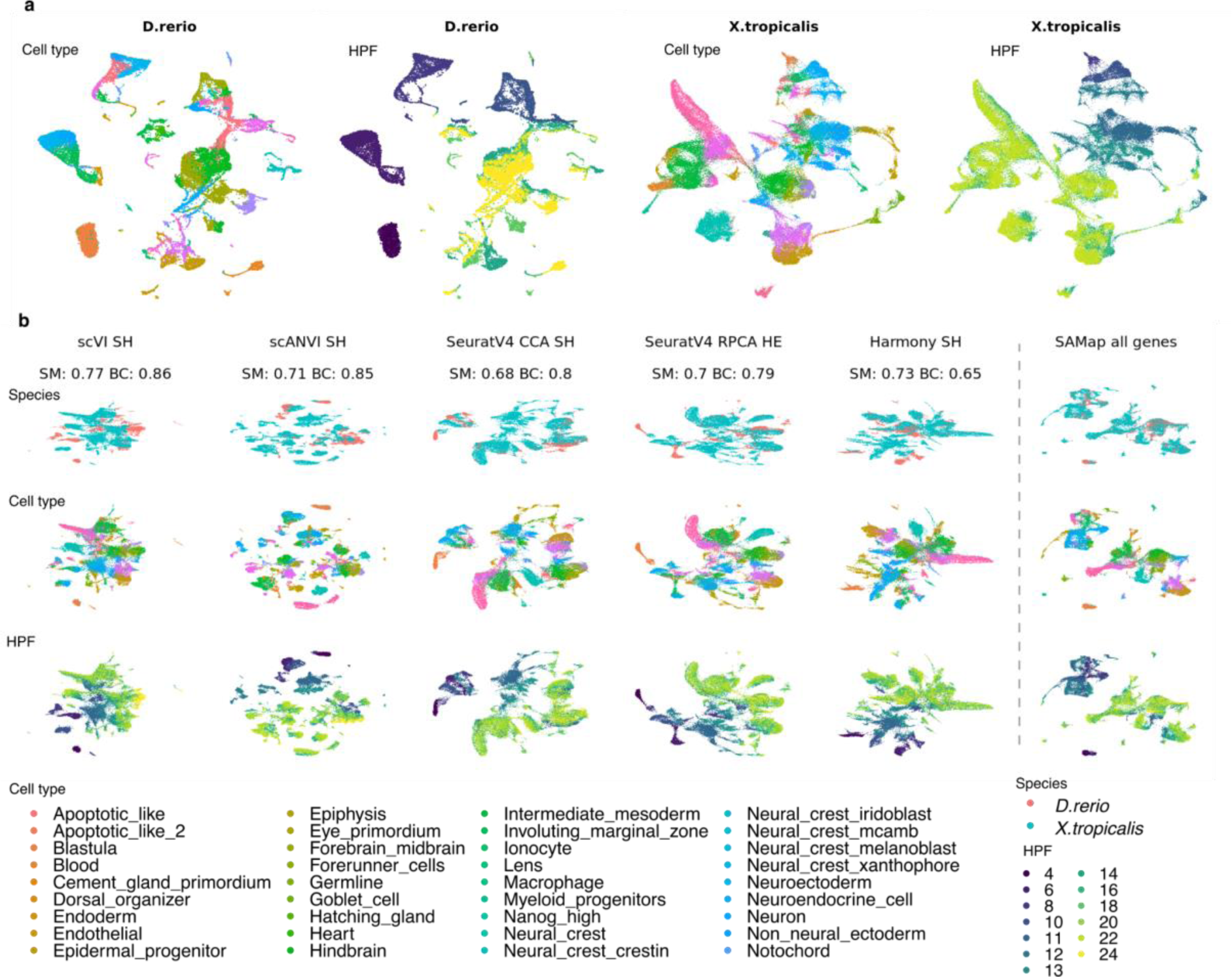
Integration of embryonic development between xenopus and zebrafish. a) UMAP visualisation of unintegrated data of xenopus and zebrafish embryonic development. Cells are coloured by their cell types and the developmental stages as in HPF. b) UMAP visualisation of integration results of five top-performing algorithms and SAMap, organised by decreasing integrated score and coloured by species or cell type. All strategies were ranked by integrated score and for the top 5 integration algorithms, the homology strategy with the highest integrated score was shown. The respective species mixing and biology conservation scores were also shown. The scores were not calculated for SAMap (not applicable, see Methods for details). SM, species mixing score; BC, biology conservation score; HPF, hour post fertilisation.

Hence, when integrating species with less well-annotated gene homology, SAMap can keep a much more complete profile of cellular expression when successfully integrating the data. On the other hand, we found that SAMap had a relatively low trajectory conservation score in SAMap (Extended Data Fig. 19). As seen on the SAMap UMAP, homologous cell types were strongly linked to each other and cells from different developmental stages were partially mixed. However, methods such as scVI/scANVI and SeuratV4 displayed a more explicit progression of developmental stages along the embedding (Fig. 5b). This observation suggests that SAMap focuses on prioritising identity-defining expression characteristics that stitch together evolutionarily related cell types.

### Cross-atlas integration of heart and aorta among five species

We designed a challenging task that is to integrate the heart and aorta tissue from five species: human, long-tail macaque, mouse, xenopus and zebrafish. While the mouse data were from an independent study to cover more cell types ^36^, all other species data were from current multi-organ atlases. We performed the integration by gradually adding one more species, starting from human and long-tail macaque, to explore the upper limit of the number of species to generate informative integration results. We have also performed pairwise integration of all species to examine the impact of divergence time on integration results.

When comparing cell type annotations between atlases, we noticed that the annotation granularity varied greatly (Extended Data Fig. 20-24). To robustly match homologous populations between atlases based on existing expert annotation, we employed the cell ontology (CL), a species-neutral, controlled vocabulary system that describes the ancestor-descendant relationship of cell type terms. CL is the current standard vocabulary used in both the Human Cell Atlas ^37^ and the EBI Single Cell Expression Atlas ^38^ for cell type annotation. Leveraging the hierarchical organisation of CL, we developed scOntoMatch, an R package which aligns the granularity of ontology annotations among scRNA-seq datasets to make them comparable across studies (see Methods for details of the scOntoMatch algorithm). Applying scOntoMatch to the heart data, we found the highest common granularity for cell type annotation in different integration runs.

After integration, scANVI reached the highest integrated score (0.62-0.69, Fig. 6a, individual metrics unscaled for each task for cross-task comparison. The same applies to the scores below). This was largely due to its high biology conservation score (0.70-0.83). On the other hand, SeuratV4 RPCA methods gave the highest species mixing score for the four tasks (0.46-0.54). For all tasks, scanorama, LIGER and LIGER UINMF had lower integrated scores than unintegrated data. In general, addition of species did not affect the relative performance of the integration algorithms and the species mixing scores, highlighting their robustness by these criteria (Fig. 6a). In contrast, biology conservation scores and ALCS dropped significantly (Fig. 6a, Fig. 3a), suggesting a general loss of biological signals of cell type specificities when distant species were added to the integration.

**Figure 6.**
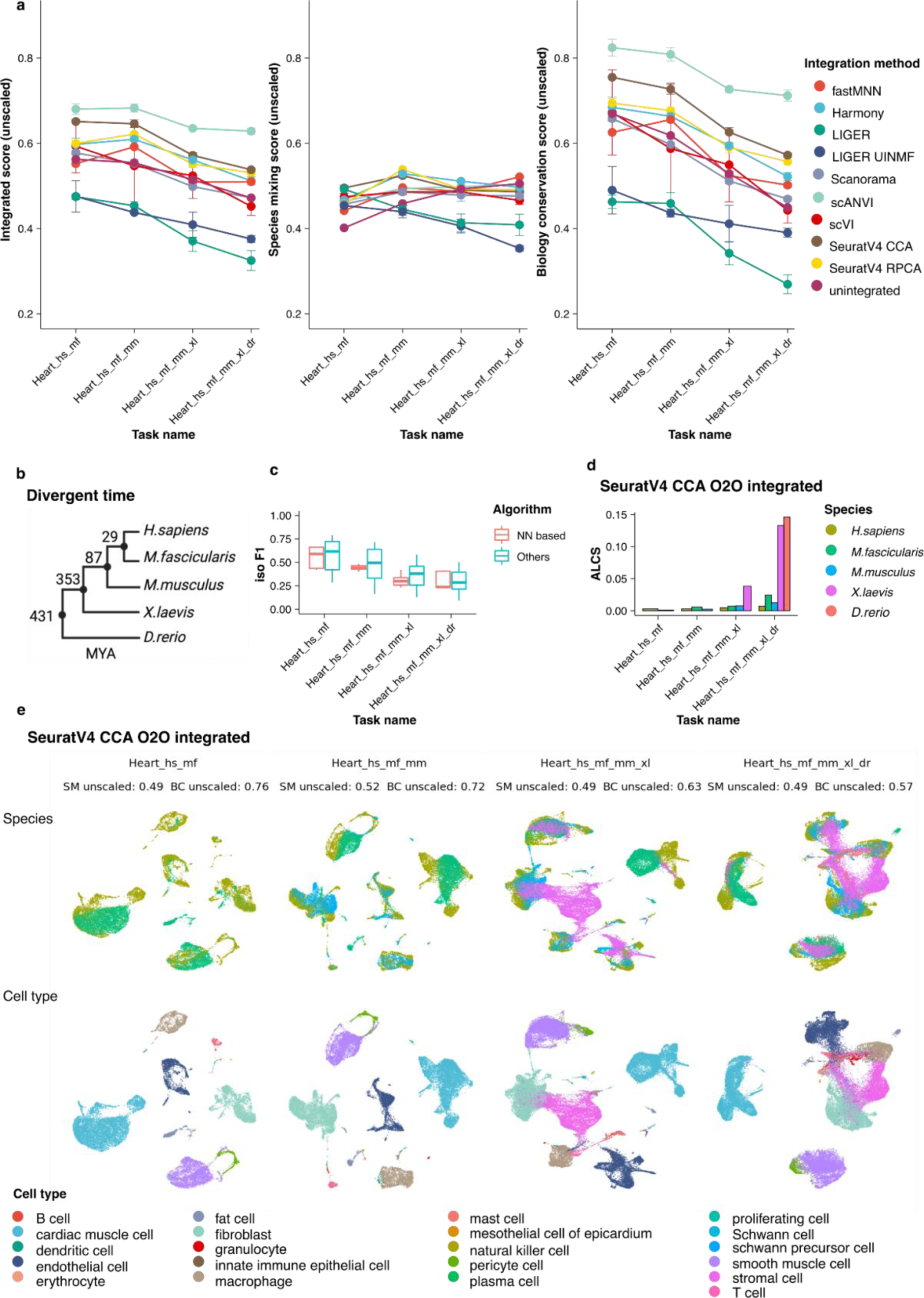
Integrating heart tissue from human, long-tail macaque, mouse, xenopus and zebrafish. a) Unscaled integrated score, species mixing score and biology conservation score of all strategies for the four heart tasks. The batch metrics and biology conservation metrics are not min-max scaled per task to enable cross-task comparison. Each point shows the average score across three homologous methods of each integration algorithm and whiskers indicate the standard deviation. b) the divergence time among the studied species in millions of years. c) The isolated label F1 score of different algorithms in the heart tasks. Nearest neighbour-based methods include fastMNN, SeuratV4 CCA and SeuratV4 RPCA. The bar in the boxplot shows the arithmetic average, lower and upper hinges correspond to the first and third quartiles and whiskers extend from the hinge to the largest value no further than 1.5 * interquartile range from the hinge. d) The ALCS of SeuratV4 CCA O2O integrated data per species of each heart task. High ALCS indicates a strong loss of cell type distinguishability due to integration (see Methods for details). e) UMAP visualisation of SeuratV4 CCA O2O integrated data in heart tasks, coloured by species and cell type. Unscaled SM and BC scores are also shown. Iso F1, isolated label F1 score; NN, nearest neighbour; ALCS, accuracy loss of cell type self-projection; SM, species mixing score; BC, biology conservation score; O2O, only use one-to-one orthologs; hs, *Homo sapiens*, human; mf, *Macaca fascicularis*, long-tailed macaque; mm, *Mus musculus*, mouse; xl, *Xenopus laevis*, african clawed frog; dr, *Danio rerio*, zebrafish.

Since only 8 among 22 cell types were shared by all species, we took this opportunity to investigate how different strategies handle cell types found in few datasets using isolated label F1 (iso_F1) score. We found that nearest neighbour-based methods have a tendency to overcorrect, leading to merging of similar cell types, especially when more species are involved in the integration (Fig. 6c). As an example, we show the SeuratV4 CCA integration of one-to-one orthologs only data for the four tasks (Fig. 6d, e). Among methods that achieved integration, SeuratV4 CCA has a low iso_F1 score, suggesting cell types specific to few species data were not well-separated (Extended Data Fig. 4-7).

**Figure 7.**
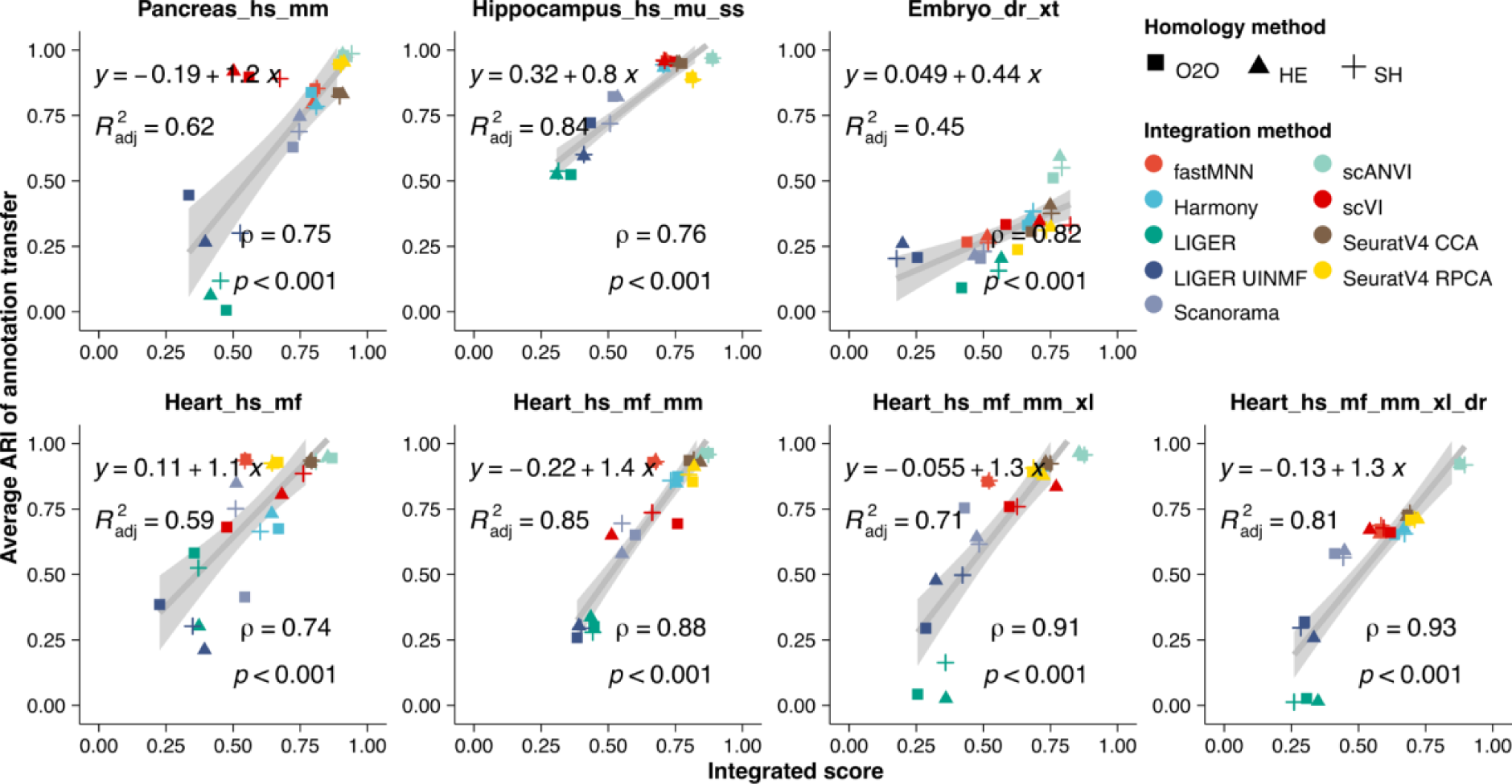
Adjusted rand index of cross-species cell type annotation transfer. The average ARIs between transferred annotation and original annotation in 7 reference tasks are shown for all integration strategies. A high ARI suggests a successful annotation transfer. ARI significantly positively correlates with the integrated score in all tasks (Spearman’s rank correlation significance P value < 0.001). Strategies are represented by points whose colour is the integration algorithm and whose shape is the homology method. ARI, adjusted rand index; 〉, Spearman’s rank correlation coefficient; p, P-value of Spearman’s rank correlation; adj, adjusted goodness-of-fit of the linear model; O2O, only use one-to-one orthologs; HE, one-to-one orthologs plus one-to-many and many-to-many orthologs matched by higher average expression level; SH, one-to-one orthologs plus one-to-many and many-to-many orthologs matched by stronger homology confidence; hs, *Homo sapiens*, human; mf, *Macaca fascicularis*, long-tailed macaque; mu, *Macaca mulatta*, rhesus macaque; mm, *Mus musculus*, mouse; ss, *Sus scrofa*, pig; xl, *Xenopus laevis*, african clawed frog; xt, *Xenopus tropicalis*, western clawed frog; dr, *Danio rerio*, zebrafish.

Increase of ALCS with addition of species was broadly observed for most strategies, indicating a loss of cell type distinction when many species data were involved in one integration (Extended data Fig. 25). For example, ALCS of SeuratV4 CCA O2O was roughly maintained between integration of human and macaque or human, macaque and mouse, but adding xenopus to the integration led to some loss of cell type distinction and this became global and stronger after zebrafish was integrated (Fig. 6d). Integrating many species that span a large evolutionary distance cannot therefore yield robust results for downstream analysis, such as *de novo* clustering, due to information loss during gene homology mapping and the integration process. Similarly, pairwise integration also suggested that integration is more prone to loss of cell type distinction between species that have diverged for a longer time (Extended Data Fig. 18).

### Cell type annotation transfer cross-species

One key application of cross-species integration is cell type annotation transfer: using annotated cell types in a well-studied species to label the cell types of a newly characterised species. To investigate the potential of different strategies to support annotation transfer, we performed pairwise cross-annotation in all tasks. In practice, we applied the SCCAF multinomial logistic classifier trained on one species to predict the cell type labels of another species. When the transferred labels covered all known sharded cell types and had an acceptable adjusted rand score (ARI) with the original label, we considered cross-annotation successful.

Overall, we observed that if data integration was well-achieved, annotation transfer was mostly successful (Fig. 7). In all tasks, strategies showed a significant positive correlation of integrated score with average ARI (Spearman’s rank correlation P value < 0.001). Compared with species mixing score, biology conservation score correlated stronger with ARI, suggesting that cross-annotation is facilitated by well-preserved cell type features (Extended Data Fig. 26). We noticed that with the addition of evolutionarily more distant species in the heart tasks, the overall annotation transfer quality did not drop for methods that achieved successful integration (Fig. 7). Inspection of UMAP visualisation of showed that transferred annotation was to a large extent accurate in methods such as scANVI and SeuratV4 CCA even between distant species, such as human and zebrafish (see Extended Data Fig. 27 for successful and unsuccessful examples). This is in line with the observation that top-performing methods stayed relatively robust with the addition of more species (Fig. 6a). Pairwise integration tasks show similar results that annotation transfer is more difficult between evolutionarily distant species for all strategies (Extended Data Fig. 28).

### Impact of one-to-many and many-to-many homologous genes on integration performance

We found that incorporating one-to-many and many-to-many homologous genes was beneficial when they accounted for a higher proportion of genes used in the integration. Specifically, in the Embryo_dr_xt task, algorithms that successfully integrated the data achieved higher integrated scores when in-paralogs were included, especially when the in-paralog with higher homology confidence was matched with the corresponding ortholog (Extended Data Fig. 29). The improvement was primarily driven by biology conservation scores, suggesting that including these genes preserved a more complete picture of cell type expression profiles that facilitated data integration. This observation is consistent with SAMap outperforming in the Embryo_dr_xt task, as SAMap used all genes and was better able to resolve the alignment between homologous cell types (Fig. 5, Extended Data Fig. 17). Although we did not observe significant improvements in benchmarking scores by adding in-paralogs in most tasks, we found that the homology method still influenced integration from the perspective of highly variable genes (HVGs). We observed different HVGs were selected, especially among evolutionarily distant species, when incorporating in-paralogs (Extended Data Fig. 30). Since integration algorithms other than SAMap require HVG selection, adding in-paralogs that contribute to cell type expression variation which would support the integration algorithms. Overall, our findings highlight the importance of carefully considering the inclusion of in-paralogs in data integration, as their impact varies depending on the specific task and species being studied.

## Discussion

We benchmarked 28 strategies for cross-species single-cell transcriptomics data integration, covering 4 ways to match genes cross-species by homology and 10 integration algorithms. Based on known cell type homology, we used 4 metrics to evaluate the degree of species mixing and 6 for scoring biology conservation. We also provided the following freely available tools: BENGAL, a Nextflow pipeline for cross-species scRNA-seq data integration and assessment of integration results; ALCS, a cross-species integration-focused biology conservation metric to quantify the loss of cell type distinguishability; scOntoMatch; an R package to help align the granularity of cell ontology annotation across datasets.

We found that deep neural network-based scVI/scANVI generally achieved an overall balance between species mixing and biology conservation. scANVI performed the best across the board, suggesting that taking cell type annotation into account is beneficial for the integration. On the other hand, SeuratV4 methods were able to integrate successfully for evolutionarily distant species but were more prone to overcorrection due to its nearest neighbour foundations. Algorithms based on NMF could not integrate some cell types and merged different cell types in basic scenarios, suggesting that they were less suitable for cross-species integration. We noticed that the tailor-made cross-species integration algorithm, SAMap, can pick up key similarities between homologous cell types that are otherwise difficult to detect in complex situations. The impact of homology method depended on the species involved in the integration. Different homology methods influenced HVG selection and had larger influences when there was a high proportion of one-to-many and many-to-many orthologs between the studied species. By performing cell type annotation transfer using integrated results, we observed that if integration is well-achieved, annotation transfer was to a large extent successful even between evolutionarily distant species such as human and zebrafish. Nonetheless, highly similar cell types could still be mislabelled between closely related species *e.g.* human and mouse.

Based on a diversity of cross-species integration scenarios investigated in this study, we provide the following guidelines on choosing the most appropriate algorithm for cross-species integration: for closely related species, scVI (scANVI when confident cell type annotation is available) or harmony carries out species mixing while preserving biological heterogeneity. For evolutionarily distant species, SeuratV4 methods can achieve strong species mixing and RPCA is more scalable than CCA for larger datasets ^17, 39^. For integration of whole-body atlases or between species lacking well-curated gene homology annotation, SAMap excel in aligning homologous cell types by resolving the gene homology mapping challenge. For species sharing a large amount of one-to-many and many-to-many orthologs, including them in the analysis can improve the integration as more information about cell type expression profiles are preserved.

Our quantitative scoring system of species mixing and biology conservation is supported by two assumptions: 1) cell type annotations are accurate; 2) known cell type homology from literature is confident. It is important to point out that our knowledge about cell types and their evolution is still rapidly growing ^8, 40^. Currently, cell type homology is discernible at decreased granularity with the increase of evolutionary distance between species due to both biological and practical reasons. Biologically speaking, more complex species can have a greater number of specialised cell types while all of them collectively are homologous to one multi-function cell type in their evolutionary ancestors. In practice, cell type annotation can only reach a relatively coarse level in less-studied species and tissues due to our limited understanding. In this study, we focussed on one-to-one homologous cell types and expected them to mix between species and stay distinctive among other cell types after integration. We demonstrated that leveraging the unified CL annotation system, we can find the annotation granularity at which one-to-one cell type mapping is established in homologous organs among vertebrates.

Future cross-species integration will require the development of novel integration strategies that overcome the limitation of gene orthology, are aware of the evolutionary homology between cell types and adopt a hierarchical structure of cell type classification. Currently, cross-species integration still heavily relies on curated gene orthology based on sequence. Most strategies also assume that orthologs have the same expression pattern in homologous cell types and play the same roles in distinguishing different cell types. From the standpoint of gene evolution, this has two caveats: 1) species-specific genes contribute to a large extent the novelty of species-specific cell types and should not be ignored when performing cross-species analysis ^11, 41^, 2) functional diversification between orthologs might be as common as between paralogs and out-paralogs can exhibit higher expression similarity compared to their corresponding orthologs through gain, loss and partition of function ^29, 42, 43^, Currently, only SAMap tackles the above by *de novo* homology mapping. However, being a heuristic approach, it does not provide explanations of the observed cell type expression similarity. Expression similarities between cells cross-species may arise for reasons other than evolutionary homology, such as convergence and concerted evolution ^8^. Novel algorithms are required to disentangle the source of the expression similarities, to achieve deeper understandings of cell types ^10^. Moreover, one-to-one homologous relationship between cell types might not hold for evolutionarily distant species, as cell types have undergone complex evolution of identity-defining gene networks ^29^. Adopting a hierarchical representation of cells in the integration outcome can benefit the comprehension of the relationship between cell types across species. According to the heart example, we conclude that when data integration is performed between non-mammals with mammals, the result can still serve as basis for cell type annotation transfer, but the embedding might be not suitable for *de novo* clustering analysis due to strong loss of biology. Other analysis, such as expression correlation based independent analysis might be more informative ^9, 14^.

### Methods Abbreviations

We use consistent abbreviations for species and homology methods across this study for clarity and brevity. For species, we take the initials of the scientific name of the species: hs, *Homo sapiens*, human; mf, *Macaca fascicularis*, long-tailed macaque; mu, *Macaca mulatta*, rhesus macaque; mm, *Mus musculus*, mouse; ss, *Sus scrofa*, pig; xl, *Xenopus laevis*, african clawed frog; xt, *Xenopus tropicalis*,western clawed frog; dr, *Danio rerio*, zebrafish. For homology methods the abbreviations are: matching only one-to-one orthologs between species, O2O; matching one-to-one orthologs and in-paralogs with which have a higher expression value, HE; matching one-to-one orthologs and in-paralogs which have a higher homology confidence, SH; include all homologous genes as well as those without homology information, all genes. We name the integration tasks following the pattern: “tissue_species” and the integration strategies following the pattern “algorithm_homology_method”.

### Datasets

We downloaded raw count matrices and published annotations of the following datasets: Pancreas_hs_mm: inDrop data from human and mouse pancreas (GSE84133) ^44^; Hippocampus_hs_mu_ss: snRNA-seq data from human, macaque and pig hippocampal and entorhinal regions (GSE186538) ^33^; Heart tasks: heart and aorta tissue from human (https://figshare.com/projects/Tabula_Sapiens/100973) ^1^, long-tail macaque (https://db.cngb.org/nhpca/download) ^3^, mouse (https://www.ebi.ac.uk/biostudies/arrayexpress/studies/E-MTAB-8810) (only no compound treatment mouse data was used) ^36^, *Xenopus laevis* (https://figshare.com/articles/dataset/Cell_Atlas_of_the_Xenopus_Laevis_at_Single-Cell_Resolution/19152839) ^45^ and zebrafish (https://bis.zju.edu.cn/ZCL/) ^5^; Embryo_xt_dr: inDrops data from zebrafish (GSE112294) ^34^ and xenopus embryo (GSE113074) ^35^.

### Quality control of scRNA-seq datasets

We follow the quality control (QC) in the original studies for each dataset. We excluded cells that didn’t pass QC from the raw count data, according to published cell type annotations. We did not apply additional QC in this study nor is the QC step part of the BENGAL pipeline, since QC criteria are highly specific to each dataset and should be performed on case-by-case while observing the data.

### Reference transcriptomes

Transcriptomes used in the BLAST step in SAMap are downloaded from ENSEMBL, except that for xt in Embryo_dr_xt task the transcriptome was downloaded from Xenbase (https://www.xenbase.org/entry/static-xenbase/ftpDatafiles.jsp). Versions are in line with the version at original publication of each dataset: Pancreas_hs_mm task: mm (GRCm38) and hs (GRCh38); Hippocampus_hs_mu_ss task: hs (GRCh38), mu (Mmul_10), ss (Sscrofa11); heart tasks: hs (GRCh38), ma (Macaca_fascicularis_6.0), mm (GRCm39), xl (v9.2), dr (GRCz11); Embryo_dr_xt task: xt (Xtropicalisv9.0), dr (GRCz10).

### Curation of cell type annotation based on known homology

We take the published cell type annotation of all datasets used in this study and curate the cell type annotations into cell ontology ^37^terms. We paid specific attention that the same cell type identifier refers to homologous cell types *i.e.* cell types from different species and trace back to the same precursor cell type in a common evolutionary ancestor ^8^.

In the Pancreas_hs_mm and Hippocampus_hs_mu_ss tasks, cell type homology is evident between species and were matched in the original study. For the Hippocampus_hs_mu_ss task, NBs, RGLs and nIPCs from macaque and pig are collectively annotated as “neuronal progenitor”. In the human data, we collectively annotate CA1 and SUB as “CA1 SUB”, CA2, CA3 as “CA 2-3”, macrophage, microglia, myeloid cell and T cell as “immune”, aSMCs and vSMCs as “smooth muscle cell”, pericytes and vascular and leptomeningeal cell as “vasculature”, COP is combined with OPC. For the Embryo_dr_xt task, we used the annotations provided in the SAMap study ^29^, as they have manually matched one-to-one homologous cell types using multiple lines of evidence including developmental hierarchies. For the heart tasks, we leverage the hierarchical structure of cell ontology to align annotation granularity across datasets, so that homologous cell populations for a given set of species with curated ontology annotation are matched (see the ‘Aligning ontology annotation with scOntoMatch’ section below). Overall, we ensure one-to-one mapping of homologous cell types among the studied species, even though this might lose granularity and lead to slightly more coarse annotation in the heart tasks.

### The BENGAL pipeline

The BENGAL pipeline is built using Nextflow (v22.04.3) DSL2 in java (OpenJDK v11.0.9.1-internal) and utilises singularity containers ^46^ for portability and reproducibility, and is compatible with the execution on high performance computing (HPC) clusters. For data integration, python-based scripts were written based on Python (v3.9.13), Scanpy (v1.9.1), h5py (v3.7.0) and anndata (v0.7.5). We used python implementation of harmonypy (v0.0.5), scanorama (v1.7.2), scVI and scANVI (v0.15.0 with pytorch v1.12.1 and cudatoolkit v11.6 to support execution with Nvidia GPU), SAMap (v1.0.2). R-based scripts are based on R (v4.0.5). We used Seurat (v4.1.1), LIGER (v0.5.0) and LIGER UINMF (v1.1.0) and fastMNN (v1.12.3 from package batchelor). For SCCAF analysis, we made modifications to the SCCAF package v0.0.10 and provide a docker container in docker://yysong123/intgpy:sccaf. For batch correction metrics and biology conservation metrics calculation, we used scIB (v1.1.3).

### Translating features cross-species by gene homology

We used ENSEMBL multiple species comparison tool (version 106), accessed via biomaRt (v2.46.3), to annotate gene homology and translate ENSEMBL gene id cross-species. We chose ENSEMBL to be the primary database for gene homology annotation, since there is manually curated homology, in addition to the automated annotation pipeline, it is a popular choice among previous and current cross-species scRNA-seq studies and it is broadly accessible for biologists from various disciplines. BENGAL is conveniently transferrable to use other homology databases, such as EggNOG or BLAST, as long as a homology matching table can be provided.

### Construct concatenated cross-species count matrix

Starting from raw count matrices of scRNA-seq data from different species, the BENGAL pipeline first identifies gene groups composed of one-to-one, one-to-many or many-to-many orthologs and species-specific genes using ENSEMBL multiple species comparison tool. One-to-one orthologs are directly matched across species. On top of that, one-to-many or many-to-many orthologs within each gene group are added and matched by selecting those with higher average expression level calculated using scanpy.pp.calculate_qc_metrics, or with stronger homology confidence defined in ENSEMBL using available attributes among: orthology.confidence, Gene.order.conservation.score, Whole.genome.alignment.coverage, query.gene.identical.to.target, gene.identical.to.query.gene in respective order from biomaRt. The method LIGER UINMF also takes unshared features (species-specific genes) and these genes were added in addition to the three types of inputs with shared features. SAMap quantifies gene homology strength using *de novo* BLAST, so the raw count matrix and the full set of genes are given to the method without the above homology-matching step, and tblastx (from BLAST v2.12.0) was run using transcriptomes data to generate blast maps for the method.

### Standard processing of scRNA-seq data

Per-species data and unintegrated data were analysed following the scanpy standard analysis workflow with following parameters: sc.pp.highly_variable_genes(adata, min_mean=0.0125, max_mean=3, min_disp=0.5); sc.pp.scale(adata, max_value=10); sc.tl.pca(adata, svd_solver=’arpack’); sc.pp.neighbors(adata, n_neighbors=15, n_pcs=40); sc.tl.umap(adata, min_dist=0.3, spread=1). For the homology concatenated data input to integration algorithms, we select highly variable genes per integration key by sc.pp.highly_variable_genes(adata, min_mean=0.0125, max_mean=3, min_disp=0.5, batch_key=integration_key). This function calculates HVGs per batch and uses the intersection of HVGs across batches as input for PCA (see below for integration key per task).

### Cross-species data integration

Taking the concatenated matrix with cells from all species and features matched by homology as input, the pipeline runs 10 integration methods using default parameters.

fastMNN: we use the fastMNN from github.com/LTLA/batchelor, via SeuratWrappers (v0.3.0) following the tutorial http://htmlpreview.github.io/?https://github.com/satijalab/seurat-wrappers/blob/master/docs/fast_mnn.html.

harmony: we use scanpy.external.pp.harmony_integrate to run harmony following tutorial: https://scanpy.readthedocs.io/en/stable/generated/scanpy.external.pp.harmony_integrate.html LIGER: we use the LIGER method via SeuratWrappers (v0.3.0) following the tutorial: http://htmlpreview.github.io/?https://github.com/satijalab/seurat-wrappers/blob/master/docs/liger.html

LIGER UINMF: we run LIGER UINMF via github.com/welch-lab/liger following the tutorial http://htmlpreview.github.io/?https://github.com/welch-lab/liger/blob/master/vignettes/SNAREseq_walkthrough.html 

SAMap: we run SAMap using github.com/atarashansky/SAMap following the tutorial and vignette in the github repository.

Scanorama: we use scanpy.external.pp.harmony_integrate to run scanorama following tutorial: https://scanpy.readthedocs.io/en/stable/generated/scanpy.external.pp.scanorama_integrate.html

scANVI: we run scANVI following documentation https://docs.scvi-tools.org/en/stable/api/reference/scvi.model.SCANVI.html. Note that although scANVI can be initialised from a pre-trained scVI model, we built scANVI models from scratch using the input AnnData object. This is to make the scANVI runs independent of scVI runs for benchmarking purposes.

scVI: we run scVI from scvi-tools.org/ following tutorial https://docs.scvi-tools.org/en/stable/tutorials/notebooks/harmonization.html

SeuratV4CCA and SeuratV4RPCA: we run SeuratV4CCA from https://satijalab.org/seurat/index.html following https://satijalab.org/seurat/articles/integration_introduction.html and RPCA following https://satijalab.org/seurat/articles/integration_rpca.html

### Integration key

SAMap is designed to perform integration between data from different species so ‘species’ were used as the integration key. For the integration methods other than SAMap, we use ‘species’ as the integration key if there is only one batch per species (Embryo_dr_xt task) or per-species batches have unbalanced cell type composition (Pancreas_hs_mm task and Heart tasks), otherwise, we use ‘batch’ as integration key if each batch has balanced cell types (Hippocampus_hs_mu_ss task). Highly variable genes were selected using the same integration keys.

### Dimension reduction and visualisation

For methods output an embedding, neighbourhood graphs were calculated with n_neighbors=15, n_pcs=40, and the UMAP representations were computed using min_dist=0.3 and spread=1.0. Points were shuffled when plotting UMAP to avoid colour overlap. For methods that output a pseudo-count matrix, we first calculate PCA, then use the same parameters to calculate neighbourhood graph and UMAP. SAMap outputs a corrected neighbourhood graph, so we only recalculate UMAP using the same parameter for visualisation.

### Species mixing score and batch correction metrics

Species mixing is assessed by 4 established batch correction metrics for scRNA-seq. This includes the principal component regression (PCR), batch average silhouette width (bASW), graph connectivity (GC) and k-nearest neighbour batch effect test (kBET). In practice, we used the scIB package to calculate the metrics ^17^. All metrics are computed on the PCA embedding for methods returning a pseudo-count matrix (SeuratV4 methods, n_pcs=20), or the output embedding for methods returning a latent embedding (n_dims=20). For metrics that operate on kNN graphs, we calculate k=20 nearest neighbours on the PCA or embedding both using the first 20 dimensions. In addition, PCR does not rely on cell type annotation whilst the other metrics take cell type labels into account. We highlight the separation of these two types of methods in detailed scores and rankings of each task (Extended Data Fig. 1-7).

We compute these 4 metrics in 28 integration strategies and 3 unintegrated data of different homology strategies. For each metric, we perform min-max scaling to reflect the relative performance of different strategies in concordance with scIB ^17^. We included the unintegrated data in the scaling as a reference, to avoid minor differences in metrics between algorithms that performed similarly well enlarged by scaling.

Species mixing score is the average of the scaled 4 metrics. This is to balance the potential biases between different metrics as none of them are on itself a comprehensive evaluation of species mixing.

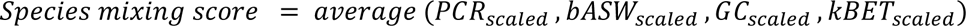

Unlike the study of scIB, we did not include batch graph Local Inverse Simpson’s Index (iLISI) score. This is because most integration outputs simply have a raw iLISI score near 0 (see Supplementary Fig. 17 for raw metrics distribution). We reason that integrated data could not show improvement in terms of iLISI score because cross-species difference is among the largest effects. All other batch correction metrics could show informative range of variation and using the average is the most appropriate for comparing between strategies. See supplementary notes for principles of the batch metrics, with further details available in the scIB study ^17^.

### Biology conservation score and metrics

Biology conservation is assessed by 5 biology conservation metrics for scRNA-seq, as well as ALCS, a novel metric we developed to reflect maintenance of cell type distinguishability. Biology conservation metrics include cell type ASW (cASW), normalised mutual information (NMI), ARI, iso_F1 score and trajectory conservation score (Traj, only applicable in the Embryo_dr_xt task).

Similarly with the calculation of batch correction metrics, we compute these 5 metrics in 28 integration strategies and 3 unintegrated data of different homology strategies using scIB and perform min-max scaling. Biology conservation score is the average of the scaled metrics.

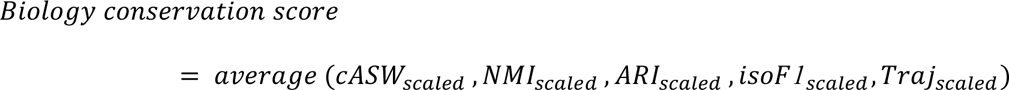

We did not include cell type graph LISI (cLISI) in scIB for a similar reason with iLISI: this score is easily fulfilled by most of the results in our study (Supplementary Fig. 17). All strategies achieved a cLISI close to 1 and thus is not informative to scale these metrics or include it in biology conservation score. See supplementary notes for principles of the biological conservation metrics, with further details available in the scIB study ^17^.

### Integrated score

Integrated score is the weighted average of species mixing score and biology conservation score. In concordance with the scIB study, we give 0.4 weight to species mixing and 0.6 weight to biology conservation.

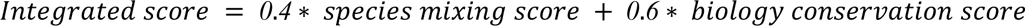

In this study, we present integrated scores that allow for comparison between integration strategies utilised in our cross-species integration tasks. It is important to note that these scores do not provide a comprehensive evaluation of the algorithms employed, as they were not initially designed for cross-species integration. These algorithms were able to perform cross-batch and cross-modality integration, as demonstrated by their benchmarking on these tasks in the scIB study ^17^.

### Alignment score

To quantitatively analyse the degree of cross-species alignment achieved by SAMap, we calculated the alignment score (AS) used in the SAMap study. AS is the average percentage of cross-species neighbours over the maximum number of possible neighbours across all cells from all species.

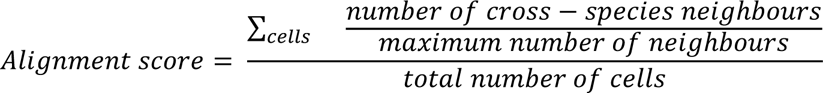

This reflects the degree of cross-species alignment and is comparable between SAMap and other strategies. We calculate AS on the kNN graph calculated from embeddings or PCA of pseudo-count matrices from other strategies with sc.pp.neighbors(adata, n_neighbours = 20, n_pcs = 20) and directly on the graph output of SAMap (SAMap ran with n_neighbours = 20). Across all tasks, SAMap shows significantly higher AS than other strategies, while other strategies show smaller improvement over unintegrated data (Extended Data Fig. 17). Undeniably, SAMap directly operates at kNN level to align cross-species neighbours, resulting in enhanced cross-species alignment. On the other hand, we treated inter-species and intra-species edges equally when kNN was calculated for other methods. Nevertheless, AS is in concordance with the observed strong cross-species alignment by SAMap from the UMAP visualisations.

### Assessing biology conservation with ALCS

We propose a new metric, ALCS, to assess the maintenance of cell type distinction after cross-species integration. This is motivated by our observation that integration algorithms tend to overcorrect and lose the separation of similar cell types on the integrated embedding, since cross-species difference is huge. This behaviour contradicts the goal of cross-species integration, as any species-specific population with subtle differences from others will become unidentifiable after integration.

To calculate ALCS, we used Single Cell Clustering Assessment Framework (SCCAF) ^32^. SCCAF is a self-projection-based approach to assess the validity of a classification system, in our case the cell type annotations. Self-projection is the process in which a machine learning classifier is trained on half of the training set and used to predict the label of the other half. The accuracy of self-projection indicates the clarity of the classification. In our case, we chose to use multinomial logistic classifiers as it has been shown to perform equally well compared with other models in the SCCAF study. We compute the loss of self-projection accuracy on the integrated embedding compared with the original per-species data, as a measurement of how much cell type distinguishability is lost due to integration.

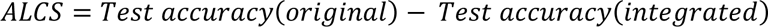

SCCAF models were trained on the PCA embedding for methods returning a pseudo-count matrix and the output embedding for those who output an embedding. It is not applicable for methods that output a kNN graph.

### Aligning ontology annotation with scOntoMatch

ScRNA-seq datasets from different studies often have varying levels of granularity in their cell type annotations, making it difficult to compare them. To address this, we developed an R package called scOntoMatch, which aligns the granularity of cell type annotation. Such alignment requires an ontology as a reference and we used Cell Ontology in this study ^37^. CL describes the ancestor-descendant relationships between cell type terms across animals, making it a reliable reference for harmonising annotation granularity for cross-species annotation.

The scOntoMatch algorithm mainly has two steps. First, it trims the cell type tree in each dataset to remove redundant terms. Second, it identifies an ontology matching across the datasets. The latter is achieved by finding terms that can be directly matched and matching descendants in one dataset to ancestors in another to find the last common ancestor (LCA) term across all datasets. The output is an ontology annotation of all cells that has the highest possible granularity, and aligned across datasets, enabling cross-species comparison. In the aligned annotations, every cell type term is a leaf node in the directed acyclic graph (DAG) created by all cell types from all the input datasets. The scOntoMatch package also provides functions for visualising the CL hierarchy. This approach is essential for batch correction and biology conservation metrics, which rely on cell type annotation and require a systematic, structured way of matching cell type annotations across studies.

### Annotation transfer cross-species

To transfer cell type annotations between species, we employ the SCCAF multinomial logistic classifiers. For each pair of species, we initially train an SCCAF classifier using data from one species on the integrated data. Subsequently, we use this classifier to infer the cell type annotation of the other species within the same integrated result. To evaluate the accuracy of the transferred annotations, we calculate the ARI between the original annotation and the transferred annotation for each species. The overall ARI for each integration strategy is computed by averaging the ARIs of all species in all transfer runs. A successful annotation transfer is indicated by an ARI close to 1.

### Data and codes availability

The BENGAL pipeline is available at https://github.com/Functional-Genomics/BENGAL. Codes and source data for generating the figures in this study are deposed at https://github.com/Functional-Genomics/BENGAL_reproducibility. The package scOntoMatch is available through CRAN https://cran.r-project.org/web/packages/scOntoMatch/index.html, which works on Seurat objects, or https://github.com/Functional-Genomics/scOntoMatch which have two versions that operates on AnnData objects or Seurat objects.

## Acknowledgements

This work was supported by: the European Molecular Biology Laboratory (Y.S., Z.M., A.B., I.P.); the EMBL international PhD program (Y.S.); and the Biotechnology and Biological Sciences Research Council (BBSRC) grant ‘Fly Cell Atlas’ [BB/T014563/1] (I.P.).

## Author contributions

All authors conceived the project. Y.S. designed the framework and curated data, wrote the pipeline and software, performed data analysis and visualization, with supervision from I.P. and inputs from Z.M. <u>and</u> A.B.. Y.S. wrote the initial draft of the manuscript and all authors participated in review and editing.

## Ethics declaration

Competing interests: the authors declare no competing interests.

