## Extended Data for "Benchmarking strategies for cross-species integration of single-cell RNA sequencing data"

Address:

CB10 1SD

United Kingdom

Tel:

+44 (0)1223 494 444

---

[1] Current address: Guangzhou Laboratory, Guangzhou International Bio Island, Guangzhou 510005, China.

**Pancreas_hs_mm**

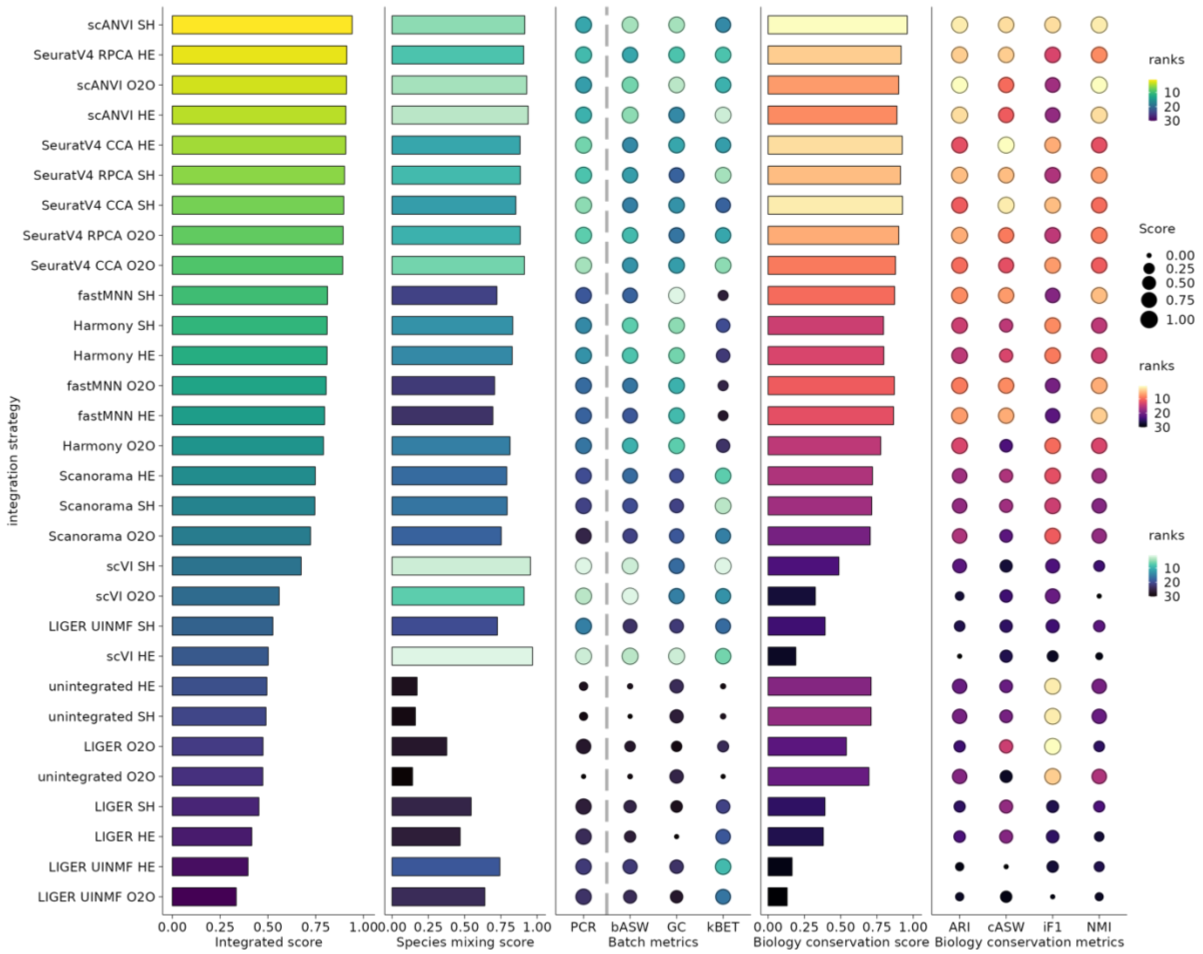

**Hippocampus_hs_mu_ss**

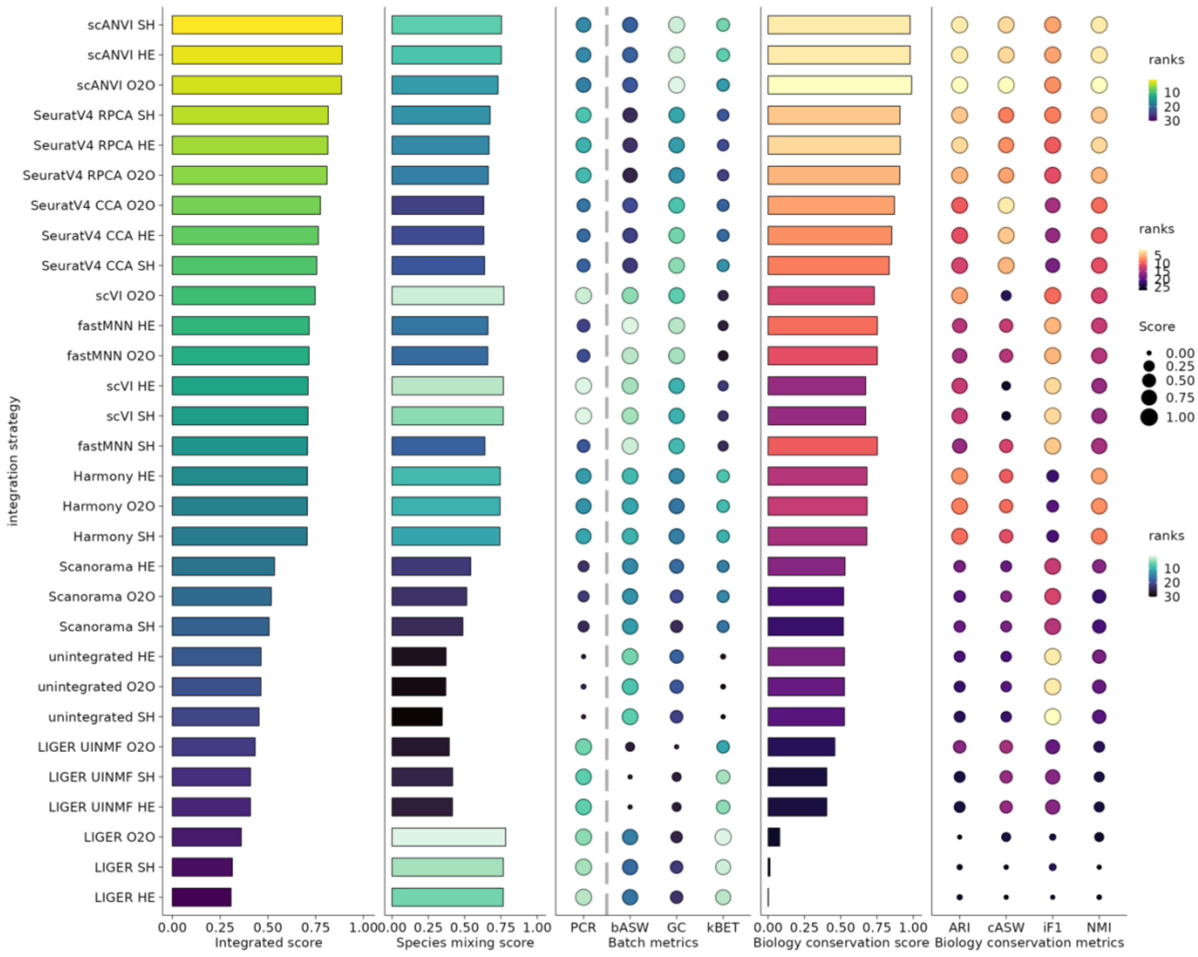

**Embryo_dr_xt**

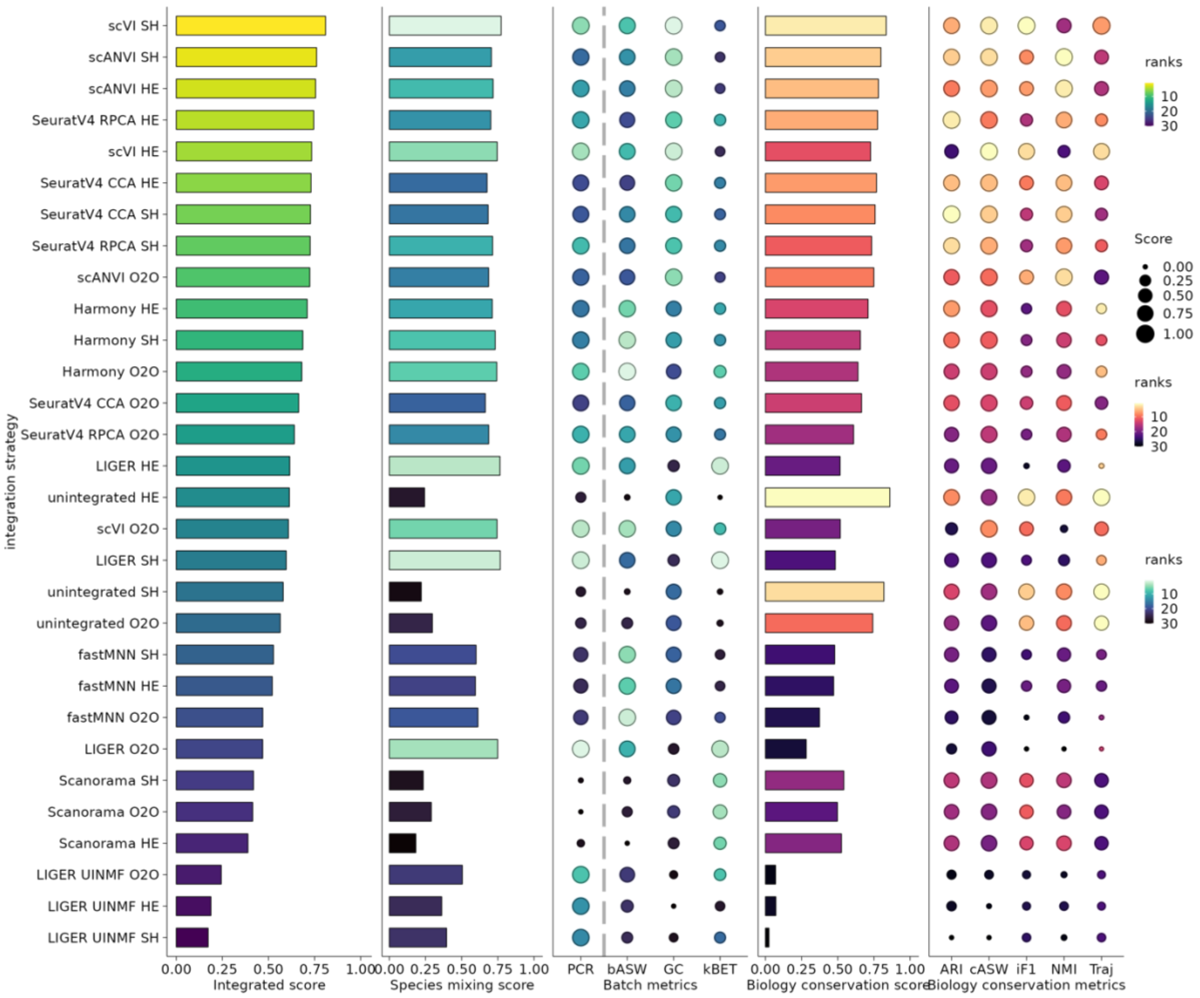

integration strategy
scVI SH
scANVI SH
scANVI HE
SeuratV4 RPCA HE
scVI HE
SeuratV4 CCA HE
SeuratV4 CCA SH
SeuratV4 RPCA SH
scANVI O2O
Harmony HE
Harmony SH
Harmony O2O
SeuratV4 CCA O2O
SeuratV4 RPCA O2O
LIGER HE
unintegrated HE
scVI O2O
LIGER SH
unintegrated SH
unintegrated O2O
fastMNN SH
fastMNN HE
fastMNN O2O
LIGER O2O
Scanorama SH
Scanorama O2O
Scanorama HE
LIGER UINMF O2O
LIGER UINMF HE
LIGER UINMF SH
0.00 0.25 0.50 0.75 1.00
Integrated score
0.00 0.25 0.50 0.75 1.00
Species mixing score
PCR bASW GC kBET
Batch metrics
0.00 0.25 0.50 0.75 1.00
Biology conservation score
ARI cASW iF1 NMI Traj
Biology conservation metrics
ranks
10
20
30
Score
0.00
0.25
0.50
0.75
1.00
ranks
10
20
30
ranks
10
20
30

**Heart_hs_mf**

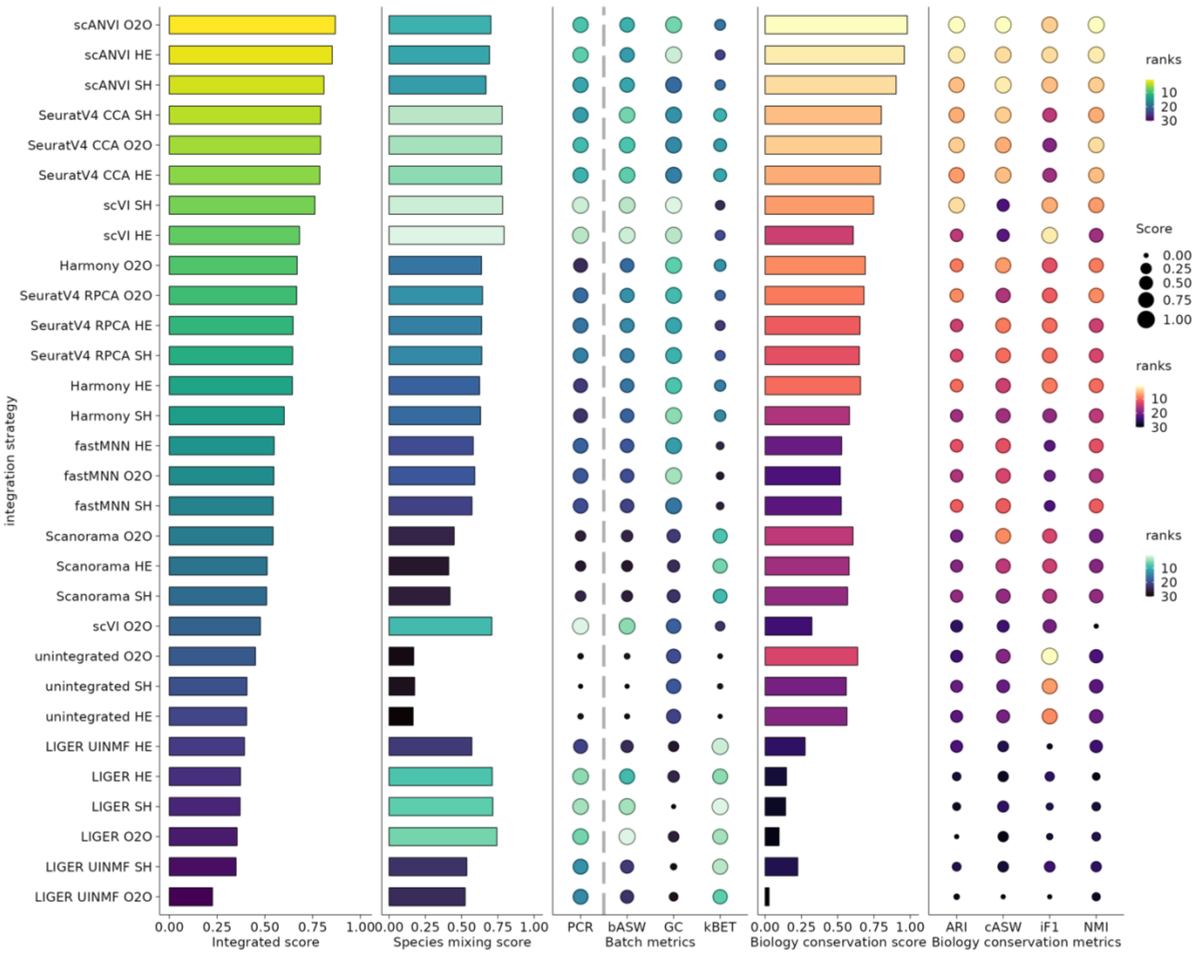

integration strategy
scANVI O2O
scANVI HE
scANVI SH
SeuratV4 CCA SH
SeuratV4 CCA O2O
SeuratV4 CCA HE
scVI SH
scVI HE
Harmony O2O
SeuratV4 RPCA O2O
SeuratV4 RPCA HE
SeuratV4 RPCA SH
Harmony HE
Harmony SH
fastMNN HE
fastMNN O2O
fastMNN SH
Scanorama O2O
Scanorama HE
Scanorama SH
scVI O2O
unintegrated O2O
unintegrated SH
unintegrated HE
LIGER UINMF HE
LIGER HE
LIGER SH
LIGER O2O
LIGER UINMF SH
LIGER UINMF O2O
0.00 0.25 0.50 0.75 1.00
Integrated score
0.00 0.25 0.50 0.75 1.00
Species mixing score
PCR bASW GC kBET
Batch metrics
0.00 0.25 0.50 0.75 1.00
Biology conservation score
ARI cASW iF1 NMI
Biology conservation metrics
ranks
10
20
30
Score
0.00
0.25
0.50
0.75
1.00
ranks
10
20
30
ranks
10
20
30

**Heart_hs_mf_mm**

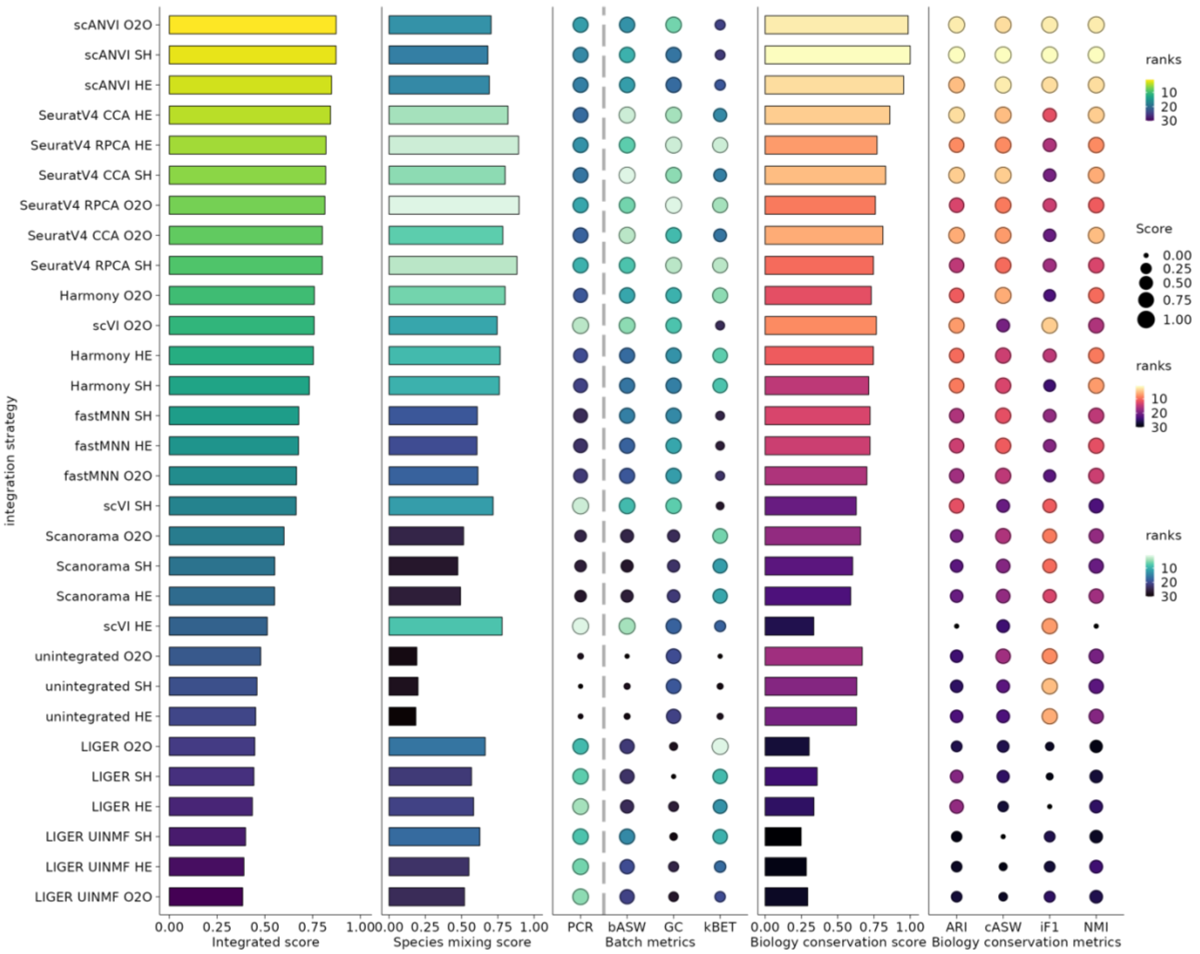

**Heart_hs_mf_mm_xl**

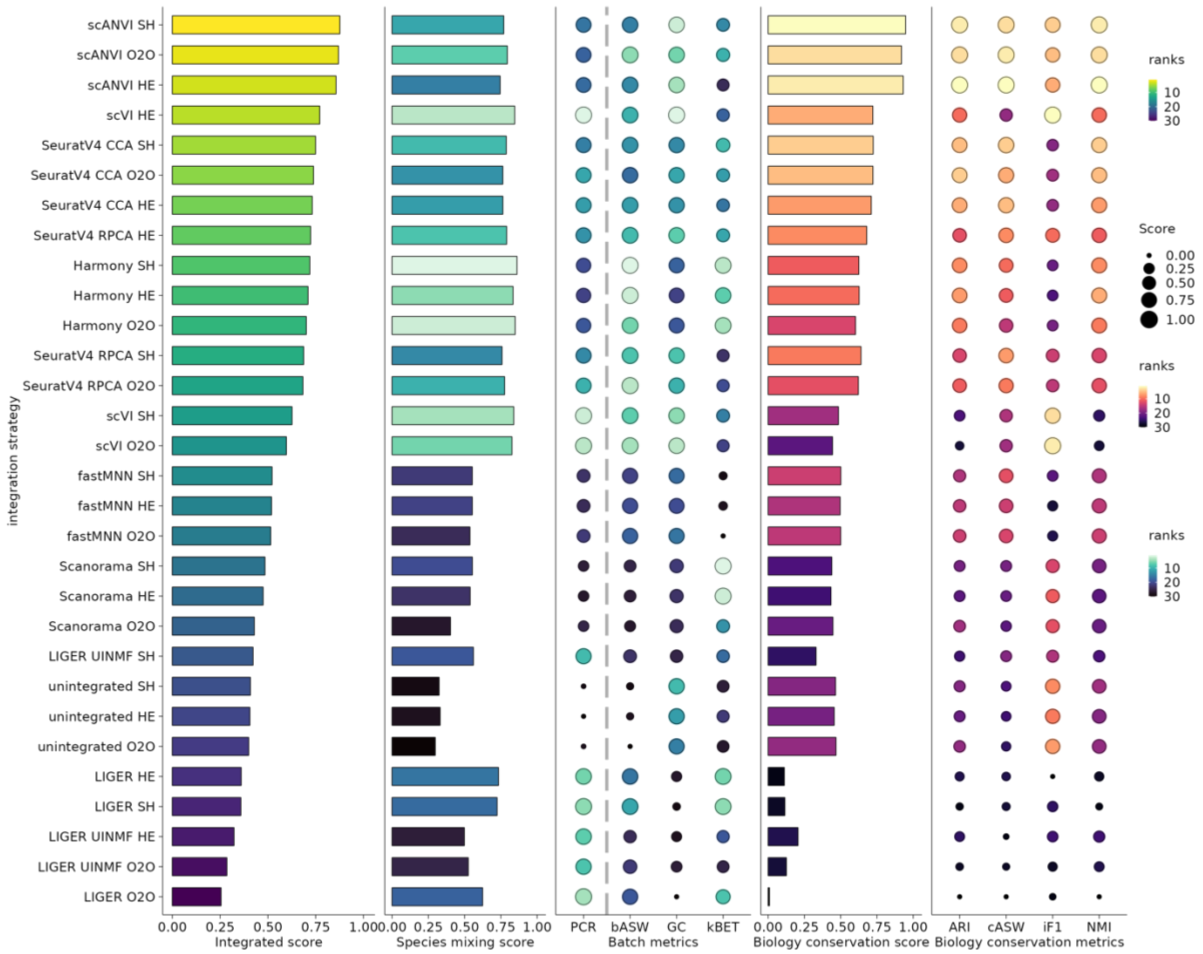

**Heart_hs_mf_mm_xl_dr**

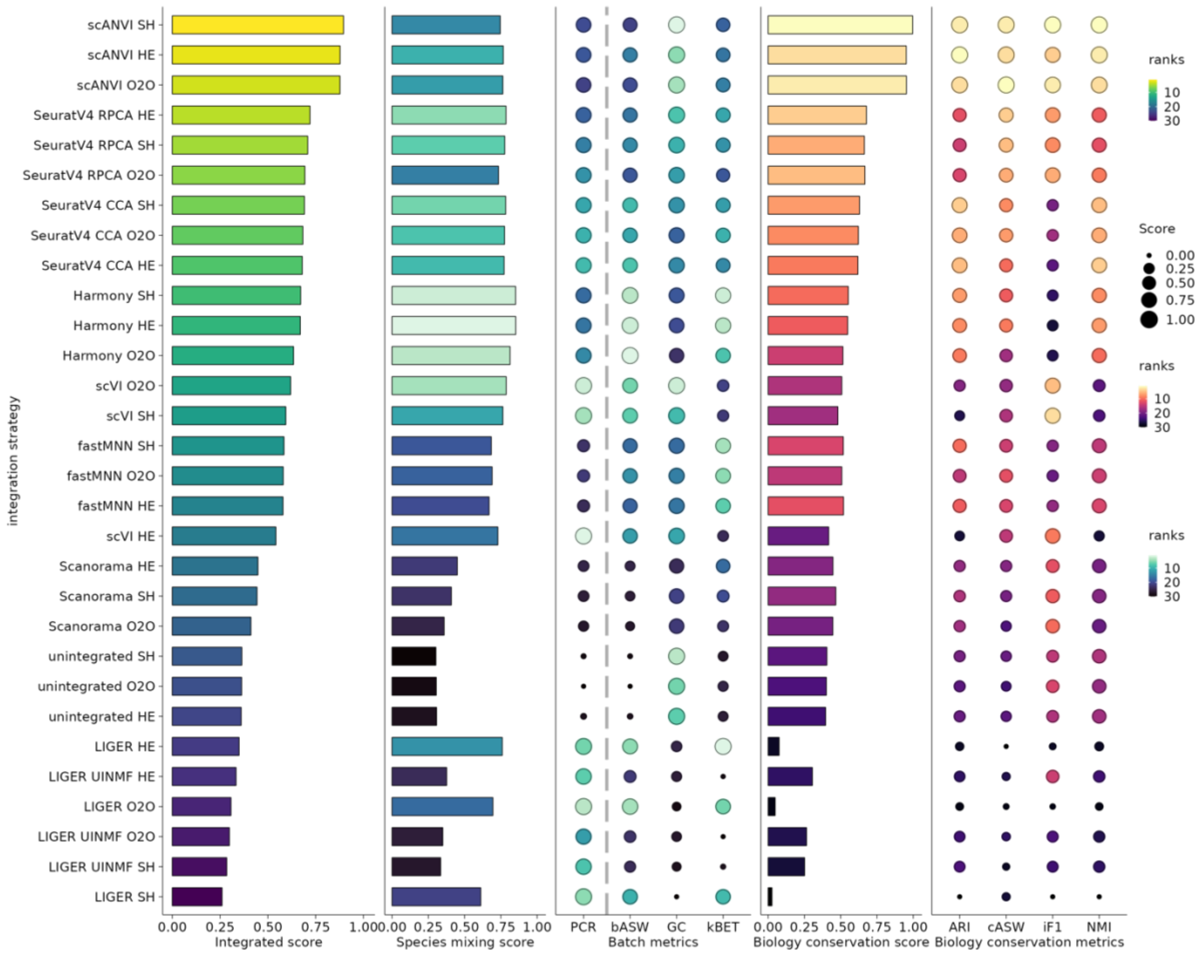

### Heart_hs_mm

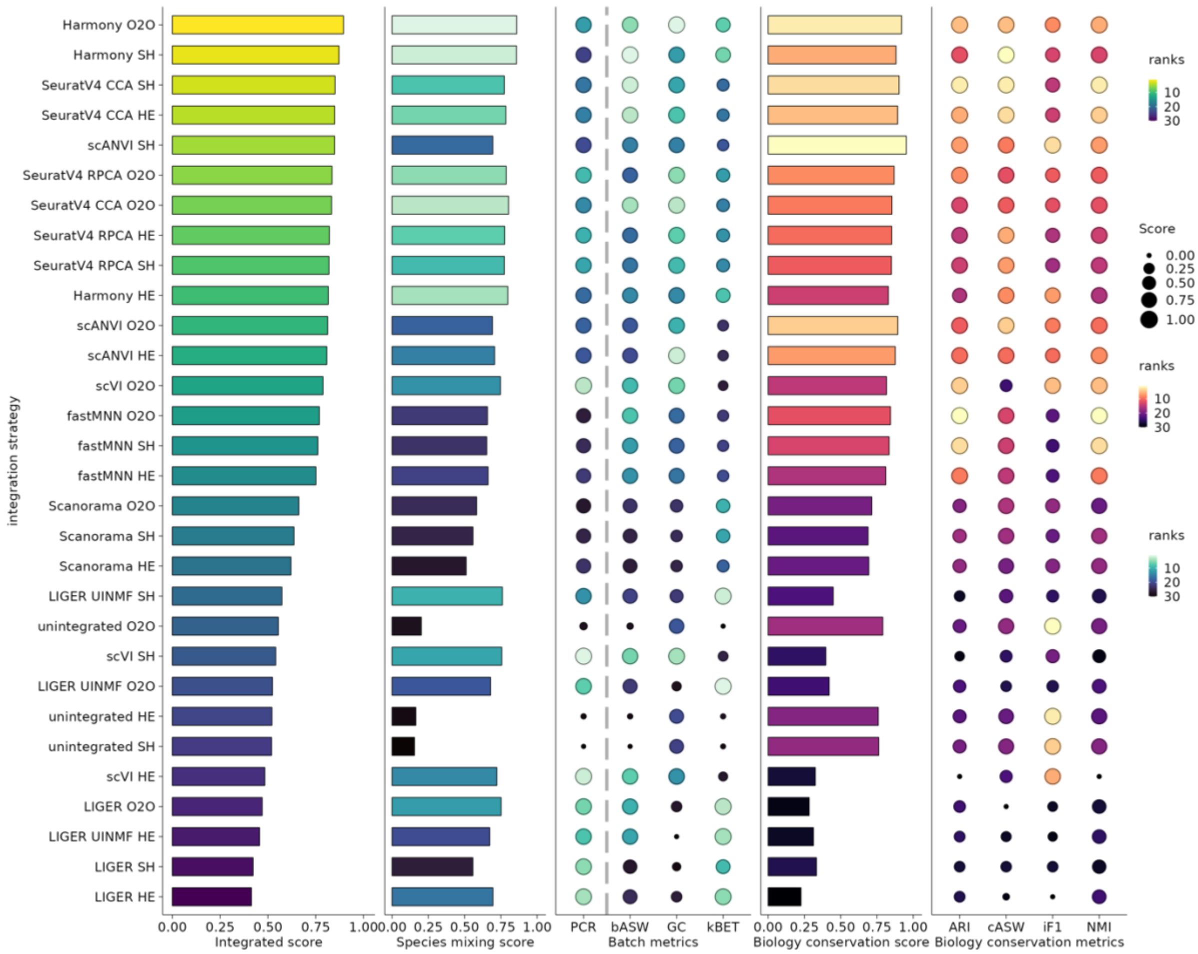

integration strategy
Harmony O2O
Harmony SH
SeuratV4 CCA SH
SeuratV4 CCA HE
scANVI SH
SeuratV4 RPCA O2O
SeuratV4 CCA O2O
SeuratV4 RPCA HE
SeuratV4 RPCA SH
Harmony HE
scANVI O2O
scANVI HE
scVI O2O
fastMNN O2O
fastMNN SH
fastMNN HE
Scanorama O2O
Scanorama SH
Scanorama HE
LIGER UINMF SH
unintegrated O2O
scVI SH
LIGER UINMF O2O
unintegrated HE
unintegrated SH
scVI HE
LIGER O2O
LIGER UINMF HE
LIGER SH
LIGER HE
0.00 0.25 0.50 0.75 1.00
Integrated score
0.00 0.25 0.50 0.75 1.00
Species mixing score
PCR bASW GC kBET
Batch metrics
0.00 0.25 0.50 0.75 1.00
Biology conservation score
ARI cASW iF1 NMI
Biology conservation metrics
ranks
10
20
30
Score
0.00
0.25
0.50
0.75
1.00
ranks
10
20
30
ranks
10
20
30

### Heart_hs_xl

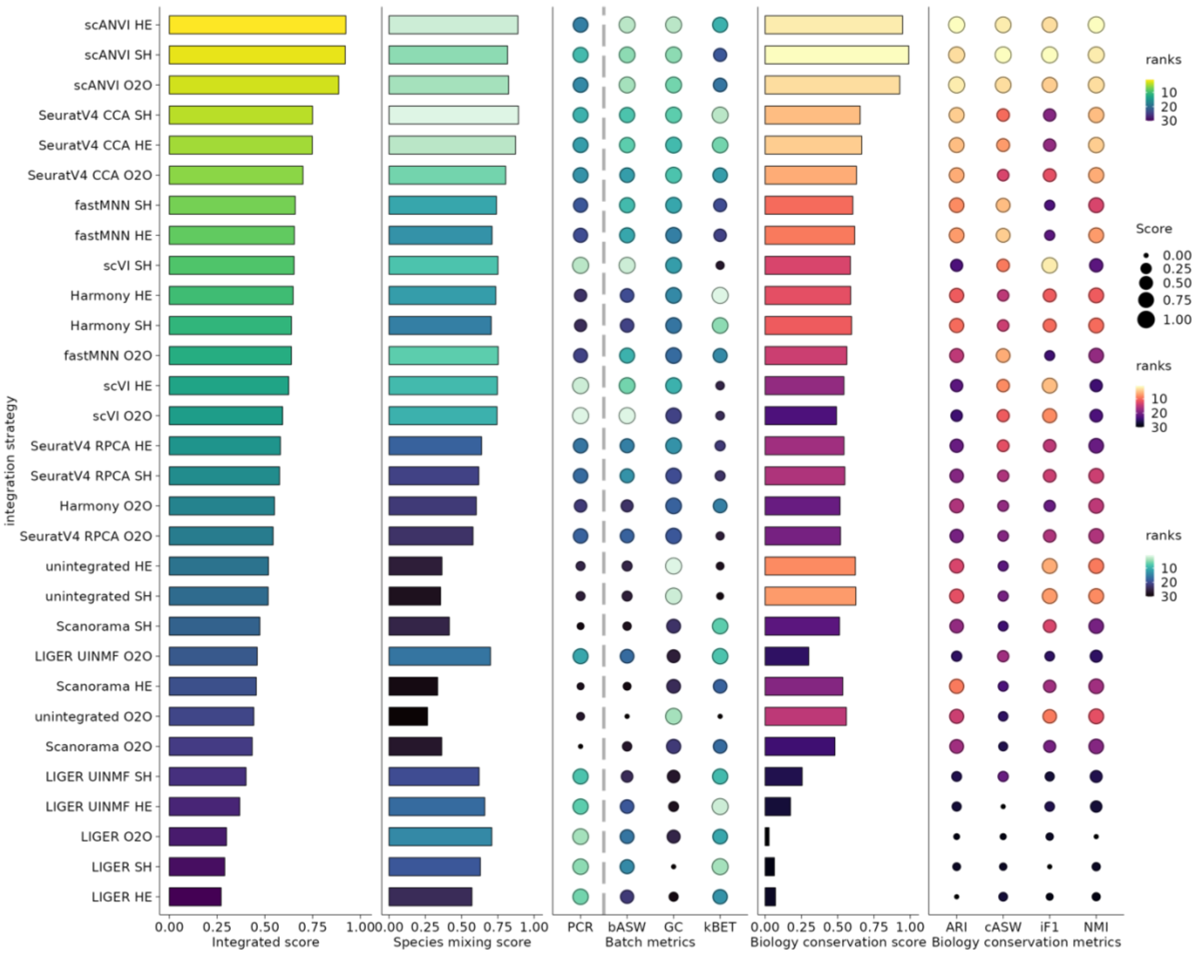

integration strategy
scANVI HE
scANVI SH
scANVI O2O
SeuratV4 CCA SH
SeuratV4 CCA HE
SeuratV4 CCA O2O
fastMNN SH
fastMNN HE
scVI SH
Harmony HE
Harmony SH
fastMNN O2O
scVI HE
scVI O2O
SeuratV4 RPCA HE
SeuratV4 RPCA SH
Harmony O2O
SeuratV4 RPCA O2O
unintegrated HE
unintegrated SH
Scanorama SH
LIGER UINMF O2O
Scanorama HE
unintegrated O2O
Scanorama O2O
LIGER UINMF SH
LIGER UINMF HE
LIGER O2O
LIGER SH
LIGER HE
0.00 0.25 0.50 0.75 1.00
Integrated score
0.00 0.25 0.50 0.75 1.00
Species mixing score
PCR bASW GC kBET
Batch metrics
0.00 0.25 0.50 0.75 1.00
Biology conservation score
ARI cASW iF1 NMI
Biology conservation metrics
ranks
10
20
30
Score
0.00
0.25
0.50
0.75
1.00
ranks
10
20
30
ranks
10
20
30

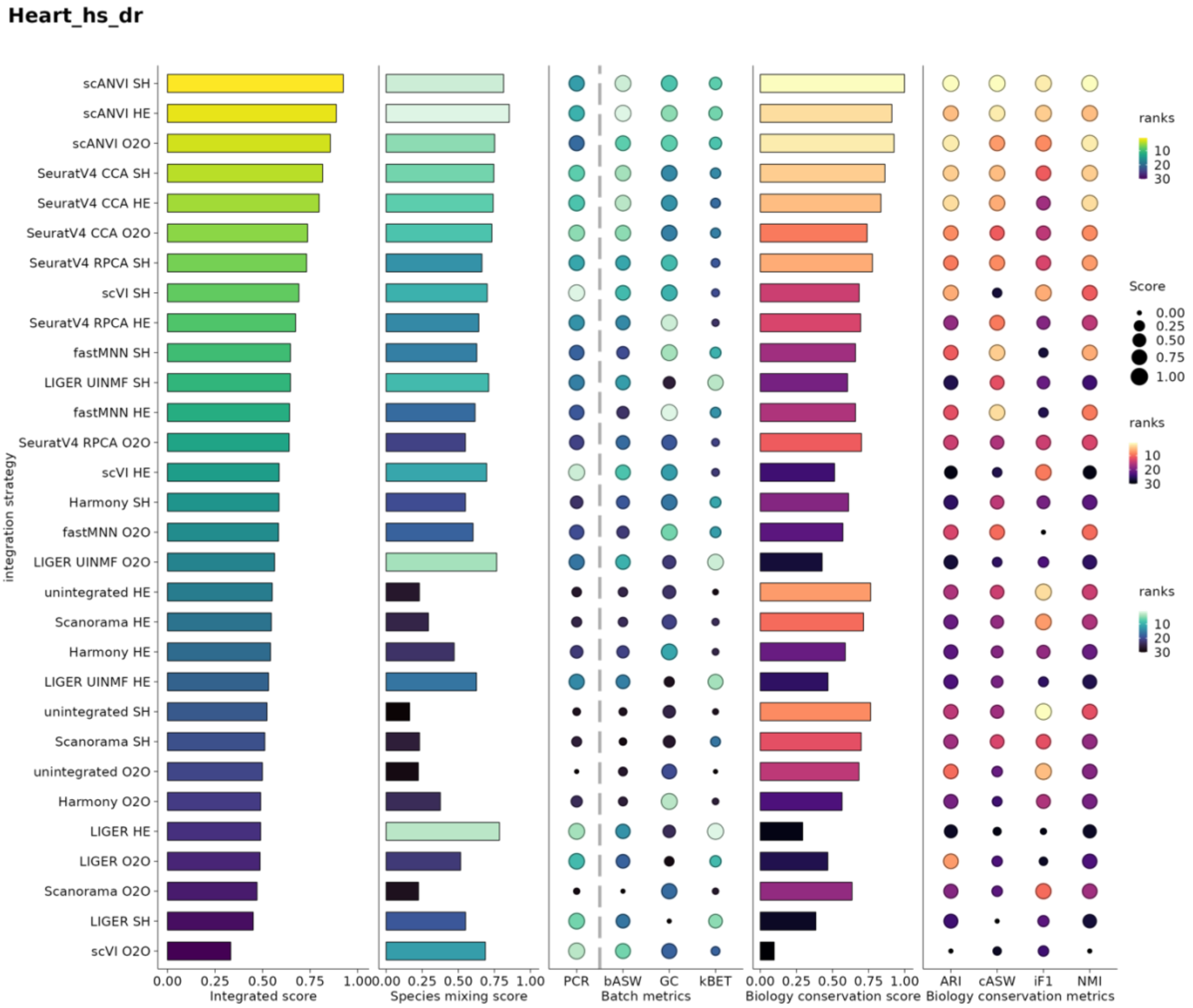
Heart_hs_dr
integration strategy
scANVI SH
scANVI HE
scANVI O2O
SeuratV4 CCA SH
SeuratV4 CCA HE
SeuratV4 CCA O2O
SeuratV4 RPCA SH
scVI SH
SeuratV4 RPCA HE
fastMNN SH
LIGER UINMF SH
fastMNN HE
SeuratV4 RPCA O2O
scVI HE
Harmony SH
fastMNN O2O
LIGER UINMF O2O
unintegrated HE
Scanorama HE
Harmony HE
LIGER UINMF HE
unintegrated SH
Scanorama SH
unintegrated O2O
Harmony O2O
LIGER HE
LIGER O2O
Scanorama O2O
LIGER SH
scVI O2O
0.00 0.25 0.50 0.75 1.00
Integrated score
0.00 0.25 0.50 0.75 1.00
Species mixing score
PCR bASW GC kBET
Batch metrics
0.00 0.25 0.50 0.75 1.00
Biology conservation score
ARI cASW iF1 NMI
Biology conservation metrics
ranks
10
20
30
Score
0.00
0.25
0.50
0.75
1.00
ranks
10
20
30
ranks
10
20
30

**Heart_mf_mm**

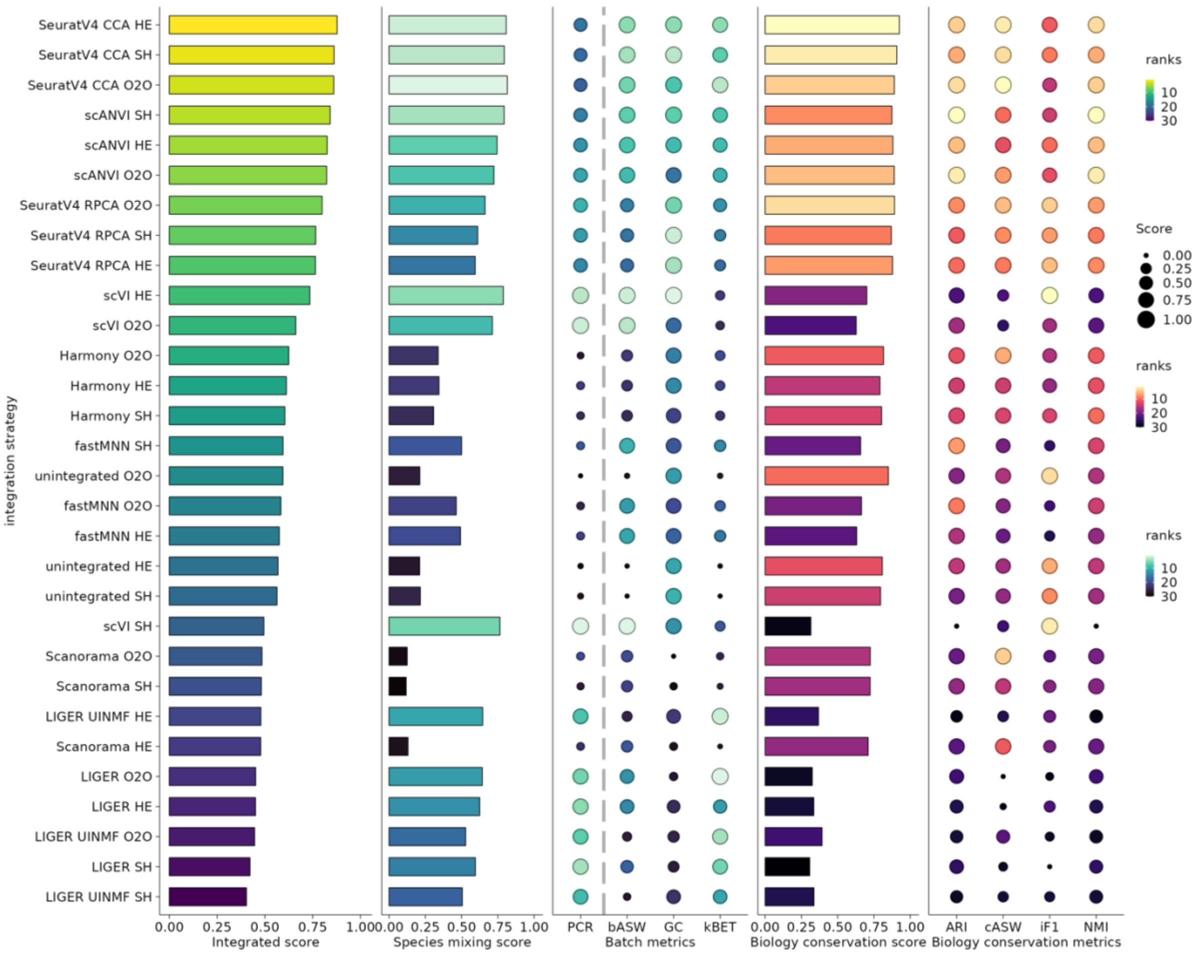
integration strategy
SeuratV4 CCA HE
SeuratV4 CCA SH
SeuratV4 CCA O2O
scANVI SH
scANVI HE
scANVI O2O
SeuratV4 RPCA O2O
SeuratV4 RPCA SH
SeuratV4 RPCA HE
scVI HE
scVI O2O
Harmony O2O
Harmony HE
Harmony SH
fastMNN SH
unintegrated O2O
fastMNN O2O
fastMNN HE
unintegrated HE
unintegrated SH
scVI SH
Scanorama O2O
Scanorama SH
LIGER UINMF HE
Scanorama HE
LIGER O2O
LIGER HE
LIGER UINMF O2O
LIGER SH
LIGER UINMF SH
Integrated score
Species mixing score
PCR
bASW
GC
kBET
Batch metrics
Biology conservation score
ARI
cASW
iF1
NMI
Biology conservation metrics
ranks
Score
0.00
0.25
0.50
0.75
1.00

**Heart_mf_xl**

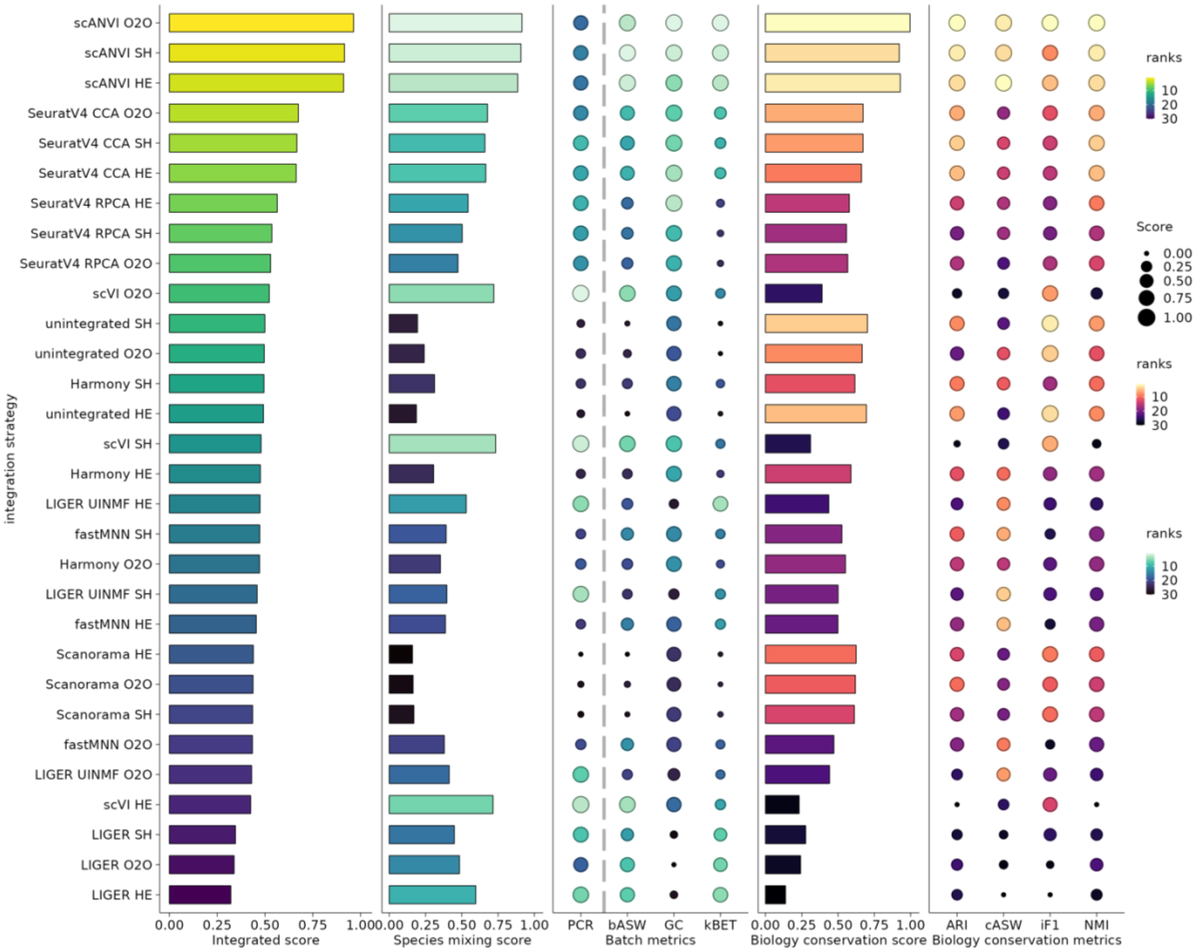

integration strategy
scANVI O2O
scANVI SH
scANVI HE
SeuratV4 CCA O2O
SeuratV4 CCA SH
SeuratV4 CCA HE
SeuratV4 RPCA HE
SeuratV4 RPCA SH
SeuratV4 RPCA O2O
scVI O2O
unintegrated SH
unintegrated O2O
Harmony SH
unintegrated HE
scVI SH
Harmony HE
LIGER UINMF HE
fastMNN SH
Harmony O2O
LIGER UINMF SH
fastMNN HE
Scanorama HE
Scanorama O2O
Scanorama SH
fastMNN O2O
LIGER UINMF O2O
scVI HE
LIGER SH
LIGER O2O
LIGER HE
0.00 0.25 0.50 0.75 1.00
Integrated score
0.00 0.25 0.50 0.75 1.00
Species mixing score
PCR
bASW
GC
kBET
Batch metrics
0.00 0.25 0.50 0.75 1.00
Biology conservation score
ARI
cASW
iF1
NMI
Biology conservation metrics
ranks
10
20
30
Score
0.00
0.25
0.50
0.75
1.00
ranks
10
20
30
ranks
10
20
30

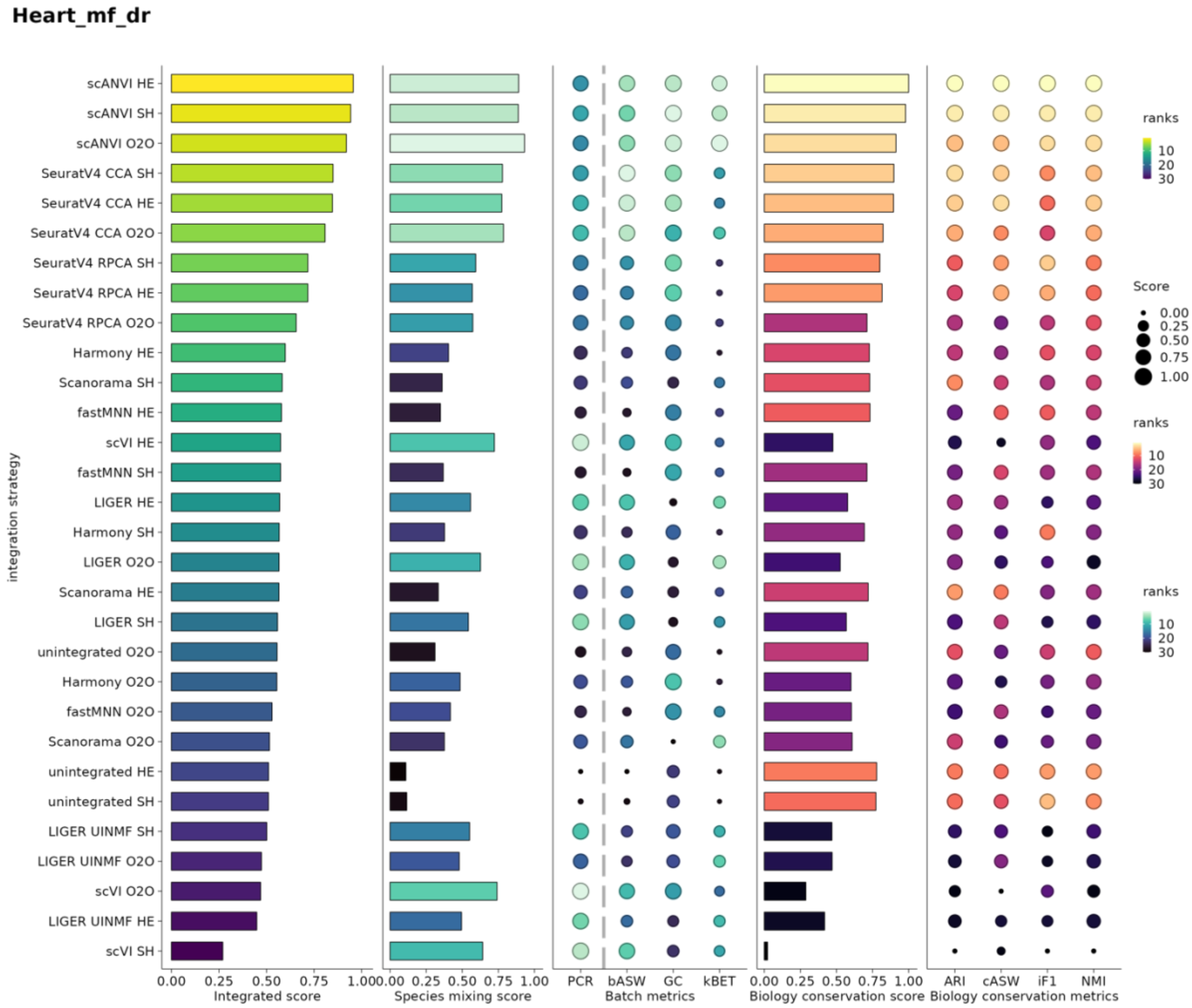

Heart_mf_dr
integration strategy
scANVI HE
scANVI SH
scANVI O2O
SeuratV4 CCA SH
SeuratV4 CCA HE
SeuratV4 CCA O2O
SeuratV4 RPCA SH
SeuratV4 RPCA HE
SeuratV4 RPCA O2O
Harmony HE
Scanorama SH
fastMNN HE
scVI HE
fastMNN SH
LIGER HE
Harmony SH
LIGER O2O
Scanorama HE
LIGER SH
unintegrated O2O
Harmony O2O
fastMNN O2O
Scanorama O2O
unintegrated HE
unintegrated SH
LIGER UINMF SH
LIGER UINMF O2O
scVI O2O
LIGER UINMF HE
scVI SH
0.00 0.25 0.50 0.75 1.00
Integrated score
0.00 0.25 0.50 0.75 1.00
Species mixing score
PCR bASW GC kBET
Batch metrics
0.00 0.25 0.50 0.75 1.00
Biology conservation score
ARI cASW iF1 NMI
Biology conservation metrics
ranks
10
20
30
Score
0.00
0.25
0.50
0.75
1.00
ranks
10
20
30
ranks
10
20
30

**Heart_mm_xl**

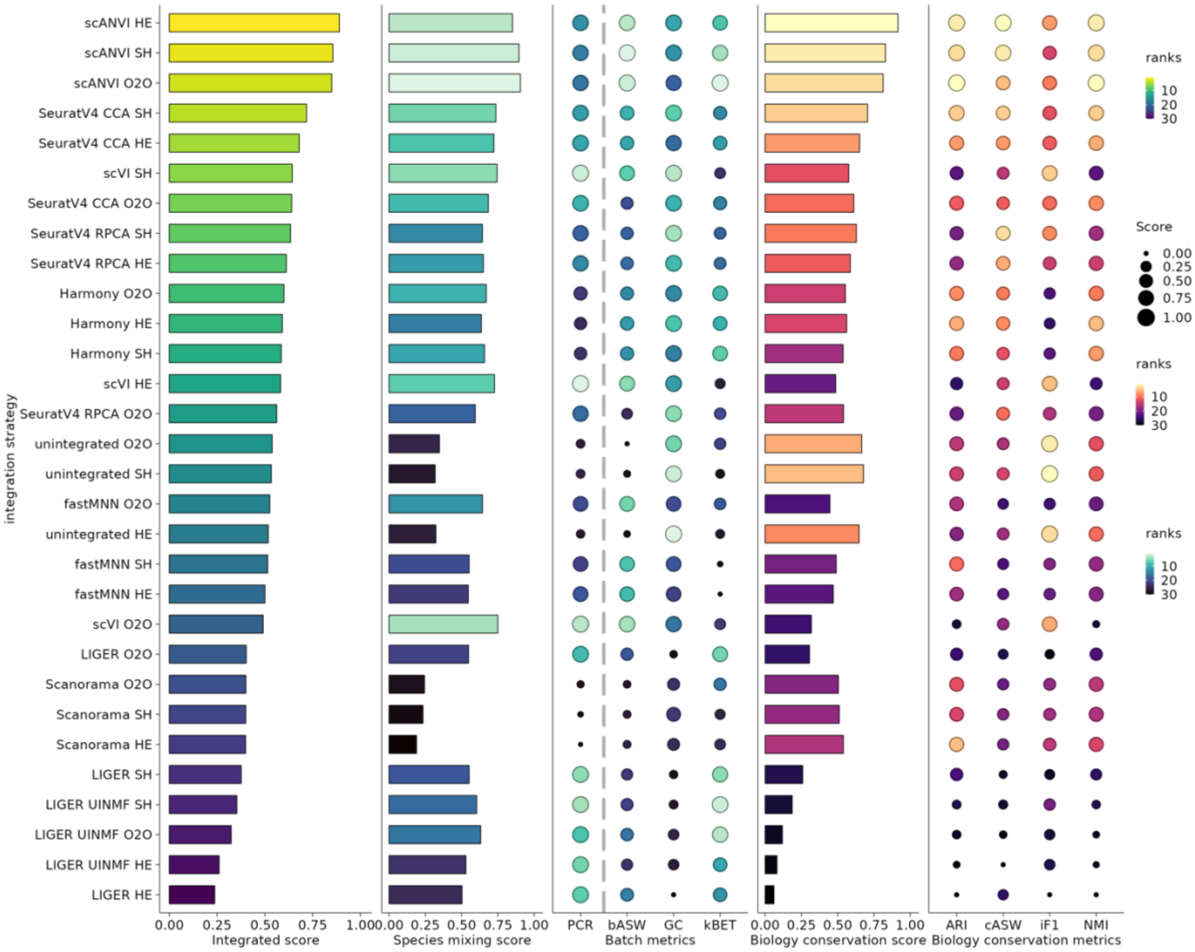

scANVI HE
scANVI SH
scANVI O2O
SeuratV4 CCA SH
SeuratV4 CCA HE
scVI SH
SeuratV4 CCA O2O
SeuratV4 RPCA SH
SeuratV4 RPCA HE
Harmony O2O
Harmony HE
Harmony SH
scVI HE
SeuratV4 RPCA O2O
unintegrated O2O
unintegrated SH
fastMNN O2O
unintegrated HE
fastMNN SH
fastMNN HE
scVI O2O
LIGER O2O
Scanorama O2O
Scanorama SH
Scanorama HE
LIGER SH
LIGER UINMF SH
LIGER UINMF O2O
LIGER UINMF HE
LIGER HE
integration strategy
0.00 0.25 0.50 0.75 1.00
Integrated score
0.00 0.25 0.50 0.75 1.00
Species mixing score
PCR bASW GC kBET
Batch metrics
0.00 0.25 0.50 0.75 1.00
Biology conservation score
ARI cASW iF1 NMI
Biology conservation metrics
ranks
10
20
30
Score
0.00
0.25
0.50
0.75
1.00
ranks
10
20
30
ranks
10
20
30

**Heart_mm_dr**

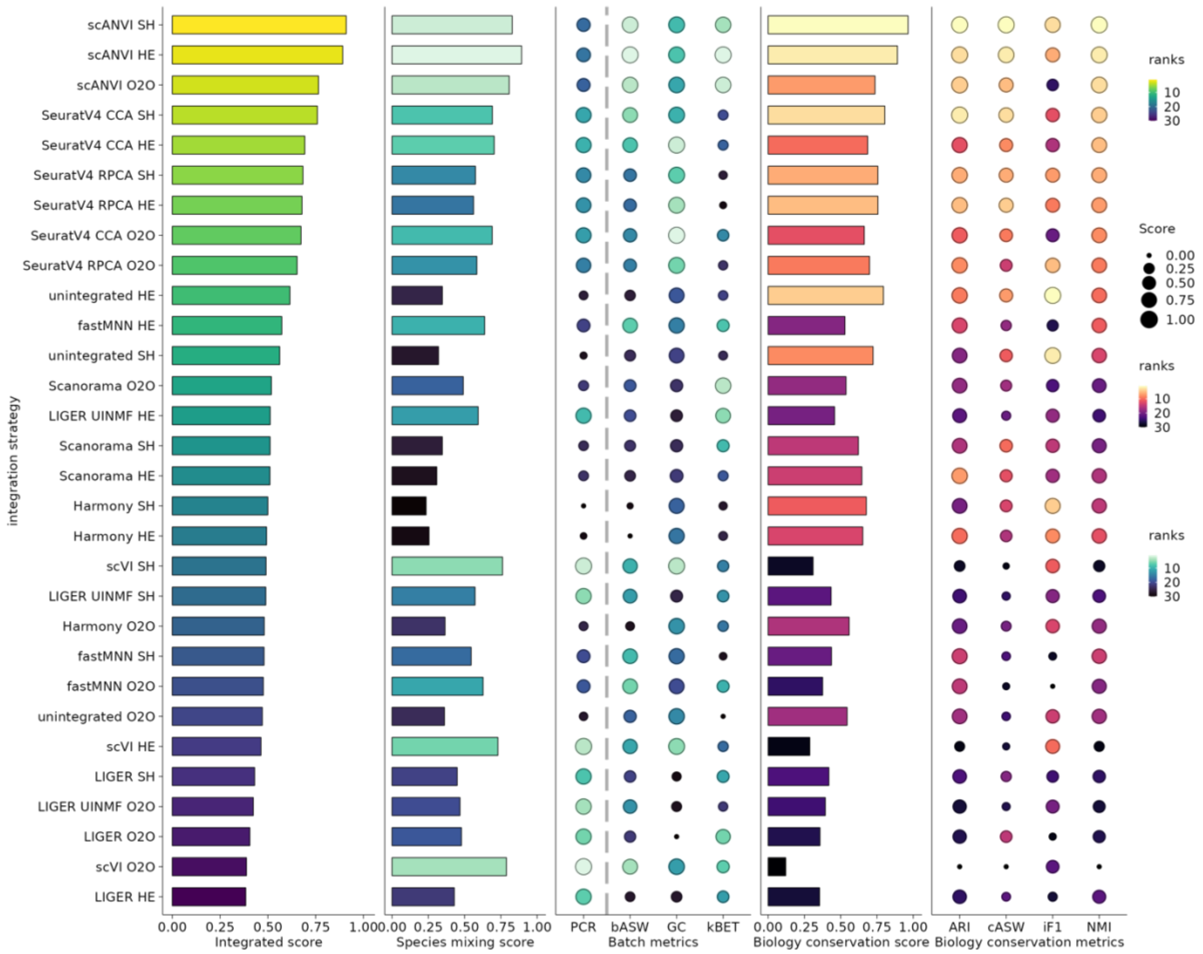
integration strategy
scANVI SH
scANVI HE
scANVI O2O
SeuratV4 CCA SH
SeuratV4 CCA HE
SeuratV4 RPCA SH
SeuratV4 RPCA HE
SeuratV4 CCA O2O
SeuratV4 RPCA O2O
unintegrated HE
fastMNN HE
unintegrated SH
Scanorama O2O
LIGER UINMF HE
Scanorama SH
Scanorama HE
Harmony SH
Harmony HE
scVI SH
LIGER UINMF SH
Harmony O2O
fastMNN SH
fastMNN O2O
unintegrated O2O
scVI HE
LIGER SH
LIGER UINMF O2O
LIGER O2O
scVI O2O
LIGER HE
0.00 0.25 0.50 0.75 1.00
Integrated score
0.00 0.25 0.50 0.75 1.00
Species mixing score
PCR bASW GC kBET
Batch metrics
0.00 0.25 0.50 0.75 1.00
Biology conservation score
ARI cASW iF1 NMI
Biology conservation metrics
ranks
10
20
30
Score
0.00
0.25
0.50
0.75
1.00
ranks
10
20
30
ranks
10
20
30

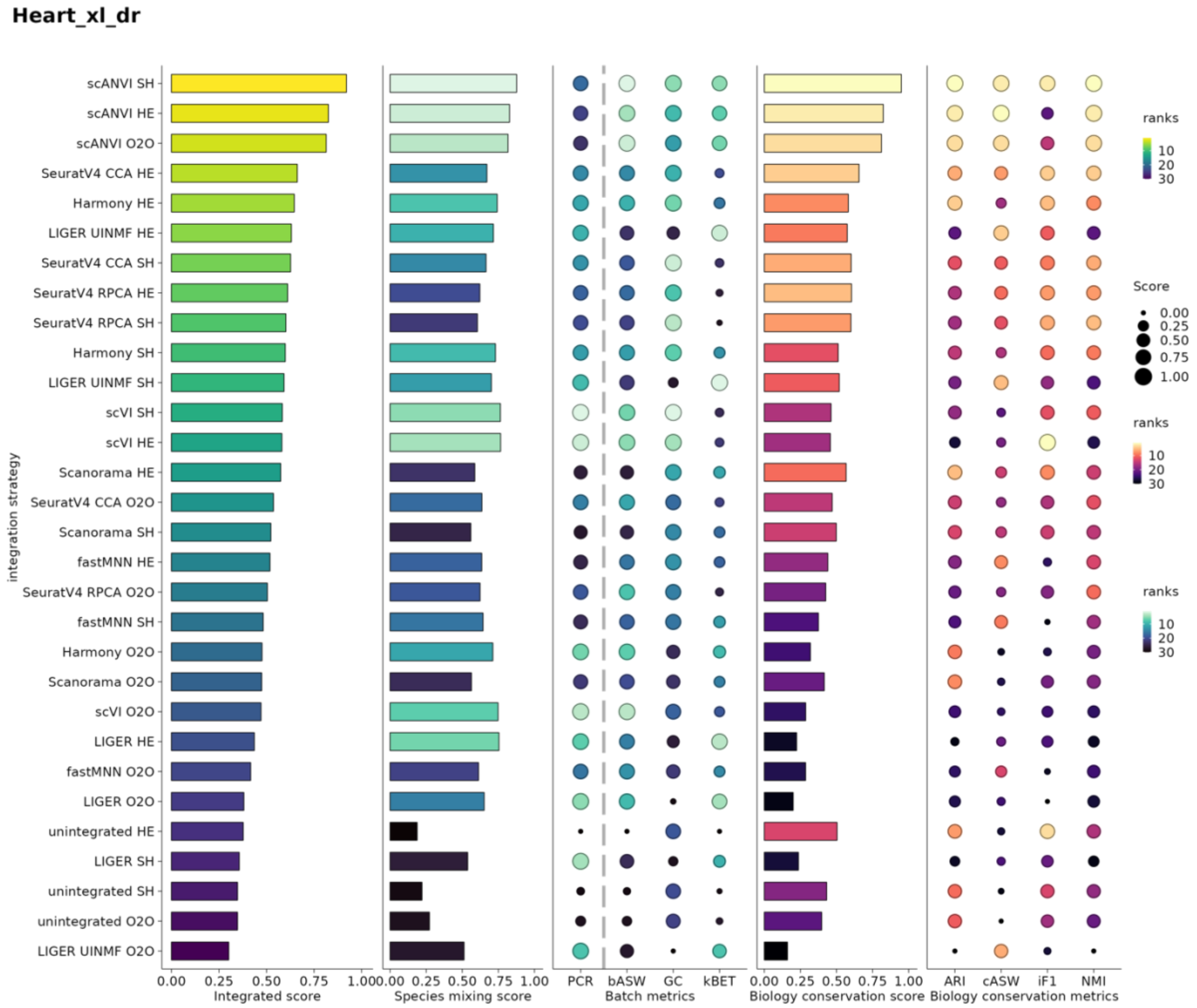


**Extended Data Figure 1-16: Per-task benchmarking scores and metrics of different integration strategies on cross-species analysis.** Batch removal metrics and biological conservation metrics are min-max scaled per task. The species mixing score is the average of 4 batch removal metrics, while the biological conservation score is the average of 5 biology conservation metrics. The integrated score is a weighted average of species mixing score and biology conservation score with 40/60 weighting. The trajectory conservation metric is only calculated for the Embryo_dr_xt task because it is only relevant to that task. Grey dash lines indicate a distinction between metrics that do not rely on cell type annotation (PCR) with metrics relying on cell type annotation. O2O, only uses one-to-one orthologs; HE, one-to-one orthologs plus one-to-many and many-to-many orthologs matched by higher average expression level; SH, one-to-one orthologs plus one-to-many and many-to-many orthologs matched by stronger homology confidence; PCR: principal component regression; bASW:

batch average silhouette width, GC: graph connectivity, kBET: k-nearest neighbour batch effect test, ARI: adjusted rand index, cASW: cell type average silhouette width, NMI: normalised mutual information of cell label, iF1: isolated label F1 score, Traj: trajectory conservation.

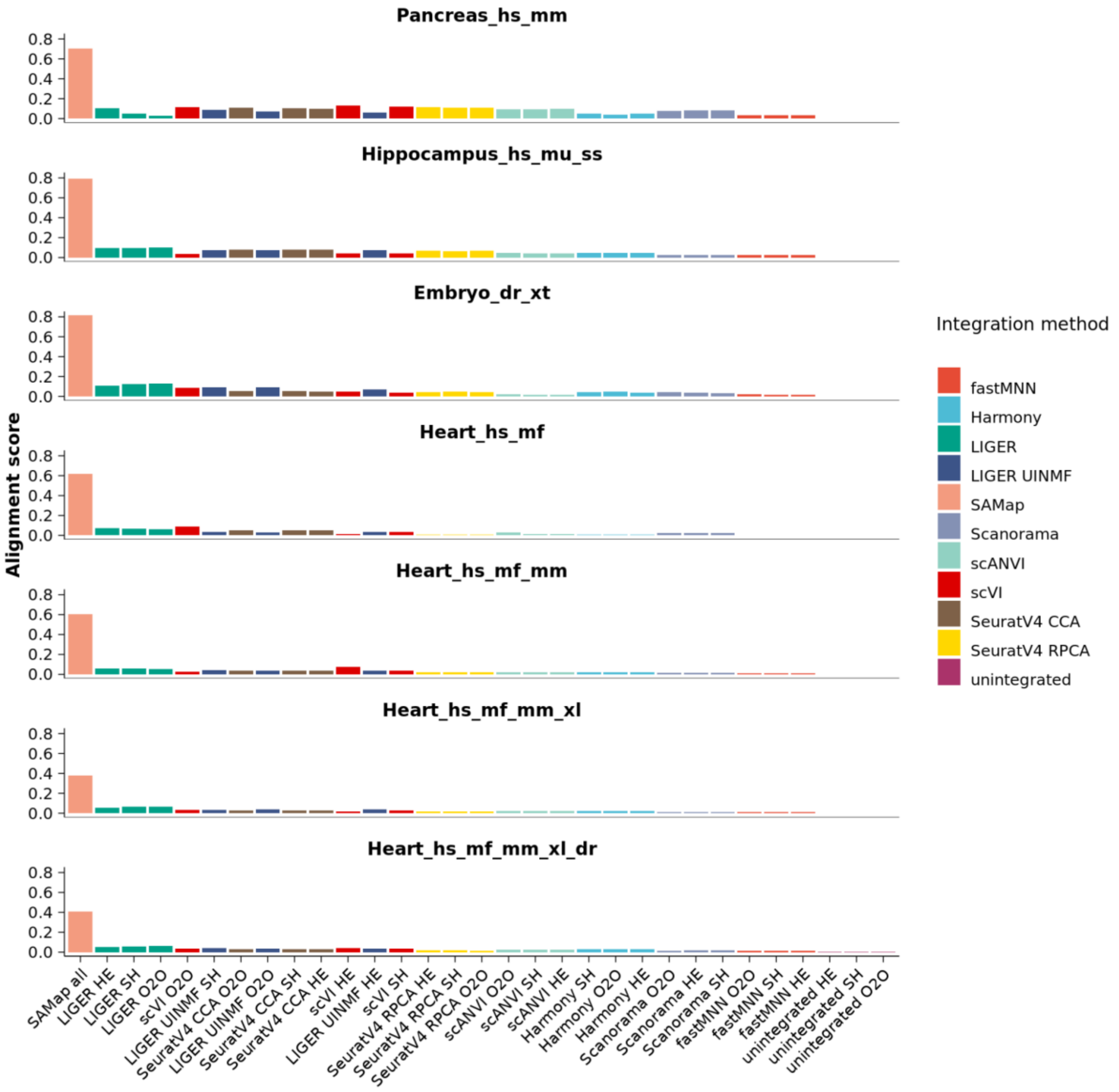


**Extended Data Figure 17: Alignment score for all strategies in 7 reference integration tasks.** The alignment score is the average number of cross-species neighbours as a percentage of the maximum number of neighbours (k=20 is used in this study, see Methods for score details). O2O, only uses one-to-one orthologs; HE, one-to-one orthologs plus one-to-many and many-to-many orthologs matched by higher average expression level; SH, one-to-one orthologs plus one-to-many and many-to-many orthologs matched by stronger homology confidence.

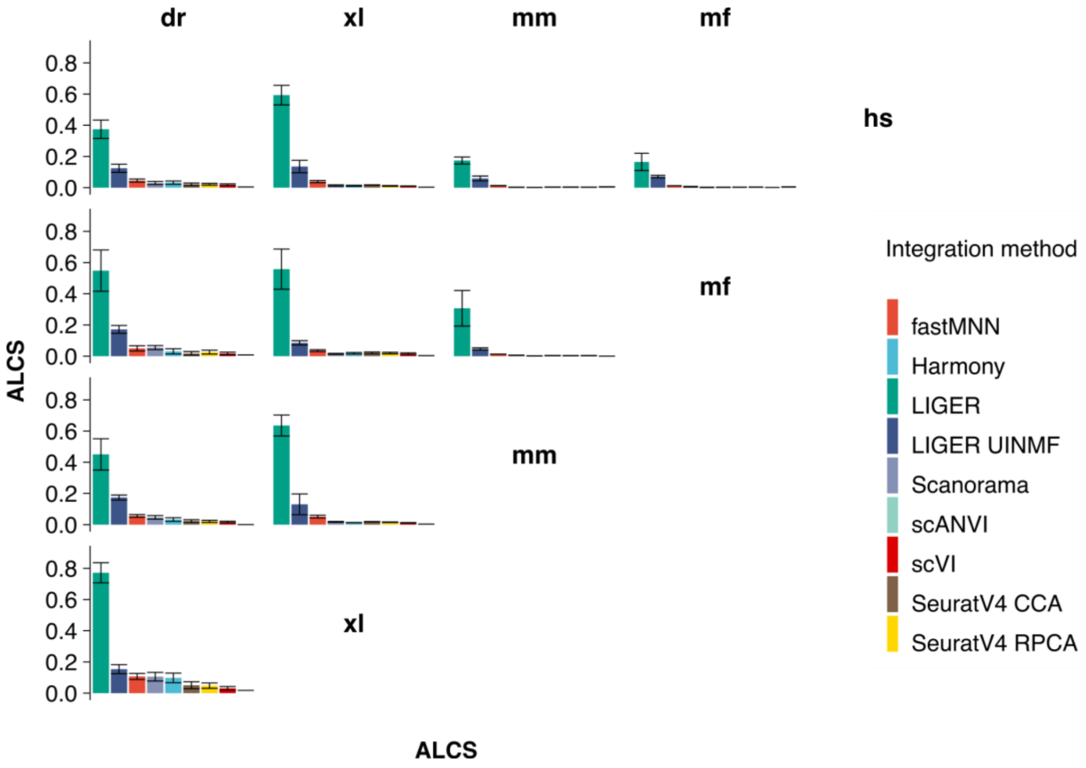


**Extended Data Figure 18: ALCS for all integration strategies in pairwise integration of the heart task.** Bars represent average ALCS across three homology strategies for each integration method and error bars indicate SD. This plot is symmetric with the top right to the bottom left as the axis. ALCS: accuracy loss of cell type self-projection, hs: *Homo sapiens*, mf: *Macaca fascicularis*, mm: *Mus musculus*; xl, *Xenopus laevis*; dr, *Danio rerio*.

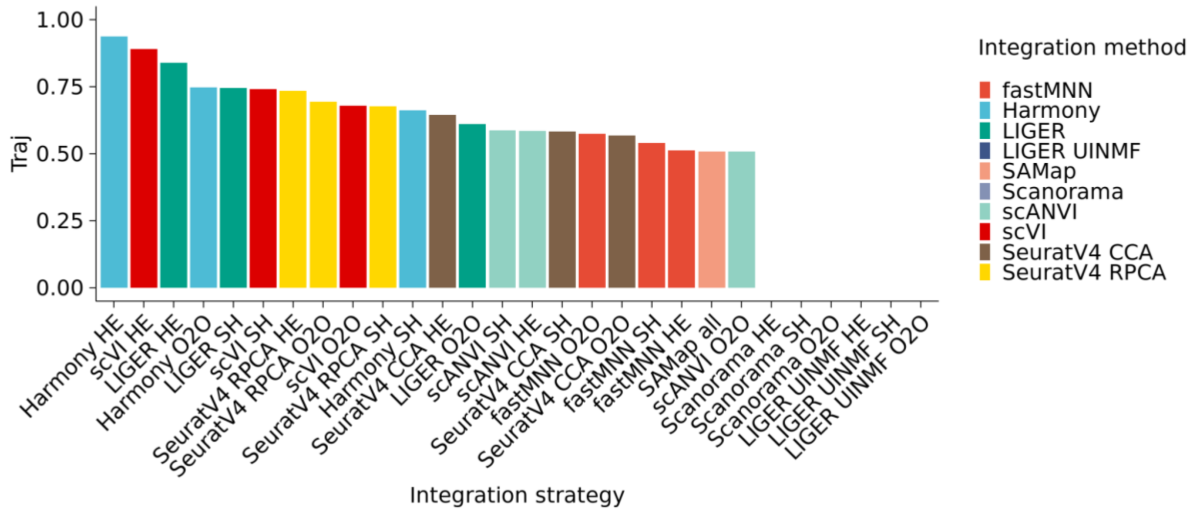


**Extended Data Figure 19: Trajectory conservation score in the Embryo_dr_xt task of all strategies.** The trajectory conservation score compares the percentage of cells with conserved dpt trajectory with unintegrated data on integrated output. Traj, trajectory conservation score; O2O, only use one-to-one orthologs; HE, one-to-one orthologs plus one-to-many and many-to-many orthologs matched by higher average expression level; SH, one-to-one orthologs plus one-to-many and many-to-many orthologs matched by stronger homology confidence.

**H. sapiens**

Cell type tree of original annotation

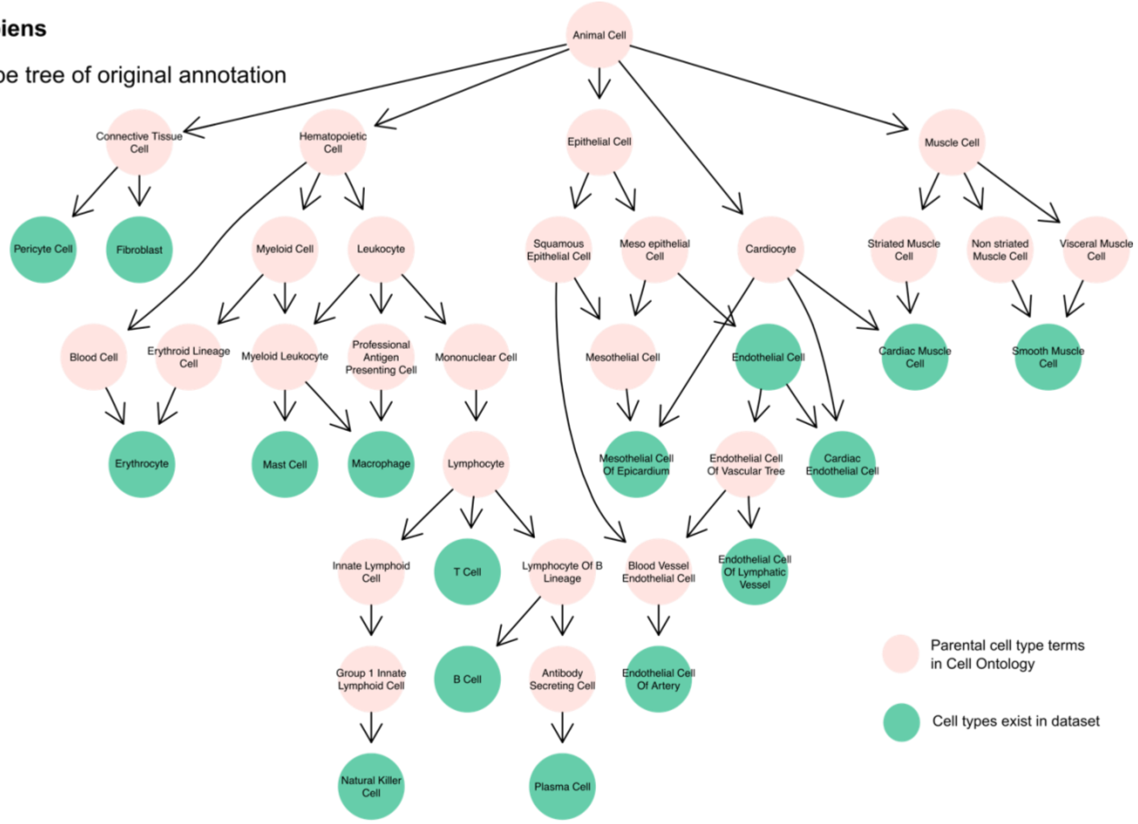


Cell type tree of mapped annotation across five species

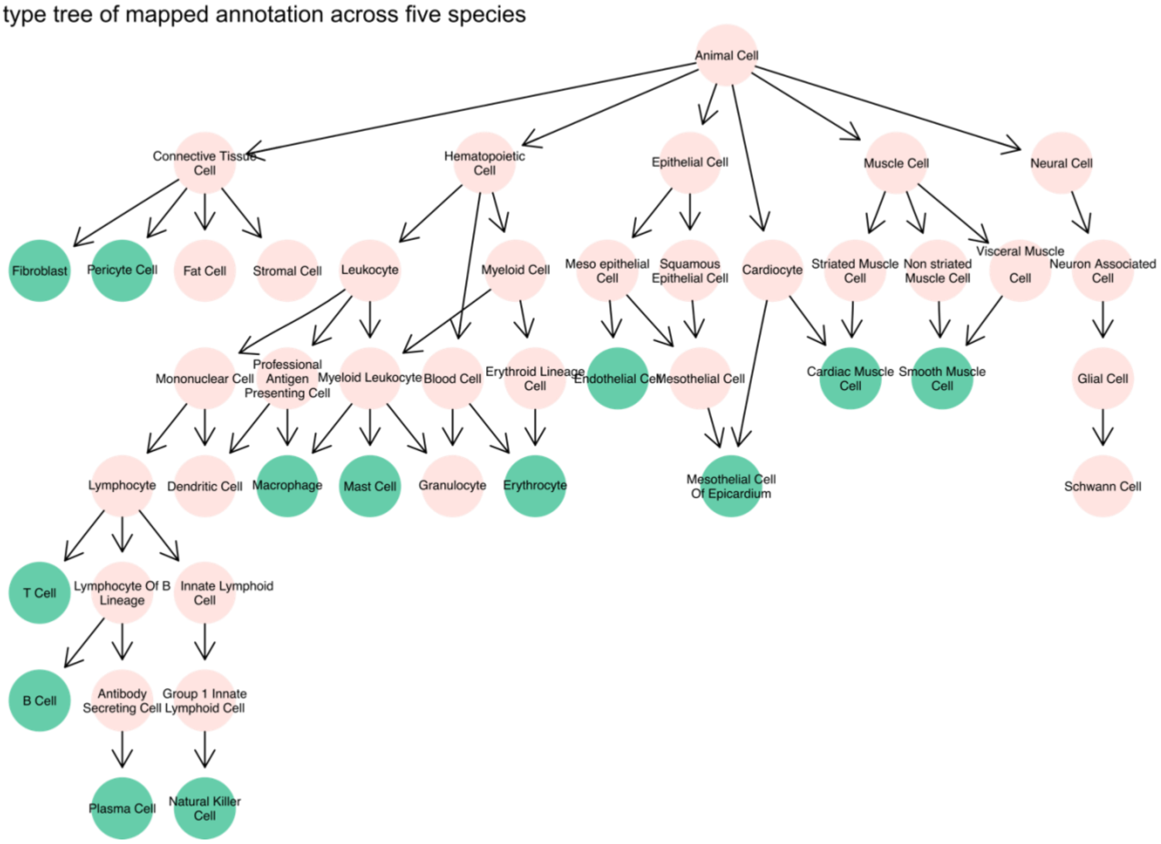

**M. fascicularis**

Cell type tree of original annotation

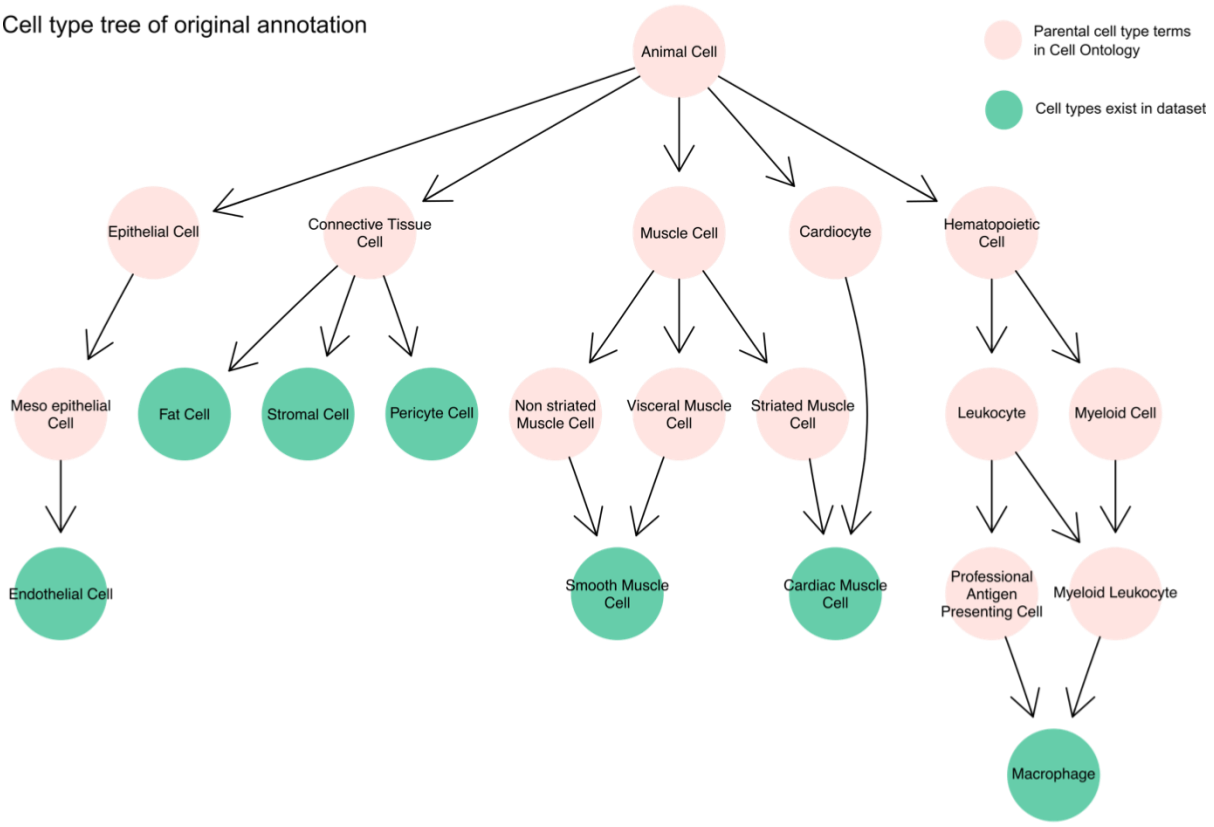


Cell type tree of mapped annotation across five species

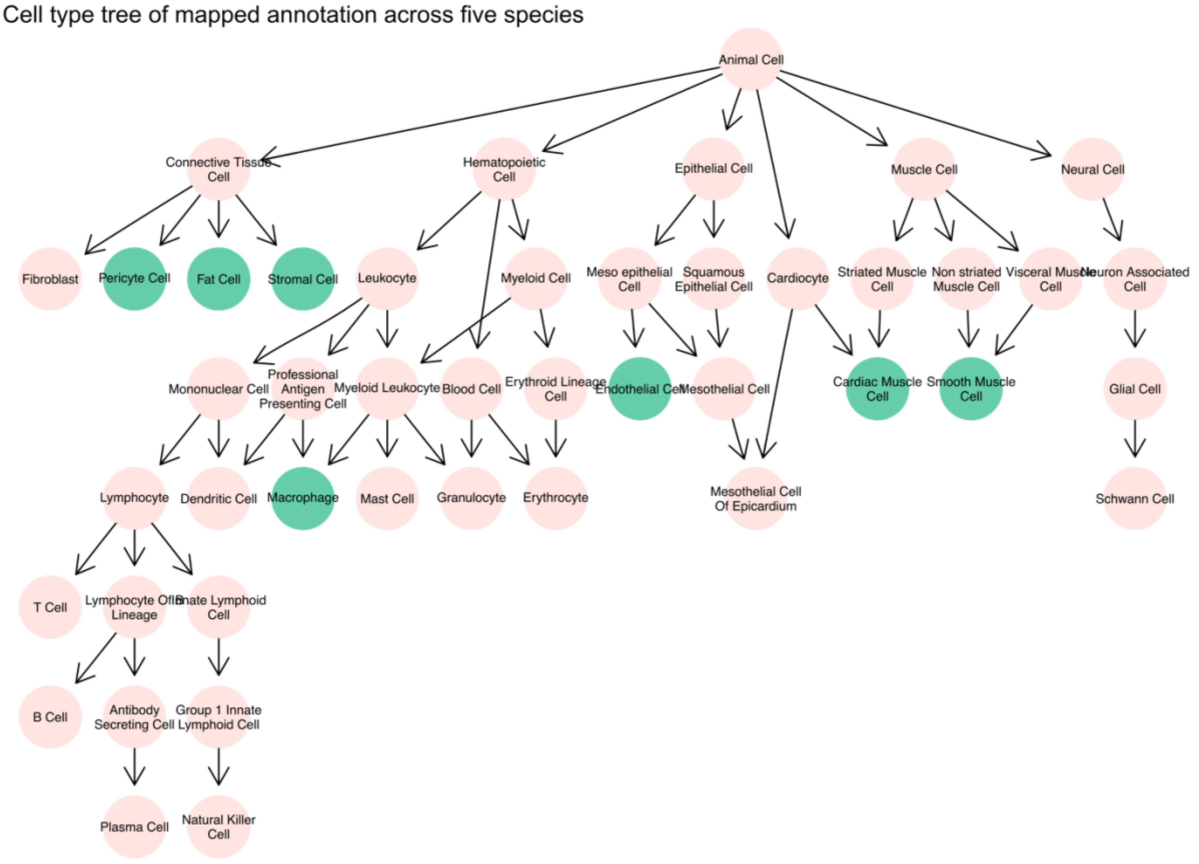

**M. musculus**

Cell type tree of original annotation

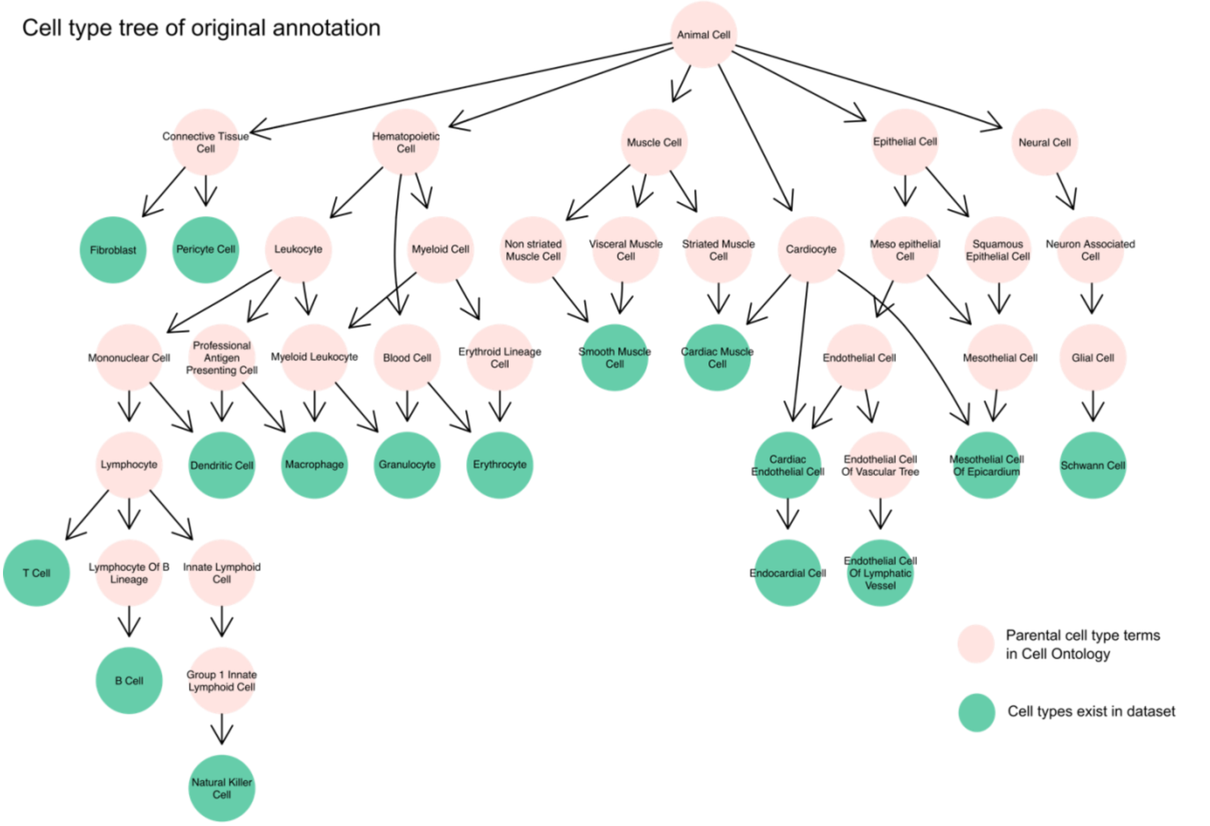


Cell type tree of mapped annotation across five species

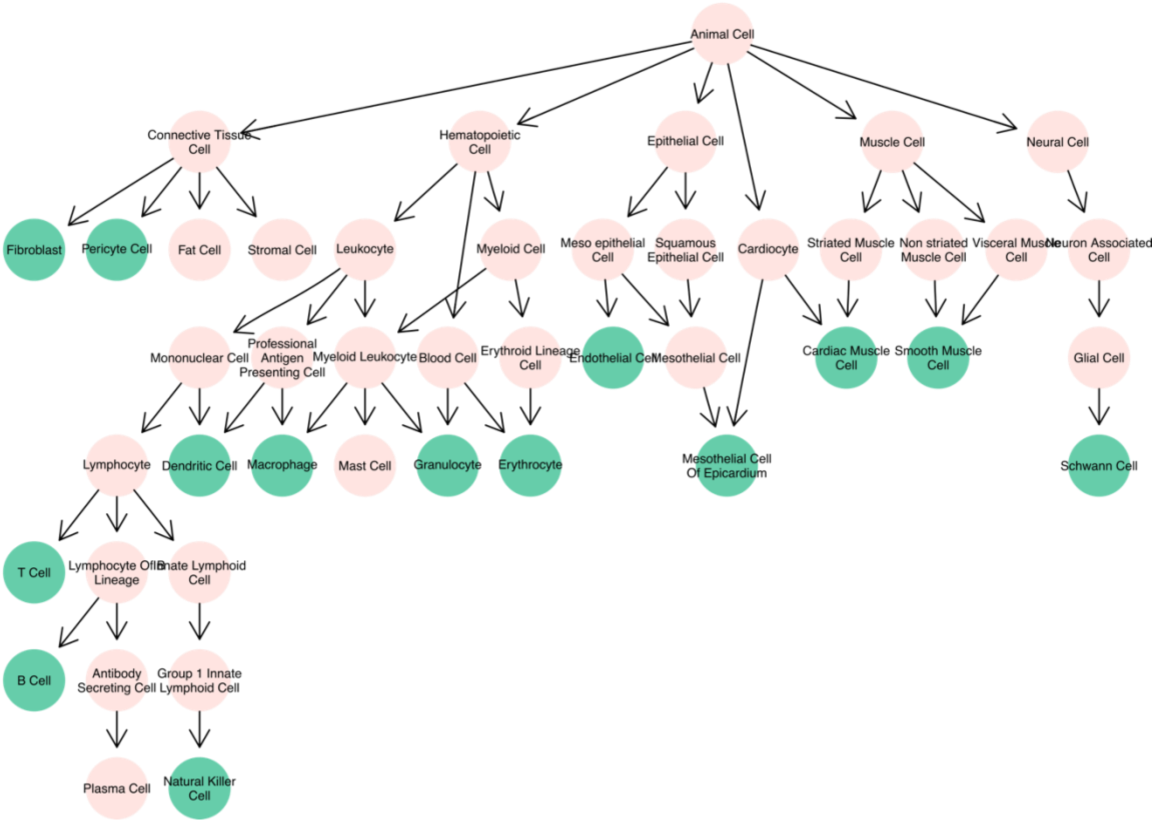

**X. laevis**

Cell type tree of original annotation

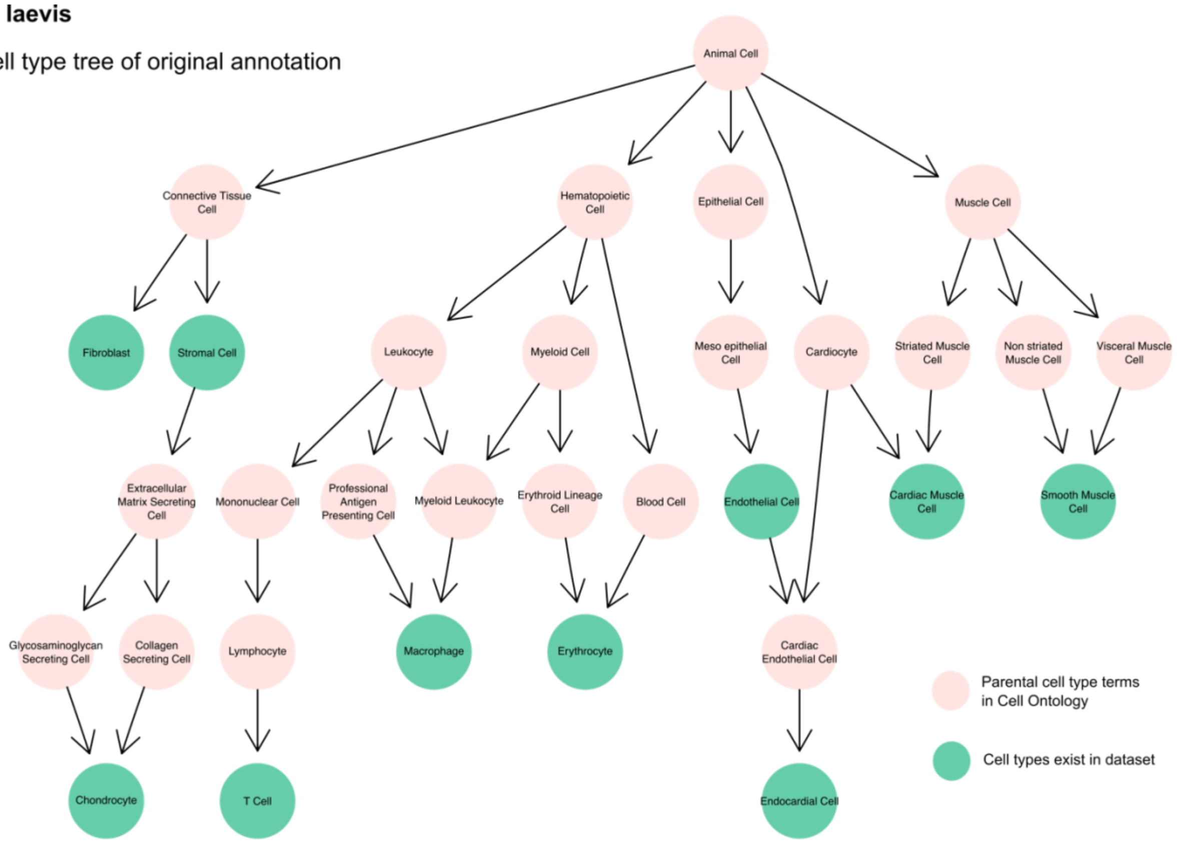


Cell type tree of mapped annotation across five species

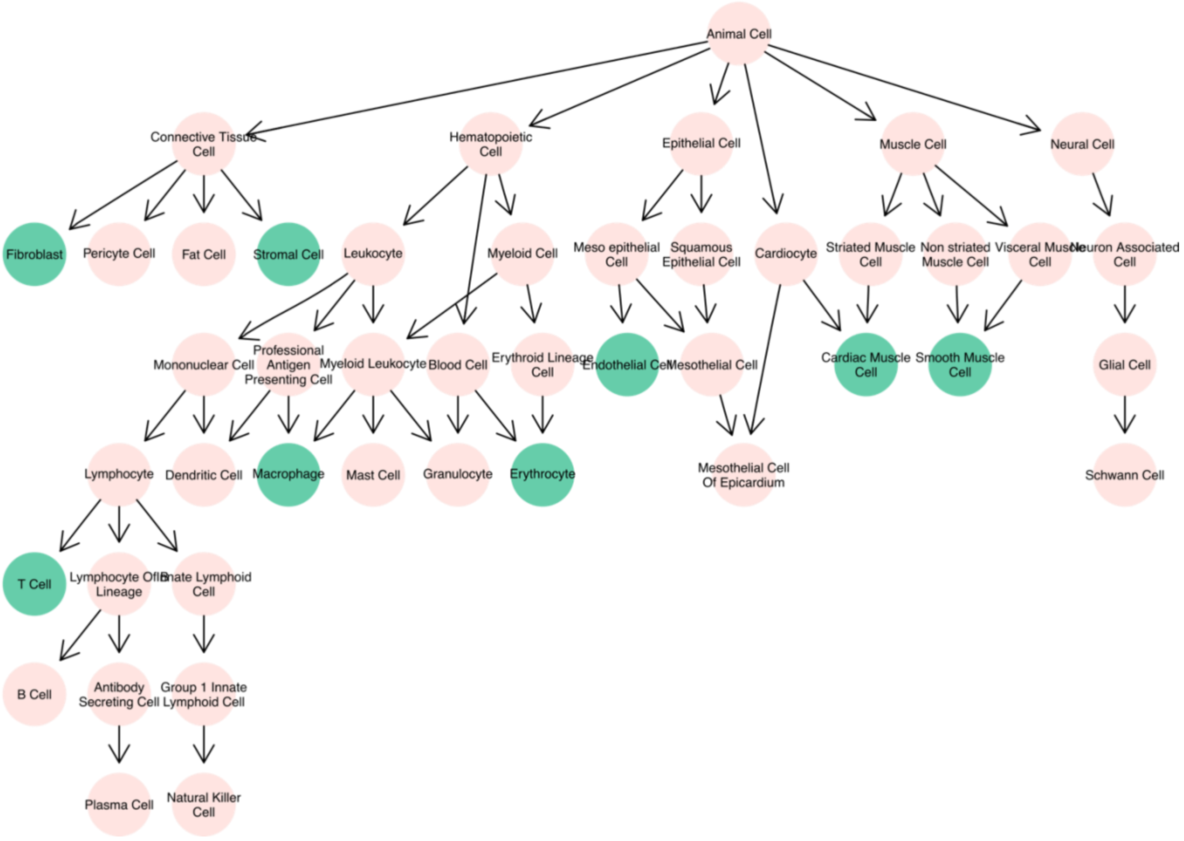

***D. rerio***

Cell type tree of original annotation

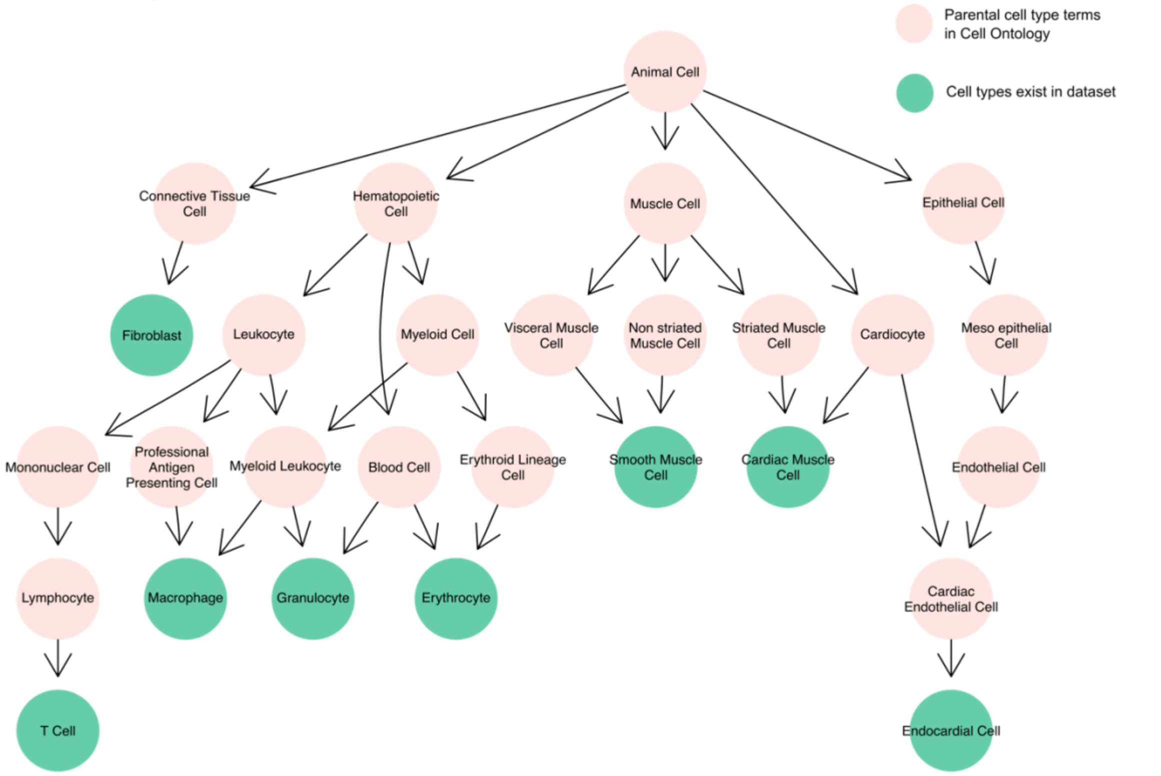


Cell type tree of mapped annotation across five species

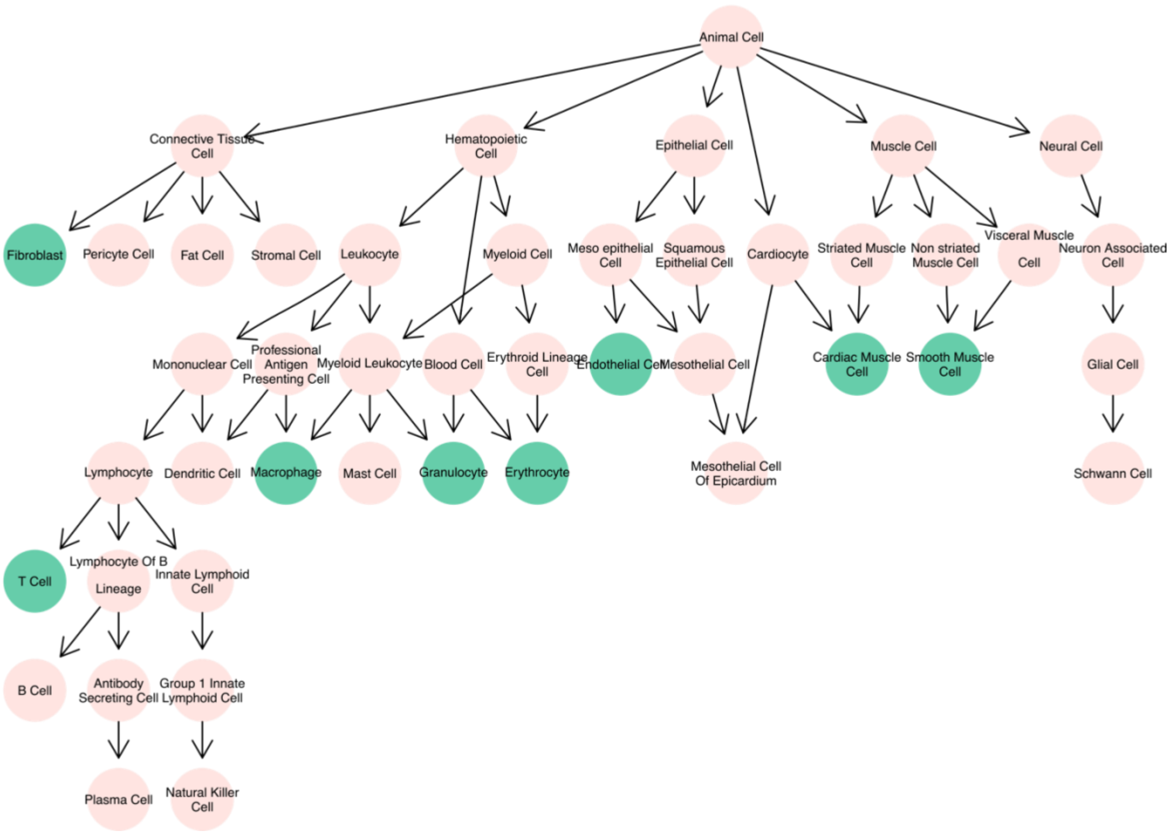

**Extended Data Figure 20-24: scOntoMatch aligned cell type tree in heart task among five species.** Annotated cell types from the heart tissue of the five species are represented as cell type trees whose hierarchy follows Cell Ontology (Osumi-Sutherland et al. 2021). Each circle represents a cell type term in the Cell Ontology system and is coloured by whether the term is present in the dataset. Each arrow starts with a parental term and ends with its children term. scOntoMatch is used to align the ontology annotation across the five species, resulting in comparable annotation granularity. The cell type tree of the original annotation is shown in the top half and the bottom half shows scOntoMatch aligned cell type annotation for each species.

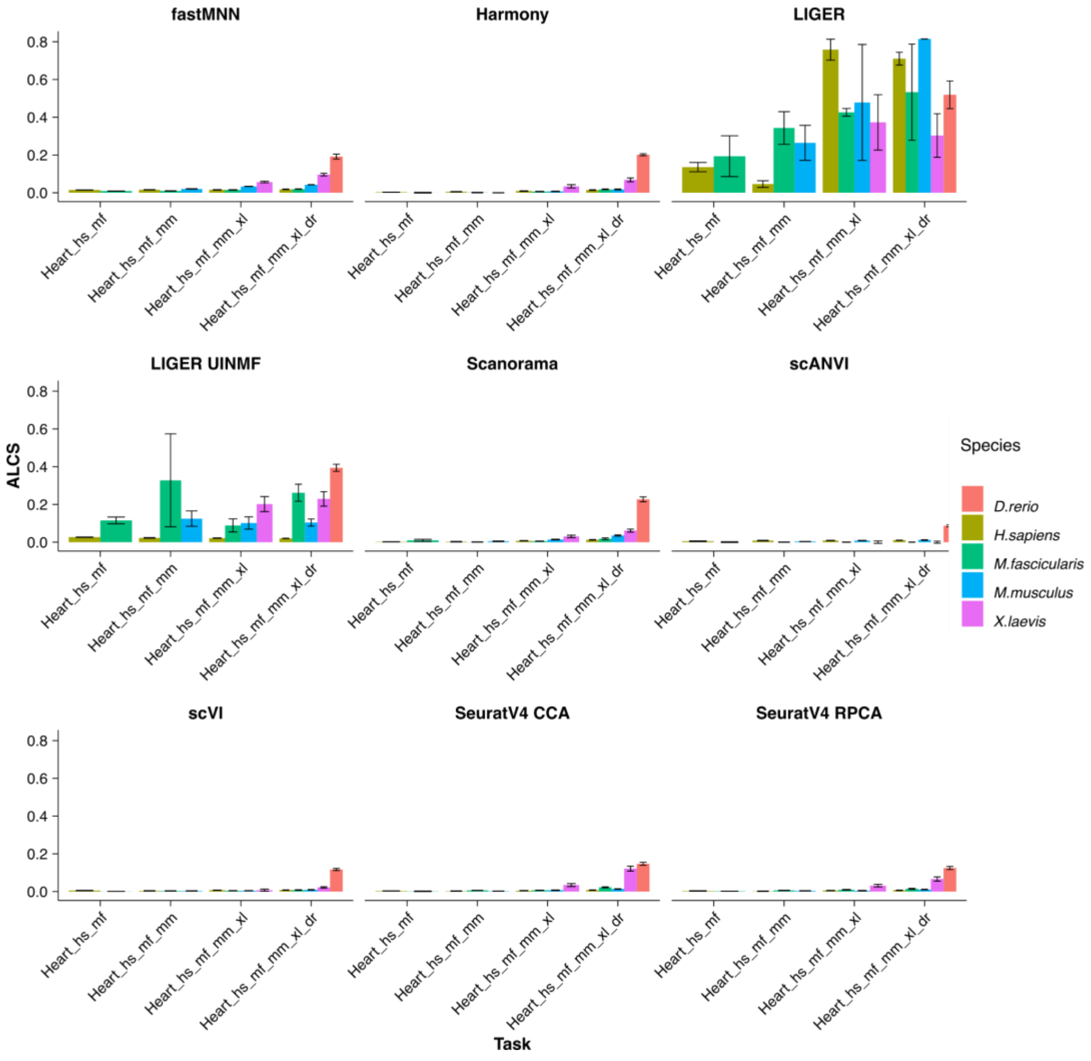

**Extended Data Figure 25: Per-species ALCS in all strategies in heart tasks.** Bars represent the mean ALCS of all homology methods and error bars indicate SD. Lower ALCS suggests less loss of cell type distinguishability after integration. ALCS: accuracy loss of cell type self-projection. hs, *Homo sapiens*, human; mf, *Macaca fascicularis*, long-tailed macaque; mm, *Mus musculus*, mouse; xl, *Xenopus laevis*, african clawed frog; dr, *Danio rerio*, zebrafish.

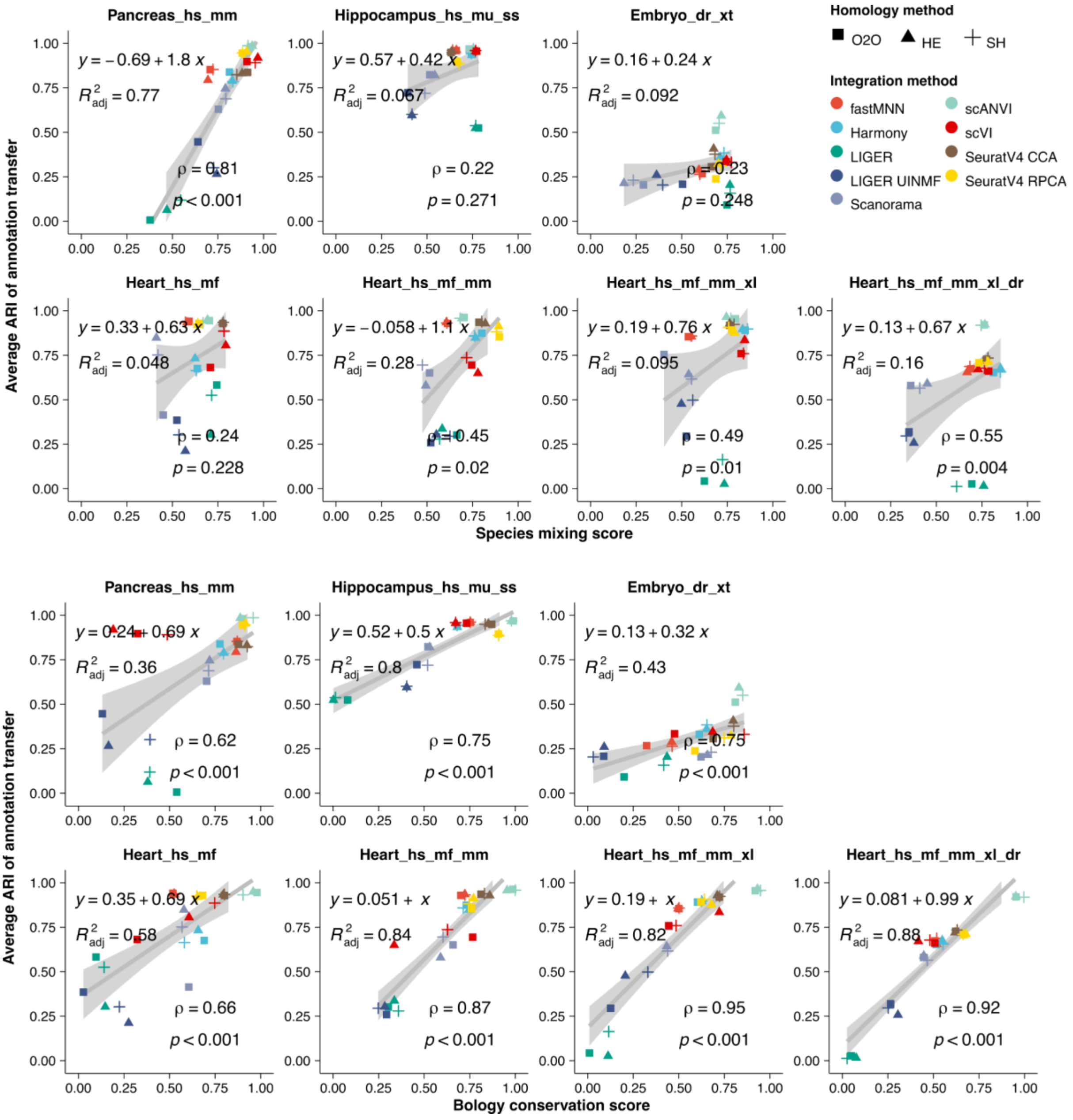

**Extended Data Figure 26: Correlation between ARI and species mixing score or biology conservation score for 7 reference tasks in all strategies.** Showing the average ARI of all pairwise annotation transfers between species in a particular integration output. A high ARI suggests a successful annotation transfer. ARI significantly correlates with the biology conservation score in all tasks (Spearman's rank correlation significance P value $< 0.001$) while the correlation is weaker for the species mixing score. Strategies are represented by points whose colour is the integration algorithm and whose shape is the homology method. ARI, adjusted rand index; ρ, Spearman's rank correlation coefficient; p, P-value of Spearman's rank correlation; $R^2_{adj}$, adjusted goodness-of-fit of the linear model; O2O, only use one-to-one orthologs; HE, one-to-one orthologs plus one-to-many and many-to-many orthologs matched by higher average expression level; SH, one-to-one orthologs plus one-to-many and many-to-many orthologs matched by stronger homology confidence; hs, *Homo sapiens*, human; mf, *Macaca fascicularis*, long-tailed macaque; mu, *Macaca mulatta*, rhesus macaque; mm, *Mus musculus*, mouse; ss, *Sus scrofa*, pig; xl, *Xenopus laevis*, african clawed frog; xt, *Xenopus tropicalis*, western clawed frog; dr, *Danio rerio*, zebrafish.

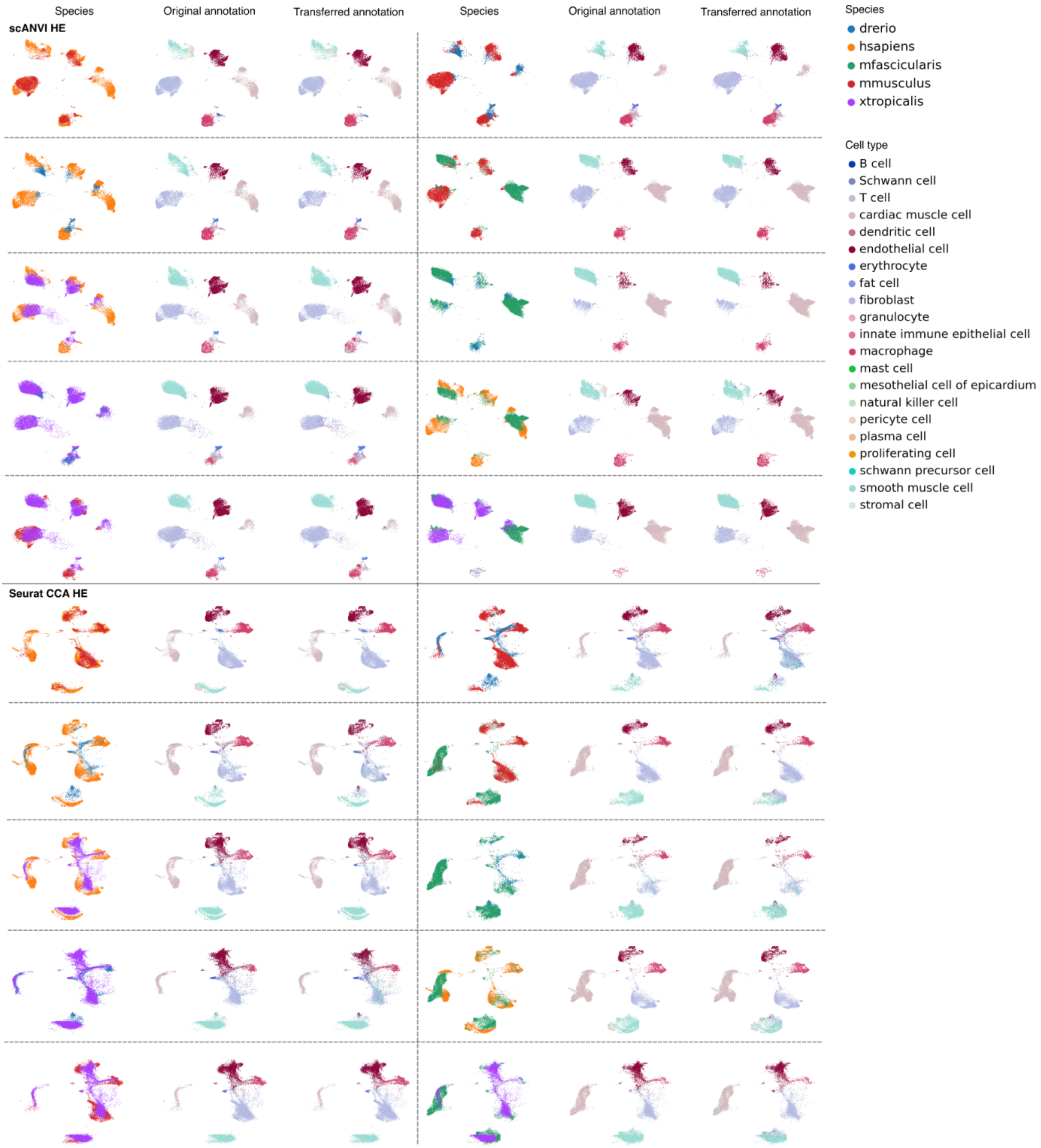
Species
Original annotation
Transferred annotation
Species
Original annotation
Transferred annotation
scANVI HE
Seurat CCA HE
Species
drerio
hsapiens
mfascicularis
mmusculus
xtropicalis
Cell type
B cell
Schwann cell
T cell
cardiac muscle cell
dendritic cell
endothelial cell
erythrocyte
fat cell
fibroblast
granulocyte
innate immune epithelial cell
macrophage
mast cell
mesothelial cell of epicardium
natural killer cell
pericyte cell
plasma cell
proliferating cell
schwann precursor cell
smooth muscle cell
stromal cell

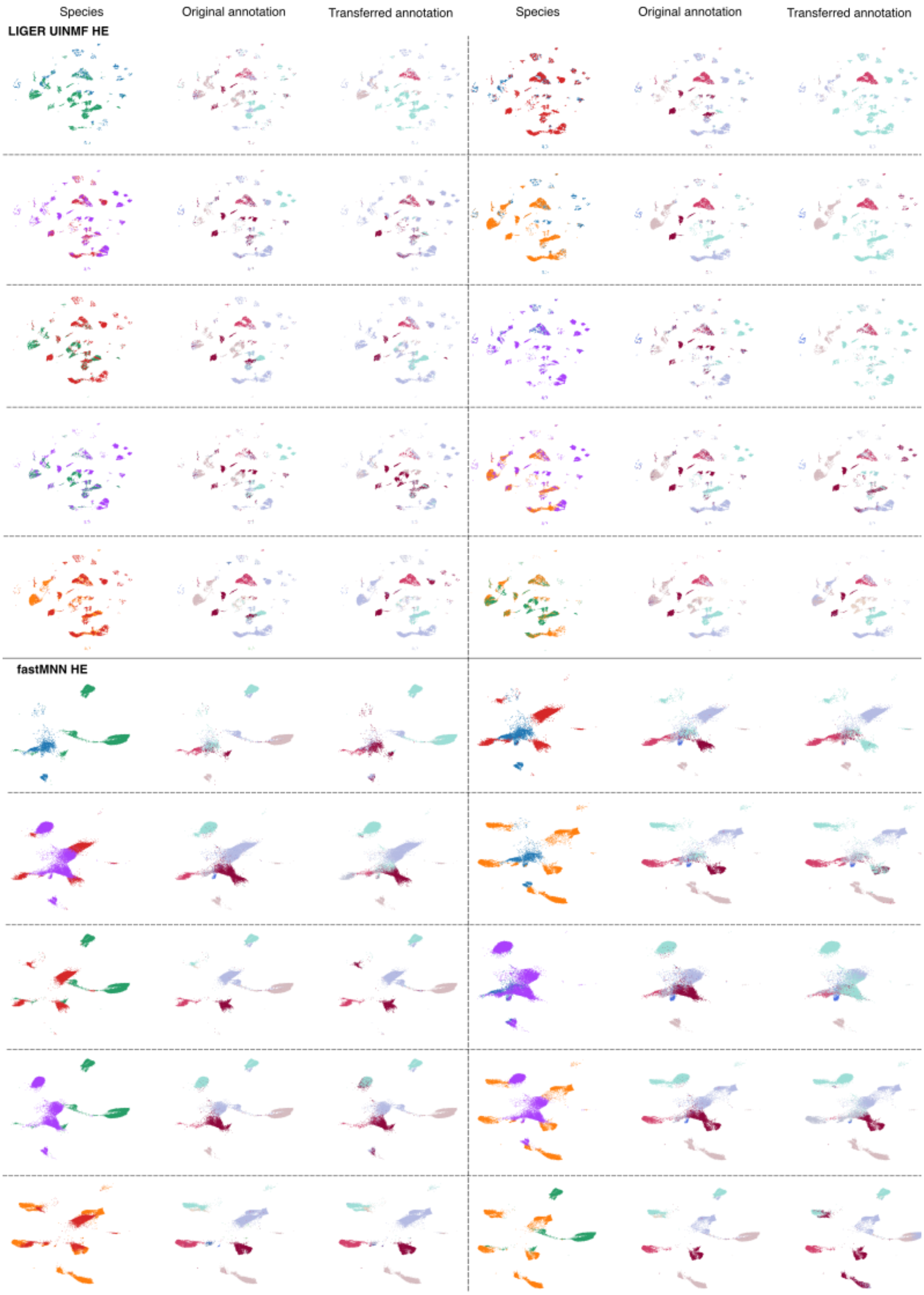


**Extended Data Figure 27: UMAP visualisation of species, original and transferred annotations in the Heart_hs_mf_mm_xl_dr task.** Showing UMAPs from scANVI HE and SeuratV4 CCA HE as examples of a successful annotation transfer. Despite minor confusions between similar cell types, such as epithelial cells and smooth muscle cells, annotation transfer is to a large extent successful even between distant species such as human and zebrafish in these two strategies. On the other hand, results from LIGER UINMF HE and fastMNN HE were unsuccessful, Transferred annotation diverged greatly from the original annotation. HE, one-to-one orthologs plus one-to-many and many-to-many orthologs matched by higher average expression level.

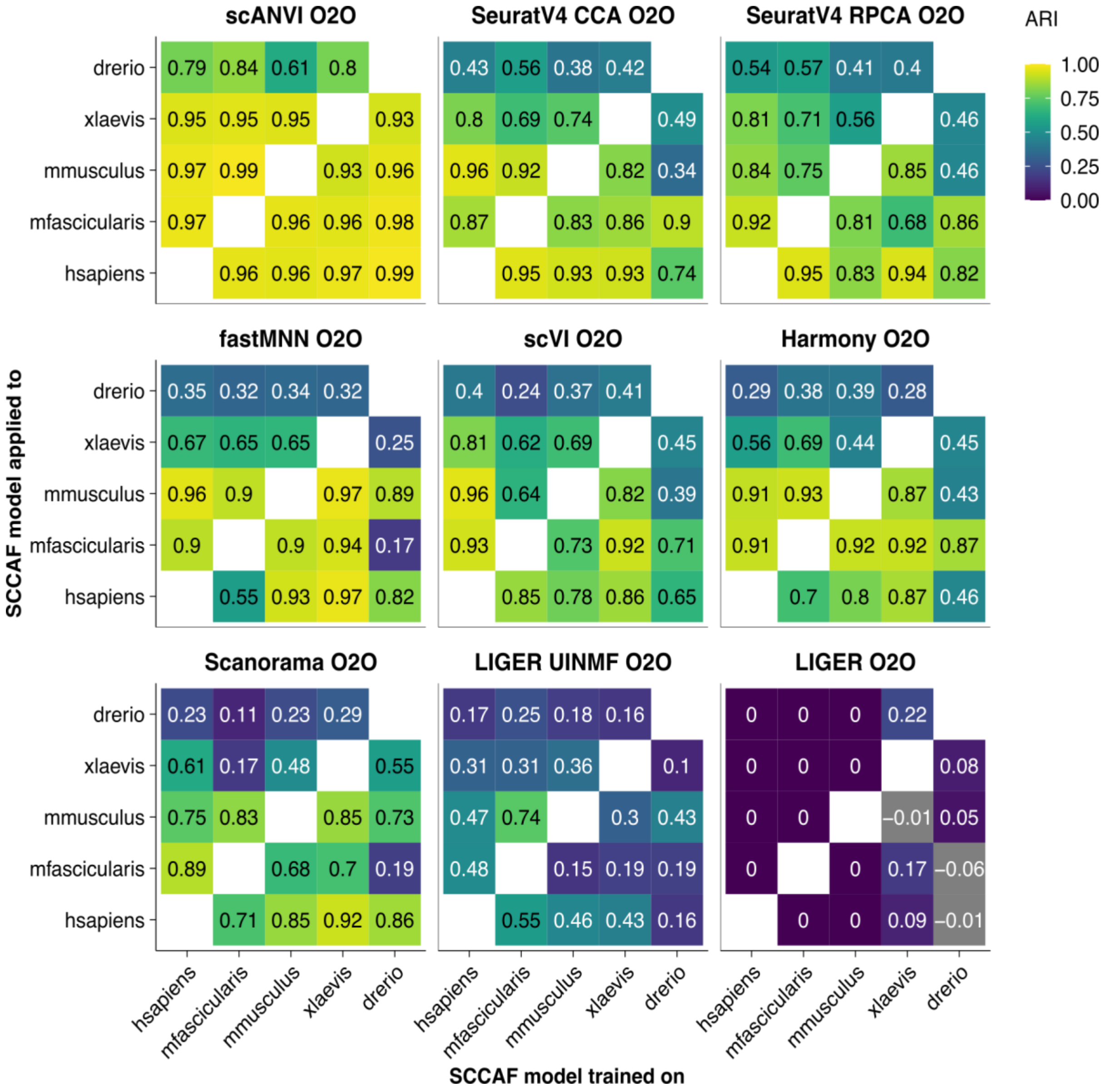


**Extended Data Figure 28: ARI for annotation transfer in one-to-one orthologs integrated data in the Heart_hs_mf_mm_xl_dr task between pairs of species.** Showing the pairwise annotation transfer ARI between all pairs of species. Generally, species that have diverged for a long time have lower ARI, suggesting a harder annotation transfer. O2O, only uses one-to-one orthologs; SCCAF, single cell clustering assessment framework; ARI, adjusted rand index.

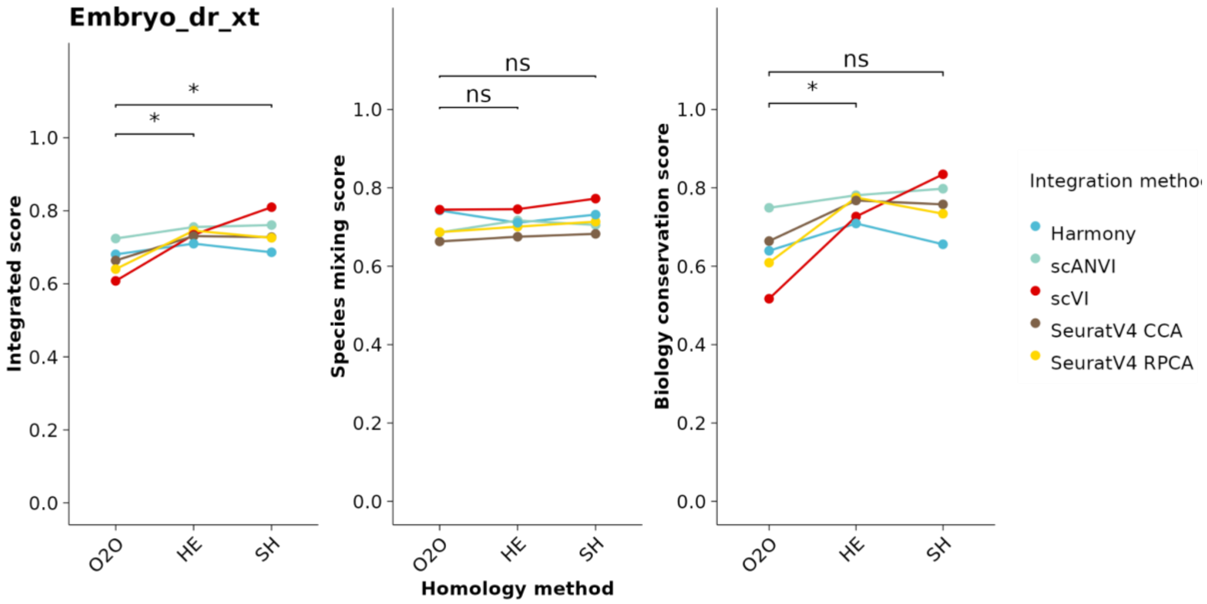


**Extended Data Figure 29: improvement of scores by adding in-paralogs in the Embryo_dr_xt task for successful algorithms.** Algorithms that achieved higher integrated scores than unintegrated data were considered successful. The number represents the P value of the Wilcoxon signed rank test adjusted by the Benjamin-Hochberg procedure.

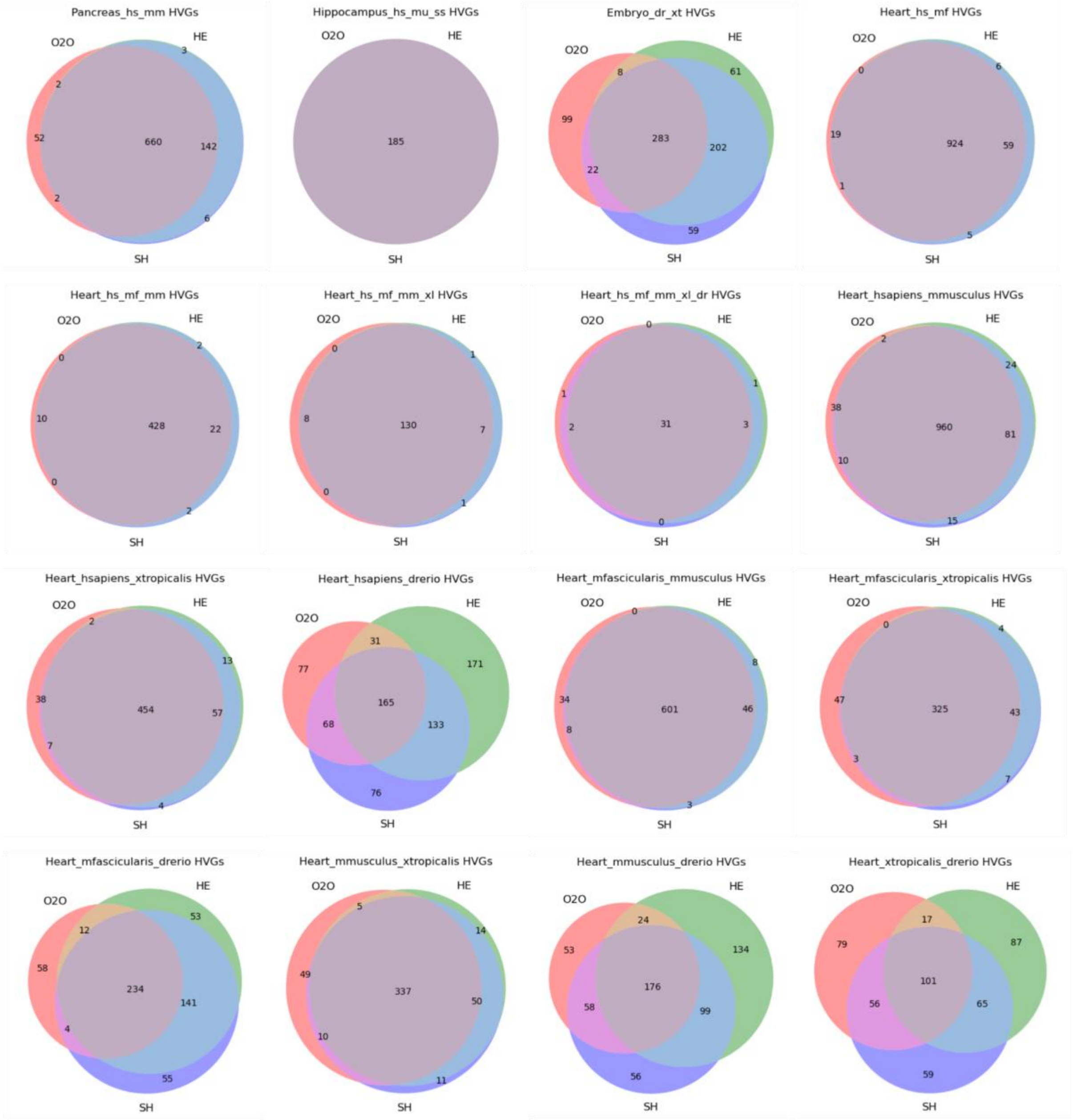


**Extended Data Figure 30 Intersection of HVGs between three types of homology concatenated data in 16 tasks.** HVGs were selected per batch key and the intersection of HVGs from all batches was shown (see Methods for the batch key of each task). Venn diagrams show the intersection of HVGs among three types of homology concatenated data. HVG, highly variable genes; O2O, only use one-to-one orthologs; HE, one-to-one orthologs plus one-to-many and many-to-many orthologs matched by higher average expression level; SH, one-to-one orthologs plus one-to-many and many-to-many orthologs matched by stronger homology confidence.
